# An encyclopedia of enhancer-gene regulatory interactions in the human genome

**DOI:** 10.1101/2023.11.09.563812

**Authors:** Andreas R. Gschwind, Kristy S. Mualim, Alireza Karbalayghareh, Maya U. Sheth, Kushal K. Dey, Evelyn Jagoda, Ramil N. Nurtdinov, Wang Xi, Anthony S. Tan, Hank Jones, X. Rosa Ma, David Yao, Joseph Nasser, Žiga Avsec, Benjamin T. James, Muhammad S. Shamim, Neva C. Durand, Suhas S. P. Rao, Ragini Mahajan, Benjamin R. Doughty, Kalina Andreeva, Jacob C. Ulirsch, Kaili Fan, Elizabeth M. Perez, Tri C. Nguyen, David R. Kelley, Hilary K. Finucane, Jill E. Moore, Zhiping Weng, Manolis Kellis, Michael C. Bassik, Alkes L. Price, Michael A. Beer, Roderic Guigó, John A. Stamatoyannopoulos, Erez Lieberman Aiden, William J. Greenleaf, Christina S. Leslie, Lars M. Steinmetz, Anshul Kundaje, Jesse M. Engreitz

## Abstract

Identifying transcriptional enhancers and their target genes is essential for understanding gene regulation and the impact of human genetic variation on disease^1–6^. Here we create and evaluate a resource of >13 million enhancer-gene regulatory interactions across 352 cell types and tissues, by integrating predictive models, measurements of chromatin state and 3D contacts, and large-scale genetic perturbations generated by the ENCODE Consortium^7^. We first create a systematic benchmarking pipeline to compare predictive models, assembling a dataset of 10,411 element-gene pairs measured in CRISPR perturbation experiments, >30,000 fine-mapped eQTLs, and 569 fine-mapped GWAS variants linked to a likely causal gene. Using this framework, we develop a new predictive model, ENCODE-rE2G, that achieves state-of-the-art performance across multiple prediction tasks, demonstrating a strategy involving iterative perturbations and supervised machine learning to build increasingly accurate predictive models of enhancer regulation. Using the ENCODE-rE2G model, we build an encyclopedia of enhancer-gene regulatory interactions in the human genome, which reveals global properties of enhancer networks, identifies differences in the functions of genes that have more or less complex regulatory landscapes, and improves analyses to link noncoding variants to target genes and cell types for common, complex diseases. By interpreting the model, we find evidence that, beyond enhancer activity and 3D enhancer-promoter contacts, additional features guide enhancer-promoter communication including promoter class and enhancer-enhancer synergy. Altogether, these genome-wide maps of enhancer-gene regulatory interactions, benchmarking software, predictive models, and insights about enhancer function provide a valuable resource for future studies of gene regulation and human genetics.

## Introduction

Noncoding DNA elements called enhancers have essential functions in controlling cell-type specific gene regulation and cellular programs, and likely contain a majority of causal variants for common, complex diseases^4–6^. Yet, it remains an open challenge to accurately identify which elements act as enhancers in a given cell type and link them to the nearby genes that they regulate (hereafter, “enhancer-gene regulatory interactions”)^1–3^.

Toward this goal, the ENCODE Project has conducted thousands of experiments to identify candidate *cis*-regulatory elements and annotate their chromatin state and 3D physical interactions across hundreds of cell types and tissues^7^. Using these and other data, various predictive models have been developed and applied to identify enhancer-gene regulatory interactions^8–16^. Despite recent progress, key challenges remain.

Many previous efforts are missing information on the accuracy of predictions, have not been systematically compared to others using a common set of benchmarks, and/or have known limitations in prediction accuracy^6,8–12,16,17^. We need larger sets of genetic perturbation data and a community framework to evaluate, compare, and develop improved predictive models of enhancer-gene regulatory interactions.

Previous predictive models have identified certain features important for prediction accuracy, including the strength of activating chromatin marks at an element (“enhancer activity”) and the frequency at which an element physically contacts a nearby promoter (“3D contact”)^9^. However, further analysis is needed to evaluate the utility of different experimental assays in estimating these features, to compare the relative importance of these or other molecular features, and to explore how these features might inform molecular mechanisms of enhancer-promoter communication.

Some predictive models are difficult to apply to new cell types because they require analysis of many cell types simultaneously^8,11,12^ or a large number of experimental inputs in any given cell type^10^. As such, we are missing a resource of enhancer-gene regulatory interactions across the hundreds of cell types and tissues profiled by the ENCODE Project that can inform fundamental properties of gene regulation, link noncoding variants to genes, integrate with other ENCODE analysis products, and expand to new cell types in the future.

Data collected in this final phase of the ENCODE Project include CRISPR perturbations of candidate enhancers^9,18–20^, high-resolution Hi-C data^21^, and maps of DNase I hypersensitivity across new cell types^22^ — providing an opportunity to build a common benchmarking framework, build improved predictive models, and apply these models to construct a map of enhancer-gene regulatory interactions in the human genome.

## An encyclopedia of enhancer-gene regulatory interactions

Here we present the ENCODE resource of enhancer-gene regulatory interactions (**Fig. 1**), which includes three components:

1. **A new set of predictive models, collectively termed “ENCODE-rE2G”**. We developed new supervised classifiers that integrate molecular features of chromatin state and 3D physical interactions to predict enhancer-gene regulatory interactions in a given cell type. ENCODE-rE2G involves supervised training directly on CRISPR perturbation data, and enables predicting which enhancers regulate which genes in a given cell type based on various combinations of cell-type specific experimental data (minimally, DNase-seq combined with a reference ENCODE Hi-C map averaged across many tissues).
2. **A new benchmarking framework to evaluate predictive models.** We collected and harmonized genetic perturbation data relevant to enhancer-gene regulatory interactions, including CRISPR perturbation experiments, expression quantitative trait loci (eQTLs), and variants from genome-wide association studies (GWAS). We developed a pipeline to quantify and annotate the accuracy of predictive models, and applied this to benchmark ENCODE-rE2G and hundreds of alternative models at key prediction tasks. An important difference between this and some previous benchmarking analyses is that we use as gold standards only data describing *regulatory interactions*, derived from genetic perturbations, as opposed to *physical interactions*, such as those measured by Hi-C or ChIA-PET. These perturbations datasets, analysis pipelines, and benchmarking results will facilitate studies to develop improved models of enhancer-gene regulatory interactions.
3. **An encyclopedia of enhancer-gene regulatory interactions in the human genome**. We applied the most general of the ENCODE-rE2G models across 352 ENCODE cell types, tissues, and contexts (collectively, “biosamples”), and identified 13,455,443 regulatory interactions between distal elements and target genes. This atlas includes an average of 23,846 unique regulatory elements and 62,509 element-gene regulatory interactions per biosample, and in total annotates 0.76% of the approximately 3 billion basepairs in the human genome. This resource enables looking up a gene to find predicted enhancers, looking up an element to find predicted target genes, and identifying predicted cell types and target genes for noncoding variants.

**Figure 1.**
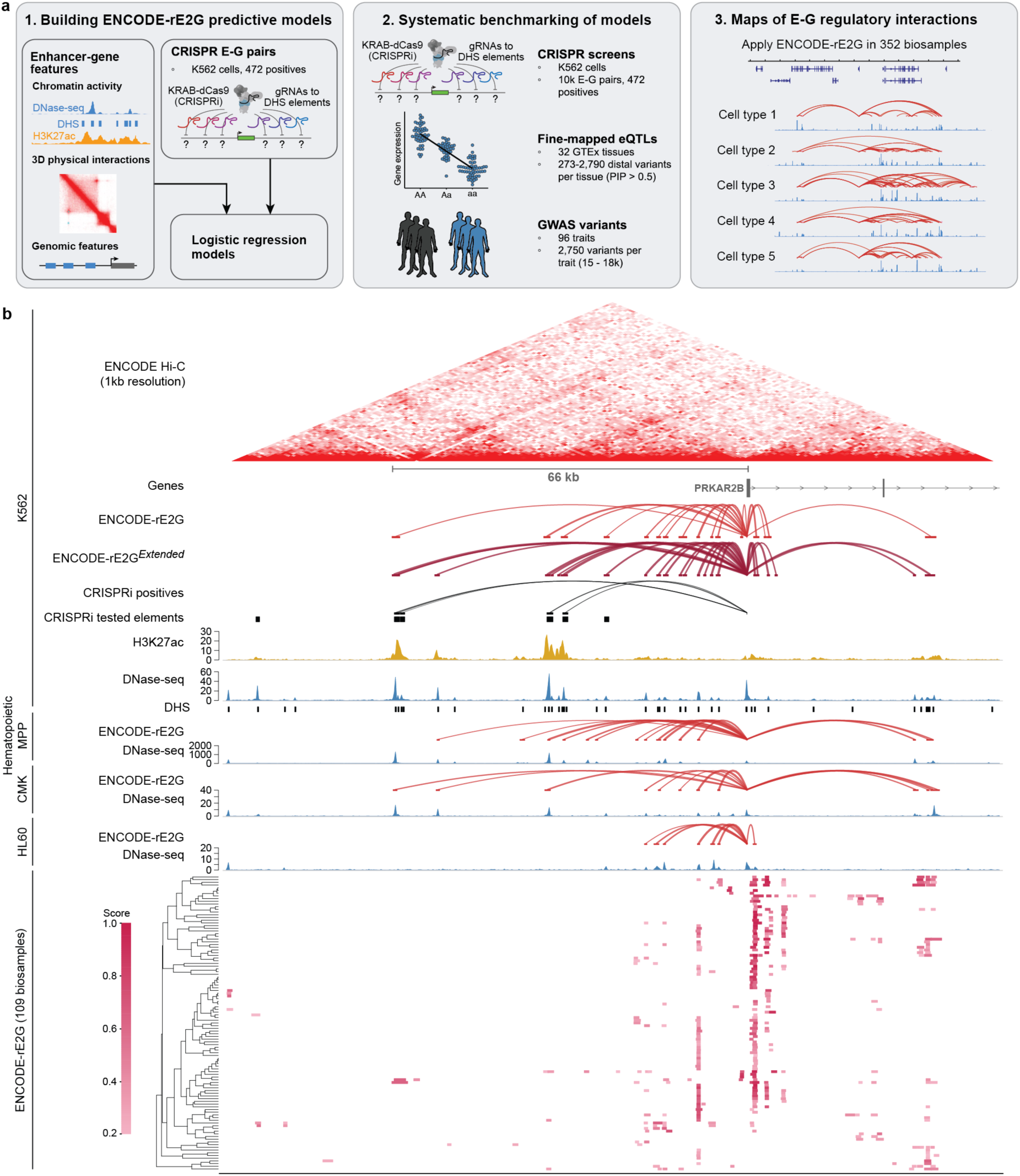
The ENCODE resource of enhancer-gene regulatory interactions. **a**, Project overview: Using CRISPR data in K562 in combination with molecular features for enhancer activity, 3D physical interaction and genomic landscape, we trained new semi-supervised predictive models on the CRISPRi data, called ENCODE-rE2G, to predict enhancer-gene regulatory connections from molecular data. We benchmarked their performance using systematic benchmarking against ground truth datasets from CRISPRi, eQTL and GWAS data and compared ENCODE-rE2G’s performance against other published models. Using the ENCODE-rE2G model we built genome-wide maps of E-G regulatory connections across 352 ENCODE biosamples. **b,** Enhancer-gene regulatory connections for the *PRKAR2B* locus, showing detailed data and predictions in K562, DNase-seq and ENCODE-rE2G enhancer-gene connections in three related cell types, and ENCODE-rE2G scores for predicted *PRKAR2B* enhancers in 92 biosamples with at least three enhancers.

Below we describe the creation, validation, and application of this resource to chart global properties of enhancer-gene regulatory interactions, annotate variants associated with complex traits, and identify new molecular features that tune enhancer-promoter communication.

## Supervised classification of enhancer-gene regulatory interactions

Previous predictive models of enhancer-gene regulatory interactions have largely involved unsupervised approaches based on enhancer-promoter correlations^12–14^, molecular rules^9,16^, or supervised learning on proxies such as 3D loops or gene expression^8,10,11,15^. We reasoned that, by combining epigenomic data with recent large-scale enhancer-targeting CRISPR perturbation datasets, we could build an improved model by training a supervised classifier directly on CRISPR data in a single cell type, and then apply the trained model across the genome and to new cell types. A similar approach was pioneered in previous work^23^, but using a much smaller CRISPR perturbation dataset.

We constructed two logistic regression classifiers to predict CRISPR-validated element-gene pairs using different combinations of molecular features (**Tables S1, S3**). In the “ENCODE-rE2G” model, we used 13 features that could be computed using only cell-type specific DNase-seq data, thereby facilitating analyses across many hundreds of ENCODE biosamples. Specifically, we included features related to (i) chromatin state (quantitative DNase-seq signals at the element and promoter), (ii) 3D contact frequency (averaged from ENCODE Hi-C in 35 diploid biosamples to capture cell-type-invariant features of genome organization); (iii) the Activity-by-Contact (ABC) model^9^, which multiplies enhancer activity and 3D contact frequency; (iv) genomic position (*e.g.*, distance from element to promoter); (v) promoter class (*e.g.*, whether the gene is uniformly and ubiquitously expressed across cell types (*i.e.*, a “housekeeping” gene)); and (vi) information about nearby enhancers (*e.g.*, the activity of all other elements within 5 kb of the perturbed element) (**Table S3**). In the ENCODE-rE2G*^Extended^* model, we sought to evaluate the utility of additional assays available in Tier 1 ENCODE cell types, and expanded to include a total of 47 features derived from DNase-seq, histone ChIP-seq, cell-type specific ENCODE Hi-C, and ChIA-PET experiments, as well as features from additional predictive models such as EpiMap^8^ and GraphReg^10^ that jointly analyze many datasets (**Table S3**).

To train and evaluate models, we aggregated a gold-standard dataset of 10,411 element-gene pairs tested with CRISPR in K562 erythroleukemia cells, an ENCODE Tier 1 cell line. We re-analyzed and harmonized data from previous studies that used genetic perturbations (mostly CRISPR interference (CRISPRi)) to inhibit candidate enhancers and measure effects on gene expression^9,19,23–25^ (see **Note S1**). Importantly, we developed approaches to compute statistical power for every tested element-gene pair, identifying 472 “positive” unique element-gene pairs where CRISPR perturbation of the element led to a significant decrease in gene expression (–1 to –93% effects, **Fig. 1c, Fig. S1.1f**) and 9,938 “negative” element-gene pairs where no significant reduction in expression was observed despite the experiment having good power to detect >15-25% effects on gene expression (**Note S1**). We trained logistic regression classifiers to distinguish positives from negatives using hold-one-chromosome-out cross-validation. Then, we applied the trained model to all pairs of element-gene pairs across the genome and to new cell types.

Each element-gene pair is annotated with a score, corresponding to the probability of a regulatory effect from the logistic regression classifier. To binarize the predictions, we selected a threshold on the score that achieved 70% recall in the CRISPR training dataset, as in previous studies^9^. At this threshold, ENCODE-rE2G identified an average of 38,225 E-G regulatory interactions per biosample (36,822 for ENCODE-rE2G*^Extended^*), 23,847 predicted “enhancers” (distal elements predicted to regulate at least one gene; 21,325 for ENCODE-rE2G*^Extended^*), and included well-studied enhancers such at the *HBE1*, *GATA1* and *MYC* loci (**Fig. S10**).

## Systematic benchmarking of ENCODE-rE2G

To quantify the accuracy of enhancer-gene regulatory interaction predictions, we developed a systematic benchmarking pipeline by aggregating genetic perturbation data from CRISPR enhancer perturbations and fine-mapped eQTL and GWAS variants (**Fig 2a**). We used this pipeline to evaluate ENCODE-rE2G and 574 other models (**Table S1**), including (i) the ABC model, which we re-computed here natively in hg38 using various combinations of ENCODE assays to estimate enhancer activity and enhancer-promoter 3D contact; (ii) previous machine learning models that infer enhancer-gene regulation, including EpiMap^8^, GraphReg^10^, CIA^16^, Enformer^11^, and EPIraction^26^; and (iii) simple baselines such as linking elements to the closest expressed transcription start site (TSS) or gene, linking elements to genes solely on the basis of loops or domains from 3D contact measurements, or correlating distal element and promoter accessibility across cell types.

**Figure 2.**
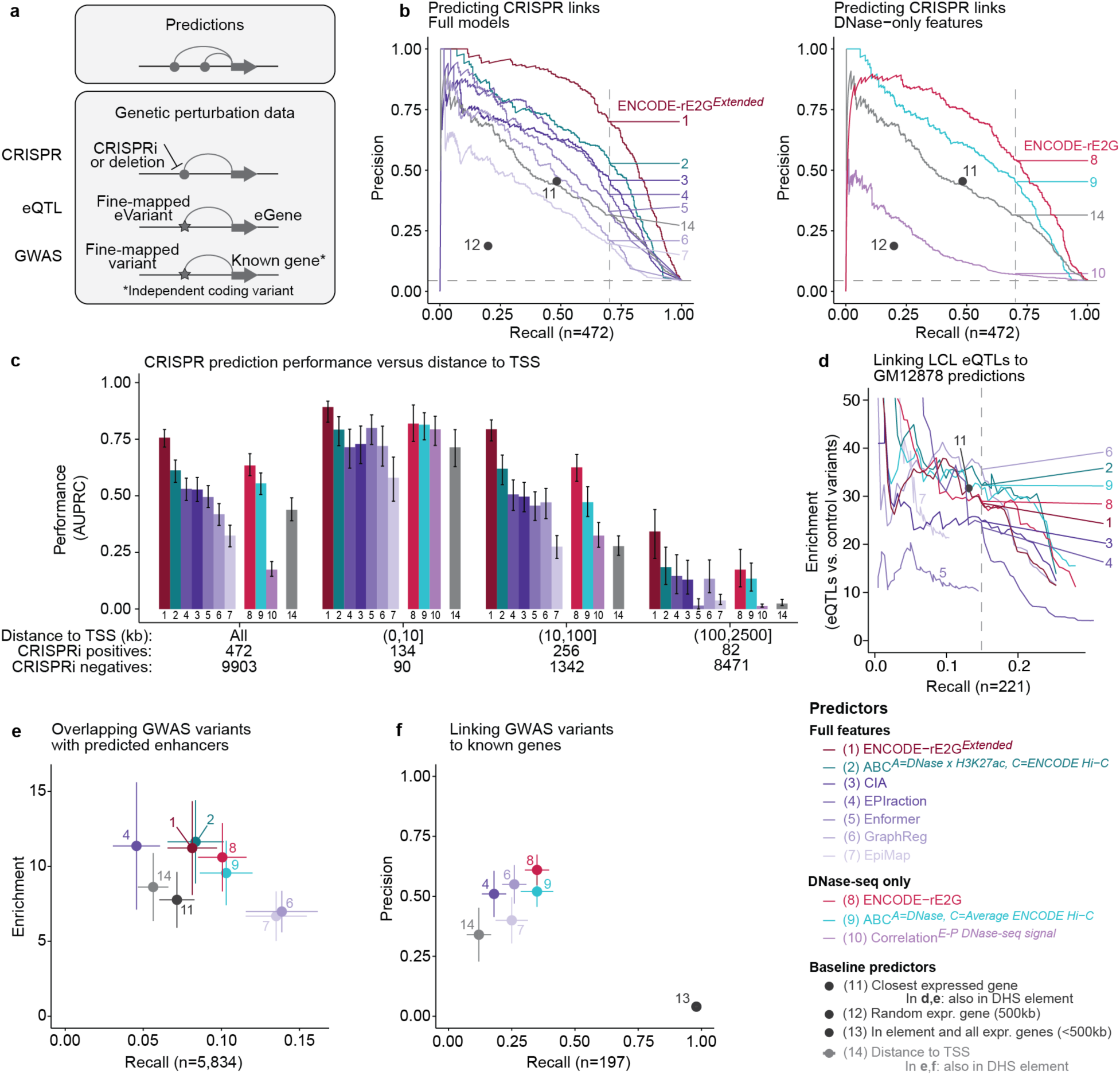
Benchmarking predictions of enhancer-gene regulatory interactions. **a**, We benchmarked the performance of predictive models for enhancer – gene regulatory connections on three different prediction tasks: 1) Linking enhancers to CRISPR-validated target genes, 2) Enrichment of putative regulatory variants from fine-mapped eQTL and GWAS datasets, and 3) Linking variants to putative causal target genes from fine-mapped GWAS datasets. **b,** Precision-Recall curves showing the performance of predictive models at predicting experimental results of CRISPRi data in K562 cells. Combined CRISPRi data was assembled by combining element-gene pairs from the re-analyzed Nasser *et al.*, 2021^6^, Gasperini *et al.*, 2019^24^ and Schraivogel *et al.*, 2020^23^ datasets (10,375 tested element-gene pairs, 472 positives). Curves represent continuous predictors with the dashed vertical line indicating performance at a threshold corresponding to 70% recall. Single dots represent performance of binary predictors. **c,** Performance (AUPRC) of quantitative predictors as a function of distance to TSS. Color legend from panel d applies. Error bars represent 95% range of AUPRC values inferred via bootstrap (1000 iterations). **d,** Enrichment – recall curves showing the ratio of fine-mapped distal noncoding eQTLs with a PIP>0.5 in EBV-transformed lymphocytes in predicted enhancers compared to distal noncoding common variants (enrichment, y-axis) versus fraction of variants overlapping enhancers linked to the correct gene in GM12878 cells (recall, x-axis) across different score thresholds for enhancers predicted by different predictive models. Numerical results are reported in **Table S15**. **e,** Average enrichment and recall of GWAS fine-mapped SNPs (PIP > 0.1) in predicted elements for each element-gene linking strategy for 5 “likely relevant” pairs cell lines and biomarker traits: 3 RBC-related (RBC count, Mean corpuscular volume, HbA1c) traits for K562 cell line and 2 Lymphocyte-related (Lymphocyte count, Autoimmune disease-combined) traits for the GM12878 cell line. Error bars denote 95% confidence intervals. **f,** For 197 non-coding credible sets corresponding to 11 blood-related traits that are linked to exactly one “putatively causal” gene with a coding fine-mapped variant within 2Mb on either side of the lead variant, we compute the precision and recall in linking it to this gene for each of the element-gene predictions in blood biosamples (see Methods). Error bars represent 95% confidence intervals. Numerical results are reported in **Table S13**.

We found that ENCODE-rE2G and ENCODE-rE2G*^Extended^* achieved state-of-the-art performance across a variety of prediction tasks (**Fig. 2b-f**):

In the CRISPR enhancer perturbation dataset used to train ENCODE-rE2G, we compared predictors to the experimental results by means of precision-recall plots (**Fig. 2b**), computing area under the precision-recall curve (AUPRC) (**Fig. 2c**), precision at a fixed recall of 70% (**Fig. S8c**), and the correlation between the predictive scores and the measured quantitative effect on expression (**Extended Data** Fig. 2f**, Fig. S9**), using hold-one-chromosome-out cross-validation for supervised models. Between the two models that used cell-type specific DNase only (ENCODE-rE2G and ABC*^A=DNase, C=Average^ ^Hi-C^*), ENCODE-rE2G performed significantly better by all measures (AUPRC = 0.63 vs 0.55, *P_bootstrap_* = 0.0001; precision at 70% recall = 54% vs 46%, *P_bootstrap_* = 0.0001; and Pearson’s *R* = –0.44 vs –0.37, *P* = 7.16 x 10^−9^; **Fig. 2b, Fig. S8c, Extended Data** Fig. 2f), including at different distance thresholds (**Fig. 2c**). Similarly, among the 7 models that use an expanded feature set, ENCODE-rE2G*^Extended^* outperformed all others, including the second-best model, ABC^Activity=DHS*H3K27ac, Contact=ENCODE Hi-C^ (AUPRC = 0.76 vs 0.61, *P_bootstrap_* = 0.0001; precision at 70% recall = 70% vs 54%, *P_bootstrap_* = 0.0001; and Pearson’s *R* = –0.49 vs – 0.43, *P* = 5.58 x 10^−8^; **Fig 2b,c, 2b, Fig. S8c, Extended Data** Fig. 2f). Other commonly used baseline methods — such as assigning each element to the closest expressed gene, correlating DNase-seq signal at elements and nearby promoters across cell types, or assigning elements to target genes solely based on features of 3D contacts — performed less well, as did other predictive models (**Fig. 2b,c**). We describe detailed evaluations of these other predictors and baseline methods in **Note S2, Note S3, Note S5,** and **Note S7**.

We next assessed the ability of ENCODE-rE2G models to transfer to new cell types to identify eQTL variants and their target genes. We first compared predictions in GM12878 to fine-mapped eQTLs from lymphoblastoid cell lines in GTEx^27^, focusing on distal noncoding eQTL variants with fine-mapping posterior probability of inclusion (PIP) > 50% (n=273). We computed the recall at identifying eQTL variant-gene links (fraction of eQTL variants that are contained in an element and predicted to regulate the correct eQTL gene) as well as enrichment of eQTL variants in predicted enhancers (with respect to all distal noncoding variants) at different thresholds on the predictor score. ENCODE-rE2G*^Extended^*and ENCODE-rE2G perform comparably to other well-performing models (both with enrichment = 28 at a recall of 15%) (**Fig. 2d, Table S15**). We also examined fine-mapped eQTL variant-gene links from 11 other GTEx tissues that were represented in ENCODE biosamples (**Table S14**). Using the score threshold corresponding to 70% in the K562 CRISPR dataset, ENCODE-rE2G achieved an average enrichment of 6.5 across tissues (range: 3.5-11.4), similar than other predictive and baseline models, while achieving a stronger recall, identifying an average of 16% (range: 10–22%) of fine-mapped eQTL variants and linking them to the correct eGene (**Extended Data** Fig. 3c,d).

Finally, we tested the ability of ENCODE-rE2G and the other predictive models to identify GWAS variants and link them to known genes, using 29.2K distal non-coding variants for 94 traits from the UK Biobank previously fine-mapped using SuSIE^28^ (PIP > 0.1). We first examined whether predicted enhancers from the K562 erythroid and GM12878 lymphoblastoid cell lines were enriched for the 5,834 fine-mapped GWAS variants for traits related to red blood cells (red blood cell count, mean corpuscular volume, hemoglobin traits) or lymphocytes (lymphocyte count and All Autoimmune Disease), respectively. ENCODE-rE2G showed the highest significant enrichment (10.6-fold; p-value = 1 x 10^−4^); GraphReg showed the highest recall (38% higher than ENCODE-rE2G) but 65% lower enrichment compared to ENCODE-rE2G (**Fig. 2e**). We next examined the noncoding credible sets for all 11 blood-related traits in which a nearby gene had strong independent evidence of association to the trait via analysis of common coding variation^28^ (197 out of 3529 non-coding credible sets). The ENCODE-rE2G model achieved the highest precision in predicting the correct target gene (61%), significantly higher than the second-best method, ABC^A=DNase, C=Avg. ENCODE^ ^Hi-C^ (52%, p-value of difference = 8 x 10^−4^), with similar recall (35%) for both methods (**Fig. 2f**). These conclusions were supported by additional benchmarking analyses (**Extended Data** Fig. 4).

Altogether, these results suggest that the ENCODE-rE2G models generalize well to new cell types and achieve state-of-the-art performance for a variety of tasks designed to evaluate enhancer-gene regulatory interactions. Based on these analyses, the ENCODE-rE2G resource annotates each predicted regulatory link with the score predicted from the logistic regression model, indicating the probability that the element has a substantial regulatory effect on the target gene. The pre-thresholded set of links can be compared to the results in Figure 2 to estimate the performance of the model at predicting the results of CRISPR perturbations, eQTL variant-gene links, and causal GWAS genes in a given locus. We share the full benchmarking pipeline to enable future refinement and comparison of predictive models (see Code Availability).

## Genome-wide maps of regulatory interactions

We next applied ENCODE-rE2G across 352 biosamples with DNase-seq data generated by the ENCODE Project, and identified a total of 13,455,443 element-gene regulatory interactions using the threshold at 70% recall in the CRISPR benchmark **(Fig. 3a, Table S12).** Properties of these predicted regulatory interactions inform the architecture of enhancer-gene regulation in the human genome:

**Figure 3.**
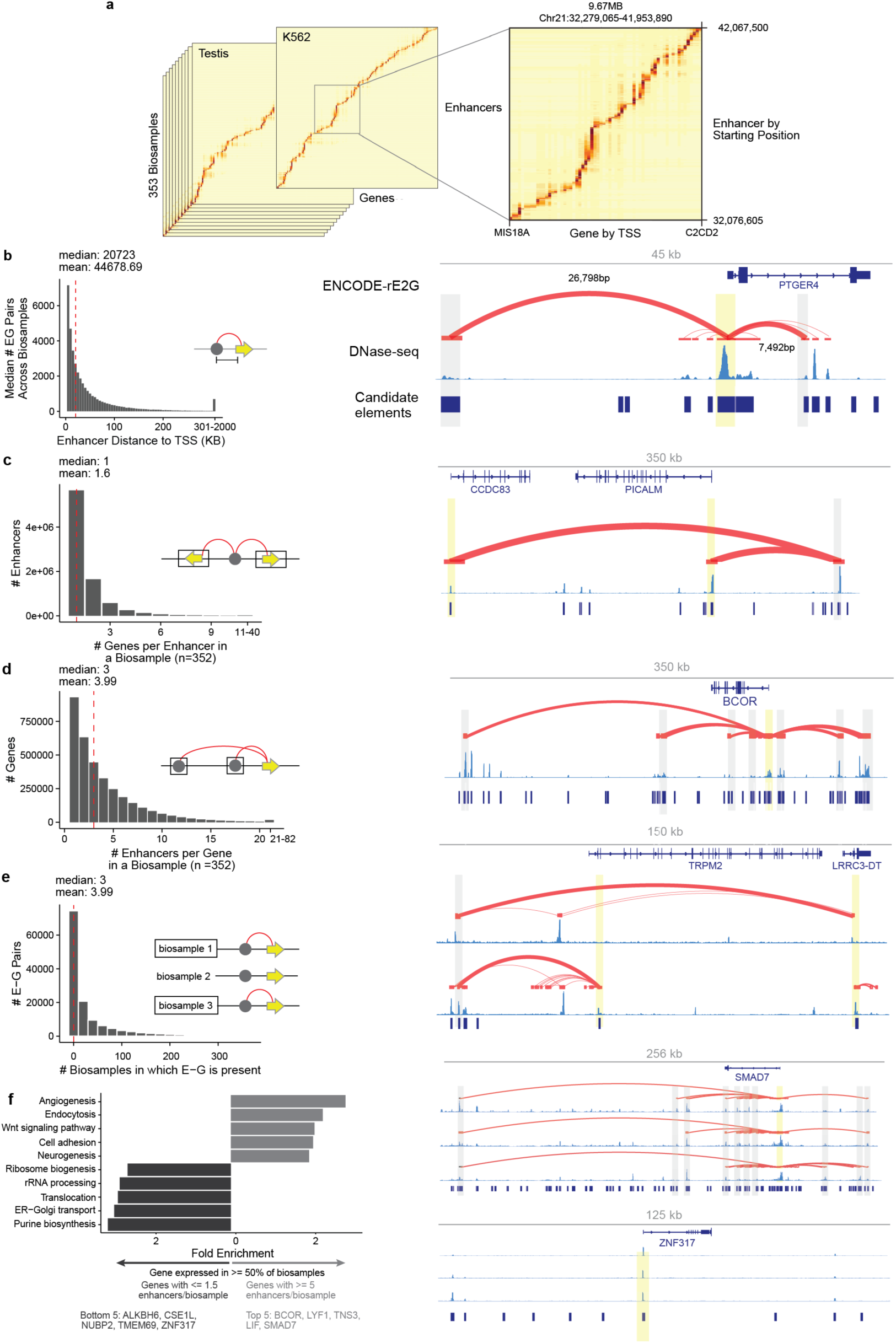
Properties of predicted enhancer-gene regulatory interactions. **a**. The overall scale of the E-G maps is shown as all enhancers x all genes x 352 biosamples. The heatmaps on the left show all ENCODE-rE2G scores for every enhancer gene pair on chr21 of K562 and Testis. The right plot shows a portion of this plot, displaying a 9.67MB region on chr21. For all heatmaps the lightest yellow represents an ENCODE-rE2G score of 0 and the darkest brown represents a score of 0.994. **b**. Distance between an enhancer and the gene it regulates across each of the 352 biosamples. Each bar represents the median number of E-G pairs at that distance bin across all 352 biosamples. The locus plot displays chr5:40,650,000-40,695,000 shows 2 enhancers for the gene *PTGER4* CD4-positive, alpha-beta T cells. **c**. Distribution of the average number of genes regulated by a given enhancer according to the ENCODE-rE2G model across each of the 352 biosamples. Summary statistics available in **Table S18.** The locus plot displays the region from chr11:85,800,000-86,150,000 and shows an enhancer that regulates two genes in pulmonary artery endothelial cells. **d**. Distribution of the average number of enhancers regulating a given gene according to the ENCODE-rE2G model across each of the 352 biosamples. Summary statistics available in **Table S19**. The locus plot shows the region on chrX:39,850,000 – 40,200,000 in which there are 7 enhancers for the *BCOR* gene in CD8-positive alpha-beta T cells. **e**. Distribution of the number of biosamples in which a given enhancer-gene regulatory interaction is detected in the ENCODE-rE2G model. The locus plot shows the region on chr21:44,300,000-44,556,000 in which an enhancer at chr21:44296889-44298071 regulates *LRRC3-DT* in brain microvascular endothelial cells (upper track) and *TRPM2* in right ventricle myocardium superior cells (lower track). **f**. Top 5 functional enrichment of genes expressed across more than half of biosamples stratified by the average number of enhancers they have across biosamples. Biological processes that are enriched in genes with an average of >= 5 enhancers per biosamples are shown in blue and those enriched in genes with an average of <= 1.5 enhancers per biosample are shown in red. All enrichments reported have an adjusted p-value < 0.05. Genes listed are those with the top 5 most enhancers/biosample and top 5 fewest enhancers/biosample. Fold enrichment is calculated relative to the whole genome background set. The locus plots show enhancers for *SMAD7* which averages 14.7 enhancers per biosample (plot range chr18:48780700-49036700) and *ZNF317* which averages 0.25 enhancers/biosample (plot range chr19:9,075,000-9,200,000).

While enhancers can be located up to millions of basepairs from their target genes^25^, the vast majority of enhancer-gene regulatory interactions predicted by ENCODE-rE2G occur over much shorter distances (*e.g.*, a median of 31.8% at <10 kb and 86.8% at <100 kb across biosamples) (**Fig. 3b**). This is consistent with the distance distributions of genetic perturbations in CRISPR experiments, eQTLs, and GWAS positive controls (**Fig. S2.1a**).

Many ENCODE-rE2G enhancers show elevated levels of active chromatin marks, such as H3K27ac and DNase-seq (mean DNase-seq signal for candidate elements across biosamples = 3.27; mean for enhancers linked to at least one gene = 7.52) — but importantly, many do not (15.6% of enhancers have DNase-seq scores below the global median) (**Extended Data** Fig 5h). This may be because elements with apparently weak activity can still have regulatory effects if they are sufficiently close to their target promoter (**Fig. S2.1d**). In part for this reason, annotations of enhancers based on 1D chromatin state alone miss 33% (chromHMM) or 61% (cCRE Catalog) of ENCODE-rE2G enhancers, as well as 30% or 12%, respectively, of the validated enhancer-gene pairs from the Nasser *et al.*, 2021 unbiased CRISPR DHS tiling dataset^6^ (**Extended Data** Fig. 6).

On average, in any given cell type, each gene had 3.99 predicted regulatory elements (median = 3) (**Fig. 3d**), and each element regulated 1.6 genes (median = 1) (**Fig. 3c**). As expected, many element-gene regulatory interactions were highly cell-type specific: 26.4% were present in only one biosample (mean = 34.49, median = 7 biosamples) (**Fig. 3e**), and predicted regulatory interactions were much more specific than the activity of either the promoters or enhancers alone (**Extended Data** Fig 5g,h). This degree of biosample-specificity included many cases where an enhancer regulates different genes in different biosamples (64% of enhancers that were predicted to regulate more than 1 gene were linked to genes in mutually exclusive sets of biosamples, **Fig. 3e**).

Global maps of enhancer-gene regulation can inform the functions of genes and identify those with tight transcriptional control^6,29^. To explore this, we stratified genes based on the number of predicted ENCODE-rE2G enhancers per biosample. Among genes expressed in at least 50% of biosamples, genes with many regulatory elements per cell types (mean >=5 enhancers; 5.9% of genes) were enriched, relative to the whole genome, for being involved in cell-type specific biological processes such as Angiogenesis (2.87-fold enrichment, *P* = 3.60 x 10^−4^), Endocytosis (2.30-fold enrichment, *P* = 0.029), and Wnt signaling pathway (2.10-fold enrichment, *P =* 0.015) (**Fig. 3f**), whereas genes with fewer regulatory elements per cell type (mean <=1.5; 15.7% of genes) were enriched for housekeeping genes with functional enrichments including Purine biosynthesis (3.08-fold enrichment, *P* = 0.04), Golgi transport (2.92 Fold Enrichment, *P* = 1.44 x 10^−14^) and rRNA processing (2.59-fold enrichment, *P* = 4.08 x 10^−5^) (**Fig. 3f**).

## Linking noncoding variants to target genes and cell types

ENCODE-rE2G regulatory links can be integrated with other ENCODE resources and external data to interpret the functions of noncoding variants associated with common disease. To illustrate this, we annotated fine-mapped GWAS variants from 94 UK Biobank traits with ENCODE-rE2G predictions across 352 biosamples describing links from distal enhancers to target genes and predictions from the Polygenic Priority Score (PoPS)^28^ about which genes and pathways are likely to be important for a disease (**Fig. 4a**). As previously observed, fine-mapped variants underlying enhancers were often predicted to regulate multiple genes (average=3.4, max=56), including 62% that are linked to different genes in different cell types; this appears to be a fundamental property of enhancer-promoter regulation^6,9,18^, and motivates combining locus-specific variant-to-gene information with orthogonal information about gene function^28,30^(**Note S7, Fig S7.1**). Accordingly, we developed and evaluated different combinations of ENCODE-rE2G predictions with the polygenic priority score (PoPS)^28^, and found that intersecting top two genes with the strongest ENCODE-rE2G predictions in a given GWAS locus with top two genes with the highest PoPS scores performed well at identifying known genes.

**Figure 4.**
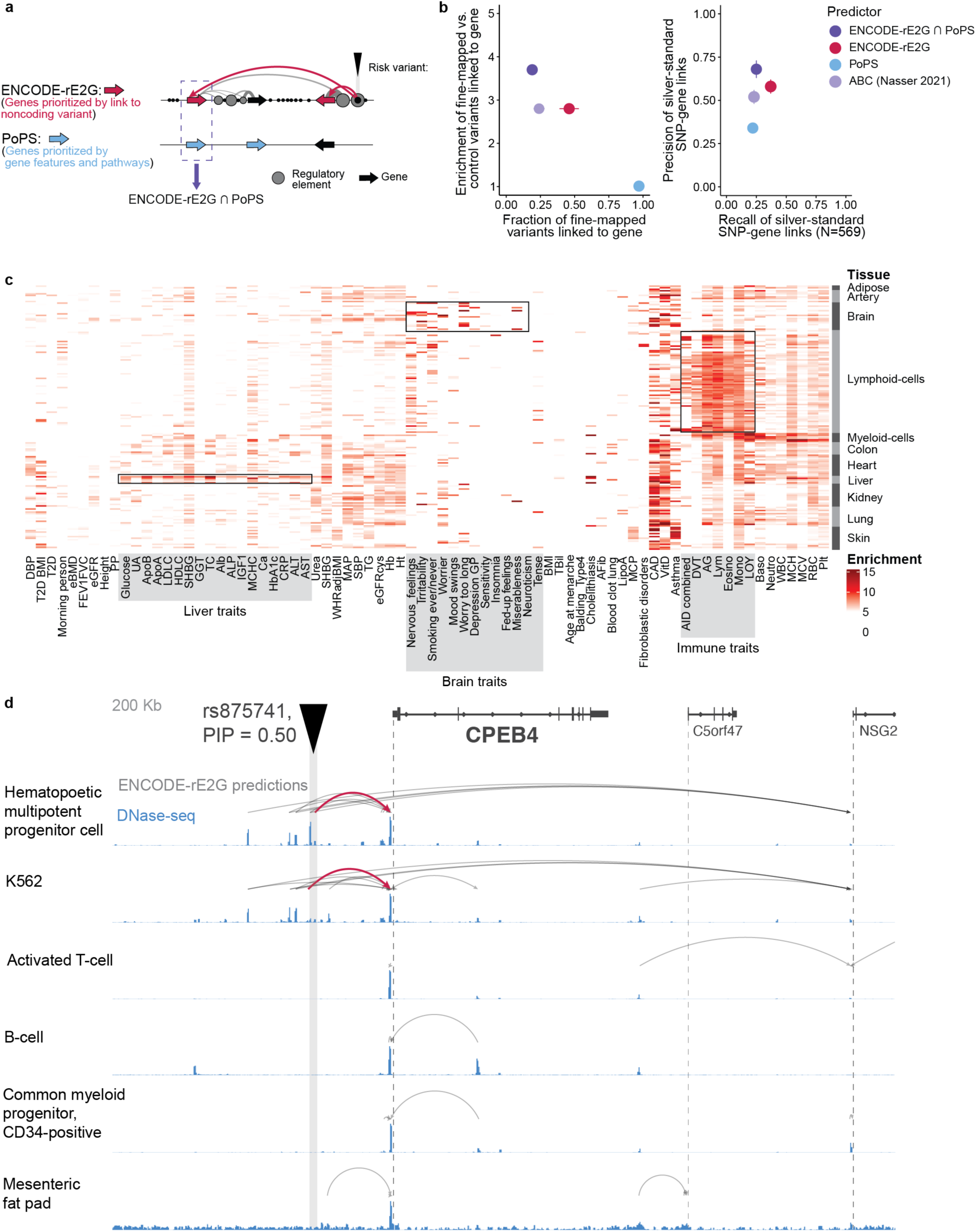
Linking noncoding GWAS variants to target genes and cell types. **a**, A schematic of how a risk variant for a disease is linked to a target gene by ENCODE-rE2G + PoPS, a combination of ENCODE-rE2G element-gene link and PoPS gene-level disease prioritization score. **b**, (left panel) Fraction of fine-mapped GWAS variants across 94 traits and the enrichment of this fraction with respect to control variants and (right panel) precision and recall in identifying the known causal gene for non-coding credible sets, as in Fig. 2f, for the ENCODE-rE2G + PoPS, and ENCODE-rE2G, PoPS and the previously published atlas of enhancer-gene links using ABC-Max^6^. For ENCODE-rE2G + PoPs we intersected the top two ENCODE-rE2G linked genes for predicted enhancers overlapping PIP>0.1 variants in the credible set with the top two PoPs prioritized genes in a 1Mb window around the credible set. For ENCODE-E2G and PoPs alone, we only considered the top one gene for each method. **c,** Heatmap representing the enrichment of GWAS-finemapped variants (PIP > 0.1) that are linked to a gene by the ENCODE-rE2G + PoPS method for each of 76 GWAS traits (along the columns) and 172 biosamples that are mapped to distinct tissues of origin (grouped along the rows). Only enrichments that are FDR significant (FDR < 0.10) in significance are shown. **d**, An example fine-mapped causal variant for mean corpuscular hemoglobin, rs875741, linked to the gene *CPEB4* (the top-scored PoPS gene in the locus) by ENCODE-rE2G only in hematopoietic progenitors and not in other closely related blood cell types like B-cells, T-cells and K562. Numerical results for each of the Figure panels are reported in **Table S13**.

This combination of ENCODE-rE2G and PoPS predicted a target gene for 3,528 out of 13,012 non-coding credible sets with a PIP > 0.1 fine-mapped variant spanning all traits; this included 5,851 out of 29.2K fine-mapped GWAS variants (**Table S13**), representing a 3.7-fold excess overlap compared to control variants (all distal noncoding variants detected in the 1000 Genomes project, *N*=9.2 million) (**Fig. 4b**). ENCODE-rE2G + PoPS identified known links between credible sets and target genes (**Fig. 2f**, **Fig. 4b**) with a precision of 68%, 1.2× and 2.0× higher than the top ENCODE-rE2G or PoPS gene alone, respectively. It also attained higher precision and comparable recall compared to our previous atlas generated with the ABC-Max algorithm^6^, including new predictions for a total of 1,811 additional credible sets that were not identified in our previous study^6^. As expected, variants were enriched for overlapping ENCODE-rE2G enhancers in expected tissues (**Fig. 4c**), generalizing the results of **Fig. 2e**; for example, mean corpuscular hemoglobin (MCH) showed 17× enrichment in ENCODE-rE2G enhancers in hematopoietic multipotent progenitors, 2.9× higher than other cell types on average. Examining individual loci identified a GWAS causal variant *rs875741* for MCH (PIP = 0.50) that had a predicted ENCODE-rE2G link to *CPEB4* in hematopoietic progenitors but not in other cell types like B-cells, T-cells and K562 (**Fig 4d**). This cell-type specificity is interesting because *CPEB4* has shown low expression specificity across human tissues^31,32^. Previous studies show that *CPEB4* knockdown inhibits terminal erythroid differentiation^33^ and lipid accumulation in adipocytes^34^, but the variant associated with MCH is predicted to overlap an enhancer specific to hematopoietic progenitors and not observed in adipocytes. Other interesting examples at the *TFRC, KIT* and *BCL11A* loci are reported in **Note S7 (Fig S7.2**). Thus, ENCODE-rE2G predictions can be combined with gene-to-pathway information to provide a unique resource for accurate inferences about causal genes and cell types.

We provide further analysis and guidelines regarding the value of adding new cell types to the ENCODE resource, assessing tissue-specificity of the ENCODE-rE2G biosample predictions, evaluating other combinations of ENCODE-rE2G and PoPS, and restricting our ENCODE-rE2G analyses to top disease-enriched biosamples for identifying possible cell types and target genes for GWAS loci in **Note S7 (Fig S7.1)**.

## Molecular features guiding enhancer-gene regulation

Toward understanding and advancing future modeling of molecular mechanisms of enhancer-promoter communication, we next sought to identify and explore the features that drive predictions of the ENCODE-rE2G classifiers (**Fig. 5, Fig. S4.1-2**, **Note S4, Note S5**).

**Figure 5.**
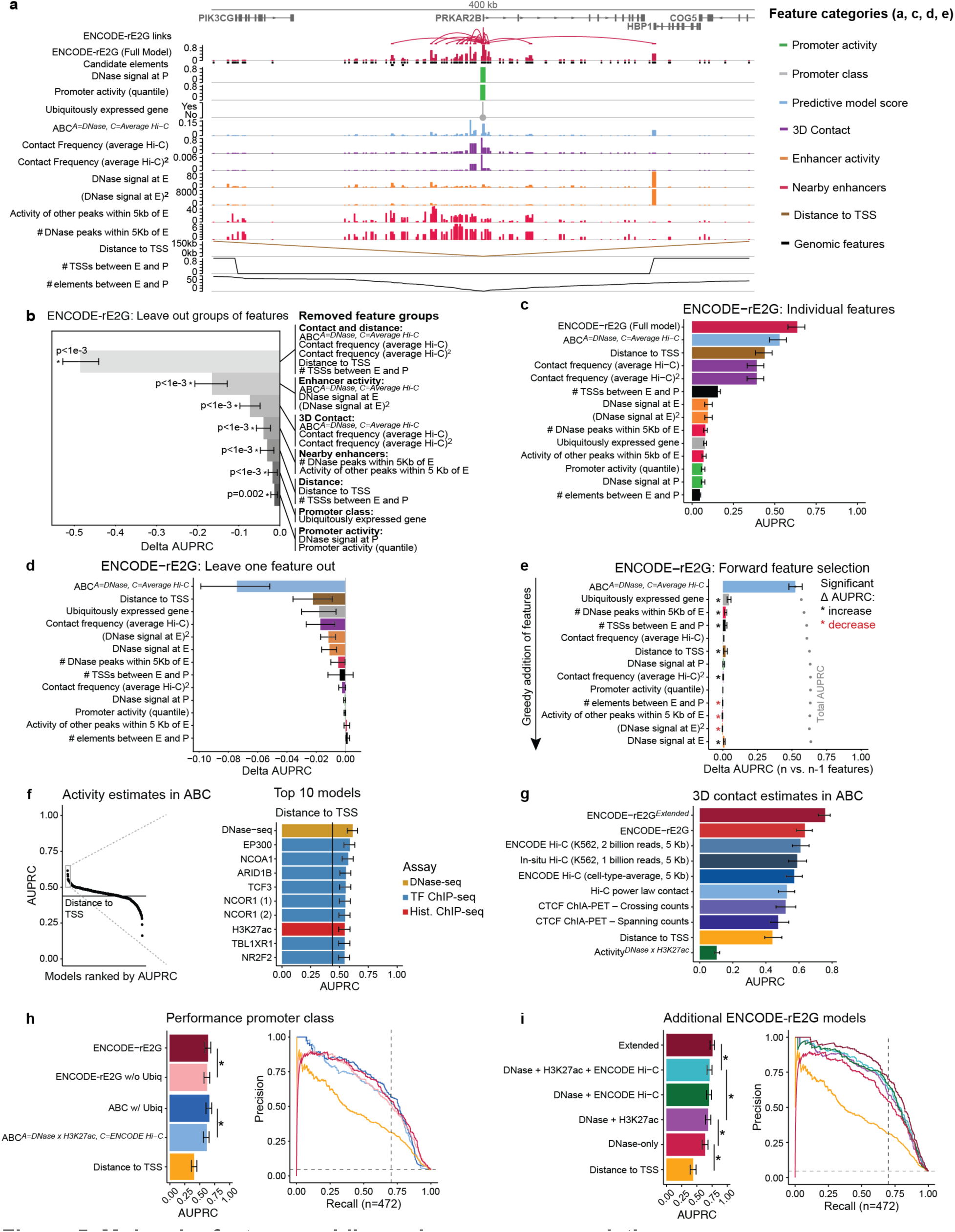
Molecular features guiding enhancer-gene regulation. **a**. ENCODE-rE2G features for the *PRKAR2B* locus. Only features for *PRKAR2B* E-G pairs are displayed. **b.** Effect of different feature categories on performance of the ENCODE-rE2G model in classifying K562 enhancer-gene regulatory interactions. Each feature category is removed from the feature set used by ENCODE-rE2G to see the amount of performance reduction. The numbers inside the parentheses show the numbers of the deleted features. Bar plots of delta AUPRC. Bar plots and error bars show the mean of delta AUPRC and 95% confidence intervals when randomly subsampling the EG pairs without replacement (bootstrapping) 1000 times. **c.** Performance of individual ENCODE-rE2G features at predicting enhancer-gene regulatory connections from CRISPR data. Bars show AUPRC of non-binary features. Error bars represent 95% range of AUPRC values inferred via bootstrap (1000 iterations). **d.** ENCODE-rE2G’s leave one out performance. Each feature of ENCODE-rE2G is ablated individually and the difference in AUPRC (Delta AUPRC) is calculated. Error bars represent 95% range of AUPRC values inferred via bootstrap (1000 iterations). Features with significant AUPRC reduction (P<0.05) are marked by stars. **e.** Sequential feature selection performed on ENCODE-rE2G, in which features were selected one at a time based on which one yields the greatest AUPRC. Bars show the change in AUPRC from the previous model including features above it. Gray dots show the total AUPRC for the model including the feature labeled and all above it. See Methods, ‘CRISPRi benchmark’ section for description of statistical test. **f.** Performance (AUPRC) of Activity-By-Contact (ABC) models using different ENCODE chromatin assays to estimate enhancer activity at predicting experimental results of CRISPRi data in K562 cells. Solid lines correspond to AUPRC of the distance to target TSS baseline predictor. For *NCOR1* data from two separate experiments (1, 2) were included. Error bars in barplot represent 95% range of AUPRC values inferred via bootstrap (1000 iterations). **g.** Performance (AUPRC) of different 3D features at classifying regulatory E-P interactions from CRISPRi perturbations in K562 cells (n = 10,292 pairs). Each feature, apart from ENCODE-rE2G and ENCODE-rE2G*^extended^*, represents different 3D contact measurements incorporated into the ABC model as the contact component. Error bars represent 95% range of AUPRC values inferred via bootstrap (1000 iterations). **h.** Precision recall curves showing that the addition of whether a gene ubiquitously expressed or not to the ABC model (hot pink curve) significantly improves on the performance of ABC (blue) (AUPRC increased by 0.044, *P* < 1.0 x 10^−4^). Error bars in barplot represent 95% range of AUPRC values inferred via bootstrap (10,000 iterations). See Methods, ‘CRISPRi benchmark’ section for description of statistical test. **i.** Performance of ENCODE-rE2G models using different assays to measure enhancer activity and 3D contact between elements and gene promoters at predicting results of CRISPRi data in K562 cells. Error bars in barplot represent 95% range of AUPRC values inferred via bootstrap (1000 iterations). See Methods, ‘CRISPRi benchmark’ section for description of statistical test.

We performed a series of analyses to gauge the importance of features in ENCODE-rE2G, including removing groups of related features (**Fig. 5b**), assessing the performance of each feature alone **(Fig. 5c**), removing one feature at a time (**Fig. 5d**), or performing sequential feature selection (**Fig. 5e**). The most informative features groups were 3D contact/distance and enhancer activity (**Fig. 5b**), and the most informative individual feature was the ABC score itself (**Fig. 5c-e**). These results are consistent with the importance of enhancer activity and 3D contact in enhancer regulation and highlight the utility of combining them as in the ABC model (**Fig. 2b,c, Fig S4.3**).

While we designed ENCODE-rE2G to use particular assays for enhancer activity and 3D contact (including DNase-seq and Hi-C), ENCODE has collected hundreds of other assays that could potentially be used. To compare the performance of other assays, we tested variations of the simpler ABC model in K562 cells in which we substituted different ENCODE assays for the enhancer activity or 3D contact frequency components:

(i) Out of 513 ENCODE 1D chromatin experiments that could represent enhancer activity, we found that DNase-seq and H3K27ac ChIP-seq were indeed among the best performing assays (**Fig. 5f**). Other ChIP-seq assays (for *EP300*, *NCOA1*, *NCOR1*, and other transcription factors) achieved equal performance (**Fig. 5f**, **Fig. S11**). Notably, ATAC-seq performed worse than DNase-seq in the ABC model (precision at 70% recall = 52% vs 41%, **Fig. S11a**).
(ii) Out of 6 methods for estimating 3D enhancer-promoter contact frequency in the ABC model, we found that the best dataset was cell-type-specific ENCODE Hi-C (precision at 70% recall = 53.8%; Hi-C depth = 2 billion reads, 5-kb resolution), which outperformed other datasets including *in situ* Hi-C (1 billion reads) or an inverse function of distance (precision at 70% recall = 50.4% or 31.8%, respectively) (**Fig. 5g**, **Note S2**). Incorporating Hi-C data led to particular improvements in performance for element-gene pairs located at longer distances (*e.g.*, >100 kb, **Fig. S2.1f,g**).

Beyond enhancer activity and 3D contact, additional features contributed to the performance of ENCODE-rE2G and ENCODE-rE2G*^Extended^*— both of which significantly outperformed the ABC model alone (**Fig. 2b,c**). Key additional features were related to promoter class and enhancer-enhancer interactions (**Fig. 5b,h, Fig S4.4**). We explore these two features below:

## Ubiquitously expressed genes are less sensitive to distal enhancers

ENCODE-rE2G and ENCODE-rE2G*^Extended^* learned that 3 features related to promoter class — whether the gene is ubiquitously, uniformly expressed across cell types, whether the promoter is predicted to be sensitive to distal enhancers in ExP-STARR-seq assays^35^, and whether the regulatory landscape around a gene is highly correlated across cell types^6^ — all predict that a gene is less likely to have distal regulatory elements. For example, ubiquitously expressed genes were 5-fold less likely (Fisher’s Exact Test *P* = 6.2 x 10^−45^) to have distal regulatory elements in the K562 CRISPR dataset and 1.13-fold less likely (Fisher’s Exact Test *P* = 0.0006) (**Fig. S6.1b**) to have a fine-mapped eQTL variant in a distal accessible site (**Fig. S6.1a**). Removing these features from ENCODE-rE2G*^Extended^* or ENCODE-rE2G led to a significant drop in performance (AUPRC decreased by 0.0140 or 0.016, *P_bootstrap_* = 0.002 or 0.0075, respectively), and combining them with the ABC score in a 4-feature logistic regression model led to a significant improvement (AUPRC increased by 0.044, *P* = 1.0 x 10^−4^) (**Fig. 5h, Fig. S6.1c-f**). Together, these observations support a model in which sequence-intrinsic features of the promoters of ubiquitously expressed genes make them less sensitive to distal enhancers^35,36^ (see **Note S6**).

## Super-additive functions of distal enhancers

For nearby enhancer activity, ENCODE-rE2G learned that 2 features related to enhancer-enhancer interactions — specifically, the number and summed activity of all other elements within 5 kb of the perturbed element — indicated that the perturbed element was more likely to have a regulatory effect on gene expression (**Fig. 5b**). Removing these features from ENCODE-rE2G led to a significant drop in performance (decrease in AUPRC of 0.0385, *P_bootstrap_* < 0.001). The importance of these features could be consistent with previous proposals that enhancers near one another in the genome can act in a synergistic manner^37–39^, and contrary to an assumption of the ABC model which assumes that enhancers act additively and independently to regulate gene expression^9^.

We analyzed and conducted additional genetic perturbation experiments to explore whether and in which cases enhancers might have super-additive effects on gene expression — here, defined as two enhancers, when added to a gene, having a larger effect than expected based on the sum of their individual effects in linear gene expression space (equivalently, such enhancers would appear “sub-subtractive” in perturbation experiments, where combined inhibition of both enhancers leads to a smaller decrease in expression than expected from an additive model). We found multiple lines of evidence for such effects:

(i) Of the 20 CRISPRi tiling experiments in which all nearby candidate enhancers near a gene were perturbed^9,19,25^, we identified 10 genes where the sum of the effect sizes of individual enhancers linked to a given gene was greater than 100% (**Fig. 6a**). This indicates that at least some pairs of enhancers at these loci have super-additive effects.
(ii) We performed combinatorial CRISPRi perturbations to all pairwise combinations of 7 enhancers near the *MYC* gene in K562 cells using CRISPRi-FlowFISH^9^ (**Fig. 6b,c**). All of the 21 *MYC* enhancer pairs displayed evidence of significant super-additive interaction effects on gene expression (Benjamini-Hochberg corrected *P* < 0.001, *F*-test), although to differing degrees (**Fig. 6d,e**). Stronger interaction effects (16-37% difference from additive model) were observed for pairs of enhancers closer in genomic distance (3–89 kb) and with higher 3D contact frequency, whereas weaker interaction effects (2-10% difference from additive model) were observed for enhancers that were located farther from one another (107 kb–1.79 Mb) (**Fig. 6f**). This suggests that enhancers nearby to one another in the genome have super-additive effects on gene expression, whereas more distant enhancers combine approximately additively.

**Figure 6.**
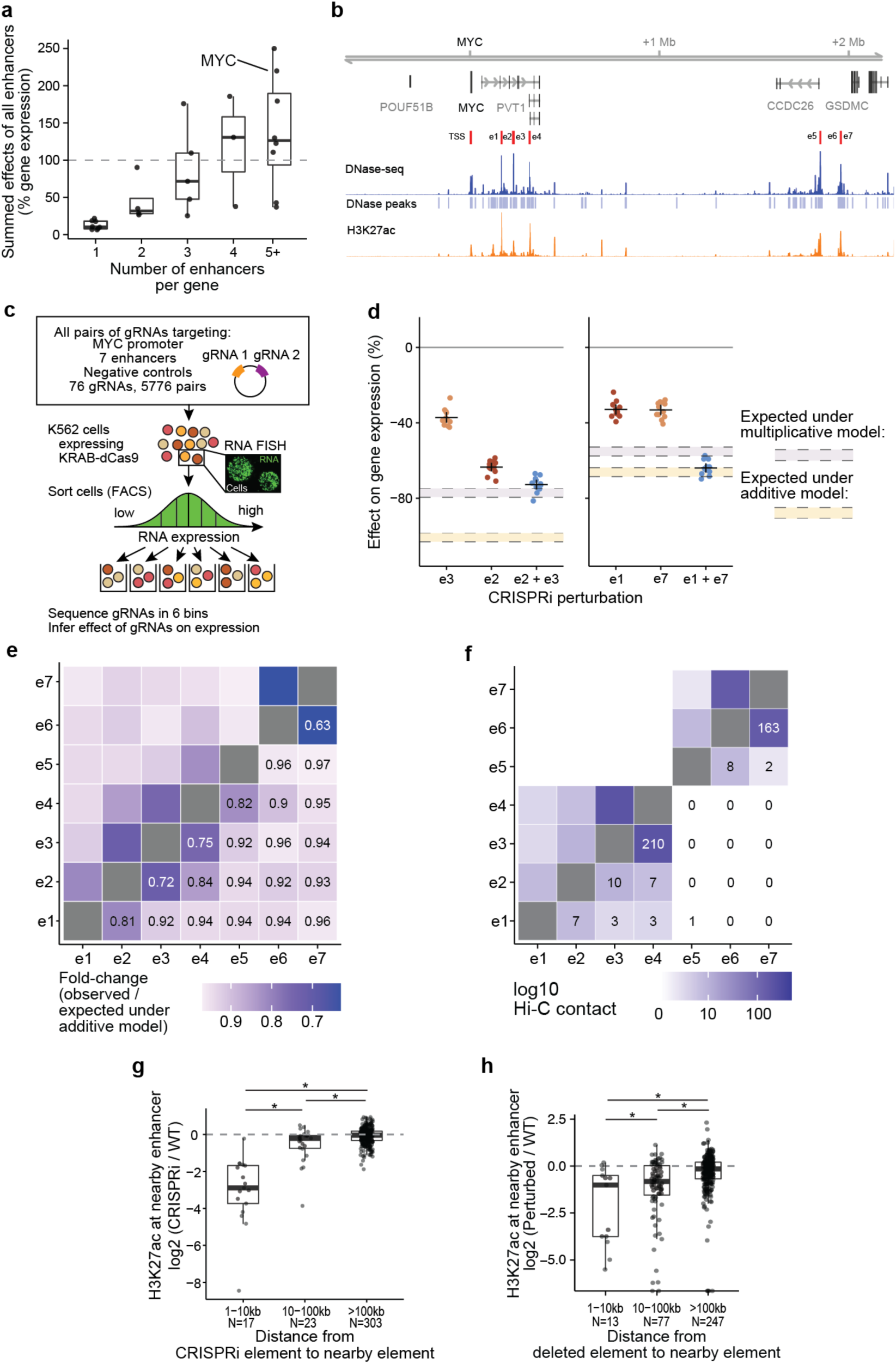
Super-additive functions of distal enhancers. **a**. Box plots showing the combined effect size on expression of a given gene from CRISPRi experiments targeting all enhancers of that gene^6,9,19^ by the number of enhancers. **b.** Acetylation and DNase-seq peaks at the *MYC* locus in K562. **c.** Experimental design panel. Individual guides targeting a specific element at the *MYC* locus are paired with every other guide and transduced into K562-KRAB-dCas9 cells. Effects are measured with CRISPRi-FlowFISH. **d.** Individual and combined effects on gene expression from perturbing e2 and e3 (left), and perturbing e1 and e7 (right), with the expected effect under additive and multiplicative models annotated. **e.** Heatmap: Fold change in observed pairwise effects on gene expression versus expected pairwise effects under an additive model from paired CRISPRi screen. **f.** Heatmap: Normalized Hi-C contact (5 kb resolution) between *MYC* enhancers in K562 cells. **g.** Box plots showing the effect of CRISPRi perturbations to an enhancer on H3K27ac at a nearby enhancer stratified by distance^25,40^. There is a significant difference in the distributions of perturbation/enhancer distances between 1 and 10 kb and distances greater than 100 kb distance bin (two-sided Wilcoxon rank sum exact test, *P*=1.79×10^−9^), between the between distributions for distances between 1 and 10 kb and 10 and 100 kb (*P*= 7.37×10^−6^), and between the distributes for distance between 10 kb and 100 kb and greater than 100 kb (*P*=0.031). See Methods, ‘Data visualization’ section for definition of box plot elements. All data points are shown in addition to box plots. **h.** Box plots showing effect of CRISPR-Cas9-mediated knockout of enhancers on nearby elements, stratified by distance^41^. There is a significant difference between distributions in each distance group (two-sided Wilson rank sum test with continuity correction, 1-10 kb vs 10-100 kb, *P* = 0.0077; 10-100 kb vs >100 kb, p=0.0026; 1-10 kb vs >100 kb, p=0.00025). See Methods, ‘Data visualization’ section for definition of box plot elements. All data points are shown in addition to box plots.

To further understand how enhancers near one another in the genome could have super-additive effects on expression, we analyzed their effects on chromatin state. We performed and collated experiments where enhancers were perturbed with CRISPRi, genetic deletions, or naturally occurring genetic variants and effects on nearby enhancers measured with H3K27ac ChIP-seq^40–42^. In each dataset, perturbations to individual enhancers on average reduced H3K27ac signal at other nearby enhancers, and the magnitude of the effect decreased as enhancer-enhancer distance increased (**Fig. 6g,h, Extended Data** Fig. 8).

Together, these data indicate that enhancers near one another in the genome often influence each other’s chromatin state and have super-additive effects on gene expression. These effects could explain the importance of enhancer-enhancer interactions in the ENCODE-rE2G model, and highlight how interpreting the model can guide mechanistic explorations of enhancer function.

## Applying ENCODE-rE2G to new cell types

Toward guiding further data collection and model application to new cell types, we systematically compared the marginal utility of adding assays to ENCODE-rE2G. We constructed logistic regression classifiers with different combinations of DNase-seq, H3K27ac, and Hi-C as cell-type specific input datasets in K562 cells. Adding either cell-type specific H3K27ac and/or Hi-C data in addition to DNase-seq data led to significant improvements in model performance, nearing the performance of ENCODE-rE2G*^Extended^* on the CRISPR benchmark (**Fig. 5i**). To facilitate constructing ENCODE-rE2G models in additional cell types in the future, we provide code for computing the input feature tables and applying pre-trained models for each combination of input datasets (see **Code Availability**).

## Discussion

Here we presented the ENCODE encyclopedia of enhancer-gene regulatory interactions in the human genome. This resource includes: (i) New improved models that can be applied to new cell types using DNase-seq data alone, along with guidelines and software for construction of new enhancer-gene maps; (ii) Benchmarking datasets and pipelines that will enable systematic comparisons and iterative improvements to enhancer-gene models; and (iii) Genome-wide maps of enhancer-gene regulatory interactions across ENCODE biosamples to identify enhancers for genes of interest, find genes regulated by enhancers of interest, and interpret the functions of human genetic variants.

Identifying enhancers and their target genes has been a long-standing challenge in genomics^43^. Over its 20-year history, the ENCODE Project has generated key datasets that enabled the development of predictive models to address this challenge. ENCODE data about chromatin states across cell types led to the first models to link enhancers to target genes genome-wide by correlating measures of enhancer activity with gene expression across tissues, cell types, or conditions^13,14,44–47^. ENCODE maps of 3D chromosome contacts provided initial pictures of the physical interactions between enhancers and promoters^48–50^ and gave rise to predictive models aimed at identifying these physical interactions^15,51^. The availability of maps of both chromatin state and 3D contacts enabled the development of models that began to combine these features, including ABC and others^9,23^. Now, high-throughput CRISPR screens have provided a sufficiently broad survey of regulatory effects directly in the genome to enable our supervised learning framework to find optimal combinations of assays, processing methods, and feature extraction approaches to make reliable predictions of enhancer-gene regulatory interactions. We expect that this ENCODE resource of harmonized data, models, and benchmarks will continue to play an important role in future efforts to develop improved models of gene regulation.

Our analysis, together with previous data, suggest a simple set of rules that specify enhancer-gene regulatory interactions in the human genome. Enhancers activate promoters dependent on their 3D contact frequencies, as in the classic looping model, with contact frequencies determined by various factors including genomic distance, loop extrusion, and the positions of CTCF sites^52^. The effect of perturbing an enhancer on gene expression is, to a first approximation, proportional to 3D contact frequency and the “activity” of the enhancer, as described by the ABC model^9^. This effect is tuned by at least two additional factors. First, promoters of ubiquitously expressed genes are less responsive to distal enhancers (**Fig. S6.1b**), possibly because they contain built-in motifs for activating transcription factors that buffer them against distal enhancers^35^. Second, the activity of an enhancer is increased by the presence of other nearby enhancers, which appear to act super-additively with those in close 3D proximity (**Fig. 6**). These four factors — intrinsic enhancer activity, enhancer-promoter 3D contacts, promoter class, and enhancer-enhancer interactions — together can explain a substantial fraction of the regulatory effects observed in CRISPR perturbation datasets.

Our observations regarding super-additive effects of enhancers require particular care in interpretation. Previous studies have investigated how enhancers might combine additively, synergistically, or redundantly by assessing effects at the level of gene expression^39,53–57^, cellular phenotypes^37^, or organismal phenotypes^58,59^. These studies have offered conflicting views, perhaps in part because these higher-order phenotypes might have varying non-linear relationships with quantitative gene expression. Here we studied pairs of enhancers using a highly quantitative assay with good power to distinguish between these models directly at the level of gene expression (**Fig. 6d-h**). We observe enhancers that show pairwise effects ranging from approximately additive to approximately multiplicative in linear gene expression space, showing that different effects are possible even at a single gene. The interaction effects between enhancers appear to depend on their 3D proximity to one another, where enhancers that contact frequently combine super-additively and enhancers that contact less frequently combine additively (**Fig. 6g,h**). Notably, neither the single-guide CRISPR datasets nor the paired-guide CRISPR data offer support for a model where enhancers have “redundant” effects at the level of gene expression: in the single-guide CRISPR data, many enhancers have significant individual effects on expression, and in the dual-guide CRISPR data we do not observe any cases of sub-additive effects.

Our study highlights the importance of benchmarking models using genetic perturbation data. Here, we developed and validated ENCODE-rE2G using an array of independent datasets across multiple tissues and cell types, including CRISPR perturbations and datasets about the impact of genetic variants. However, these genetic perturbation datasets have limitations. The CRISPR datasets still have a relatively small number of positive examples; were designed with certain biases in enhancer and gene selection; do not distinguish *cis* from *trans* effects; are conducted largely in a single cell type (K562); and are underpowered to detect regulatory interactions with small effect sizes (*e.g.*, <25%) (see **Note S1**). The predictions of the ENCODE-rE2G model could therefore be biased in similar ways. Similarly, the eQTL datasets^27^ are largely from tissues, lack cell-type resolution, and can be underpowered to detect small effects. Future efforts to collect larger, unbiased CRISPR perturbation and eQTL datasets with resolution for individual cell types are needed to train improved models.

In summary, this study develops a new approach to map enhancer-gene regulatory interactions in the human genome, and provides an expansive resource for building, benchmarking, and applying such maps. Our strategy included collecting and labeling high-quality genetic perturbation data, establishing benchmarking pipelines in a collaborative framework, and developing predictive models that can be applied across human cell types. This strategy may be more generally applicable to learning rules of gene regulation beyond enhancer-gene regulatory interactions, including identifying the target genes of transcription factors, mapping effects of variants on chromatin state, and others. Continued efforts to build such a human gene regulation map will provide a foundation for understanding the biology of health and disease, programming gene expression, and interpreting the functions of genetic variants.

## Data availability

ENCODE-rE2G predictions and input epigenomics dataset are available on the ENCODE Portal (www.encodeproject.org). ENCODE file accessions are listed in **Table S2** and **Table S10.**

ENCODE portal accession ids and public links for GraphReg and EPIraction predictions are available in **Table S8** and **Table S9.**

The combined CRISPR dataset is available as part of the CRISPR benchmarking pipeline at https://github.com/EngreitzLab/CRISPR_comparison/blob/main/resources/crispr_data/EPCrispr <u>Benchmark_ensemble_data_GRCh38.tsv.gz</u>. The CRISPR data is also available on the ENCODE portal under the accession ENCSR998YDI.

For the eQTL benchmarking pipeline, GTEx eQTL variants and RNA expression data for each tissue used in the eQTL benchmarking pipeline are available on Synapse: https://www.synapse.org/#!Synapse:syn52264240

For the GWAS benchmarking pipeline, fine-mapped GWAS data for UK Biobank traits is available from https://www.finucanelab.org/data/.

Files containing baseline predictors used for benchmarking analyses can be found on Synapse: https://www.synapse.org/#!Synapse:syn52234396.

Data from the pairwise CRISPRi enhancer perturbation experiment at the *MYC* locus in K562 is available on the ENCODE portal under: ENCSR443VTK.

CRISPRi-H3K27ac ChIP-seq experiments at the *MYC* locus are available at the NCBI Gene Expression Omnibus, Accession GSE225157.

## Code availability

ENCODE-rE2G: https://github.com/karbalayghareh/ENCODE-rE2G

ABC: https://github.com/broadinstitute/ABC-Enhancer-Gene-Prediction

GraphReg: https://github.com/karbalayghareh/GraphReg

CRISPR benchmarking pipeline: https://github.com/EngreitzLab/CRISPR_comparison

eQTL benchmarking pipeline: https://github.com/EngreitzLab/eQTLEnrichment

GWAS benchmarking pipeline: https://github.com/EngreitzLab/ABC-Max-pipeline

EPIraction: https://github.com/guigolab/EPIraction

CIA and CCD features: https://github.com/wangxi001/CIA

EpiMap: https://github.com/KellisLab/EpiMap_GRCh38_linking

CRISPR data analysis: https://github.com/argschwind/ENCODE_CRISPR_data

## Supporting information

Table S1. Collected enhancer-gene predictions

Table S2. ENCODE-rE2G DNase-only Metadata

Table S3. Features of enhancers, promoters, and enhancer-promoter pairs

Table S4. Hi-C files for computing average interaction frequency

Table S5. Powerlaw gamma, scale values

Table S6. Input DNase-DNase correlation

Table S7. Input DNase-RNA correlation

Table S8. GraphReg_LR input data

Table S9. EPIraction metadata

Table S10. ENCODE-rE2G extended input data

Table S11. Baseline predictors input data and predictions

Table S12. CRISPR benchmark performance summary

Table S13. GWAS benchmarking and guidelines

Table S14. GTEx tissue and biosample pairs used in eQTL benchmarking

Table S15. GM12878 enrichment-recall table

Table S16. Baseline predictors GM12878 enrichment-recall table

Table S17. Enhancer activity ABC input files

Table S18. ENCODE-rE2G summary statistics per biosample

Table S19. ENCODE-rE2G summary statistics per gene

## Acknowledgements

We thank members of the ENCODE Consortium, Nuclear Architecture Working Group, and Distal Regulation subgroup for data generation, analysis support, and discussions to develop the manuscript. We thank Molly Gasperini for discussions about CRISPR screen analysis. We thank Anna Shcherbina, Kun Xiong, Sarah Gradesieck, Jacob Schreiber, Nate Tippens, Heini Natri, Ray Jones, and Gamze Gursoy for jamboree support and participation. We thank Georgi Marinov for consulting on enhancer synergy screens. M.U.S. acknowledges the support of an NSF Graduate Research Fellowship (DGE-1656518). K.K.D. acknowledges the support of R00HG012203, P30CA008748, and the Josie Robertson Investigator Fellowship. E.J. was supported by the Novo Nordisk Foundation (NNF21SA0072102). A.R.G. and L.M.S. acknowledge the support of 1R01HG011664-01A1 and the NHGRI Impact of Genomic Variation on Function Consortium (UM1HG011972). Z.W. and K.F. acknowledges the support of U24HG009446. A.Kundaje acknowledges support of U01HG009431 and U01HG012069. R.G. and R.N.N. acknowledge the support of the Spanish Ministry of Economy and Competitiveness (MEC) (BIO2011-26205). D.Y. acknowledges the support of the National Science Foundation Graduate Research Fellowship (DGE-1656518). B.T.J. was supported by the National Science Foundation Graduate Research Fellowship (Grant No. 1745302). B.R.D. was supported by the National Science Foundation Graduate Research Fellowship (DGE-1656518). A.L.P. acknowledges the support of NIH grant U01 HG012009. M.S.S. acknowledges the support of the Paul and Daisy Soros Fellowship. K.Andreeva acknowledges the support of U24HG009397 and U41HG006992. M.A.B. acknowledges the support of R01HG012367 and U01HG009380. C.S.L. acknowledges the support of U01HG009395 and U01HG012103. W.J.G. and M.C.B. acknowledge the support of NIH ENCODE UM1HG009436. J.M.E. acknowledges support from an NIH Pathway to Independence Award (K99HG009917 and R00HG009917); a NHGRI Genomic Innovator Award (R35HG011324); Gordon and Betty Moore and the BASE Research Initiative at the Lucile Packard Children’s Hospital at Stanford University; ENCODE UM1HG009436; the NHGRI Impact of Genomic Variation on Function Consortium (UM1HG011972); the Novo Nordisk Foundation (NNF21SA0072102); and the Chan Zuckerberg Initiative DAF (2022-249191), an advised fund of Silicon Valley Community Foundation.

## Author Contributions

A.R.G., K.S.M., A.Karbalayghareh, M.U.S., K.K.D., R.N., E.J., W.X., and J.M.E. conducted analyses and wrote the manuscript with input from all authors. A.R.G., K.S.M., and J.M.E. co-led the analysis group and contributed to most analyses in the paper. K.S.M. coordinated ENCODE epigenomics data download and processing. A.R.G. and J.N. developed the CRISPR benchmarking pipeline. M.U.S. and J.N. developed the eQTL benchmarking pipeline. K.K.D. and K.S.M. developed and applied the GWAS benchmarking pipeline. A.Karbalayghareh developed the ENCODE-rE2G logistic regression models. A.R.G., J.N., and B.R.D. collected and analyzed published CRISPR datasets. K.S.M. and W.X. compared the performance of 3D contact measurements. A.R.G. compared performance of enhancer activity measurements. M.U.S., H.J., J.N., and A.Karbalayghareh analyzed evidence for enhancer synergy. R.N., A.R.G., and A.Karbalayghareh analyzed the impact of correlation features. E.J. and K.S.M. analyzed the global properties of E-G predictions. E.J. and A.Karbalayghareh analyzed the impact of features related to ubiquitously expressed genes. K.S.M., M.U.S. A.S.T. and X.R.M. updated and applied the ABC model. B.J. and M.K. updated and generated predictions from the EpiMap model. R.N. and R.G. generated predictions from the EPIaction model. Z.A. and D.K. generated predictions from the Enformer model. A.Karbalayghareh and C.S.L. generated predictions from the GraphReg model and analyzed the impact of GraphReg features. W.X. and M.A.B. generated predictions from the CIA Model. D.Y. designed the paired-guide *MYC* screen. D.Y., H.J., and T.N. conducted experiments for the paired-guide *MYC* screen. H.J. analyzed the paired-guide *MYC* screen. E.M.P. designed and conducted ChIP-seq experiments for *MYC* enhancers. K.S.M, M.S.S., R.M., N.C.D., S.S.P.R., and E.L.A. contributed to analysis of Hi-C data. J.M.E., R.N., K.S.M., A.R.G., K.A., K.K.D., M.U.S., and A.Karbalayghareh, developed file formats for predictive models. J.C.U. and H.K.F. contributed fine-mapped eQTL data. K.F., J.E.M., and Z.W. contributed to analysis of ENCODE cCREs. A.L.P., M.A.B., R.G., A.Kundaje, L.M.S., C.S.L., and J.M.E. supervised members of the writing team. A.L.P., M.A.B., R.G., A.Kundaje, L.M.S., C.S.L., J.A.S., E.L.A., W.J.G. and J.M.E. contributed to data analysis and interpretation. J.M.E. provided overall supervision for the study.

## FIG

