## Supplementary Figures and Notes for "An encyclopedia of enhancer-gene regulatory interactions in the human genome"

### Extended Data Figures

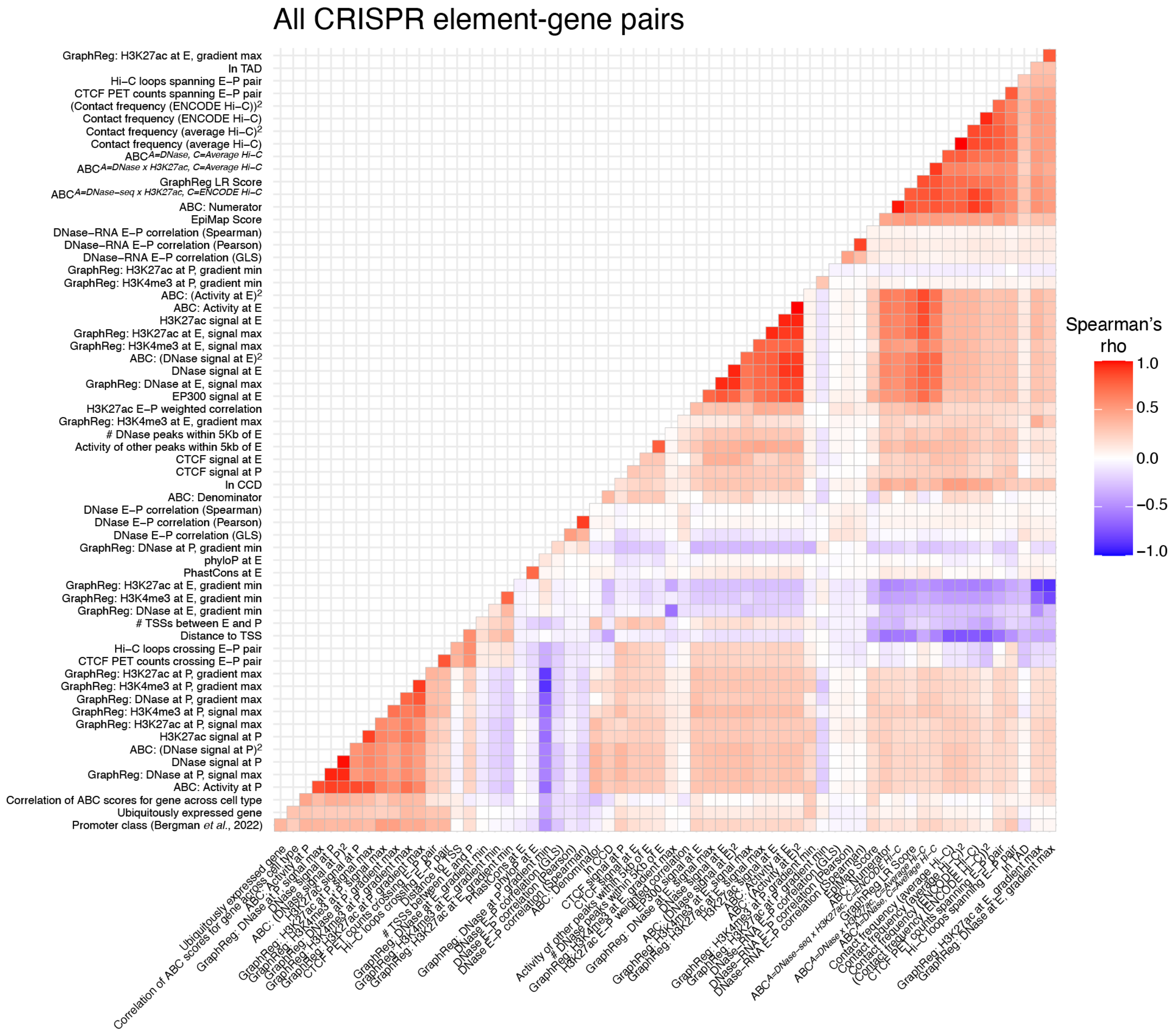


**Extended Data Fig. 1 | Correlation between features used as input for ENCODE-rE2G*Extended***
Correlation of collected and generated molecular features of element-gene pairs in combined CRISPR dataset.

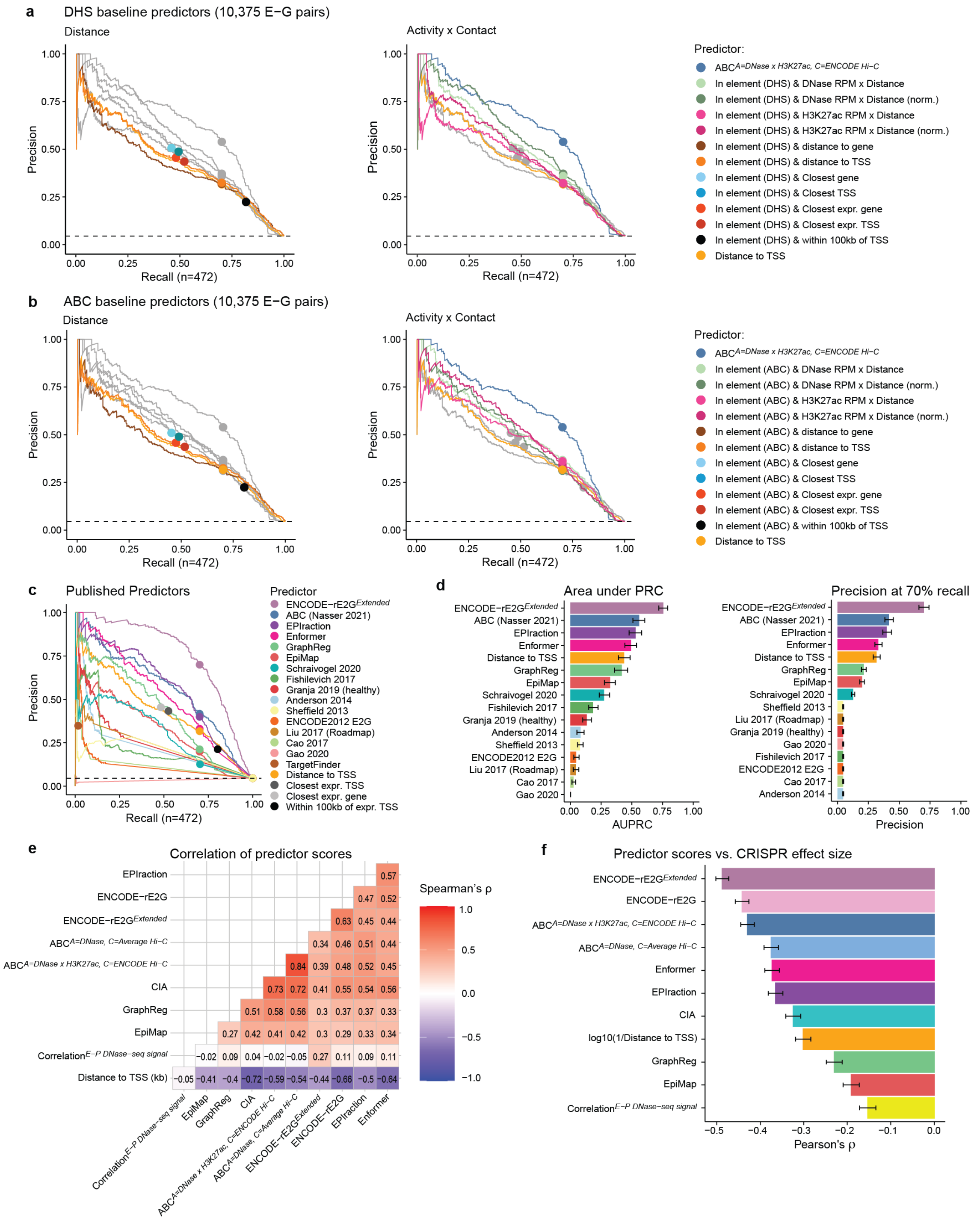


### Extended Data Fig. 2 | CRISPR benchmarking

**a**, Precision-Recall curves for simple baseline predictors based on ENCODE4 DNase hypersensitivity sites (DHS) grouped by genomic distance or activity (DNase-seq or H3K37ac ChIP-seq at DHS) x contact (1 / distance to TSS). The ABC model was added as reference.

**b**, Precision-Recall curves for simple baseline predictors based on ABC candidate elements grouped by genomic distance or activity (DNase-seq or H3K37ac ChIP-seq at element) x contact (1 / distance to TSS). The ABC model was added as reference

**c,** Precision-Recall curves for previously published semi-supervised predictors.

**d,** Area under Precision-Recall curve (AUPRC) and precision at 70% recall for previously published semi-supervised predictors. Error bars represent 95% range of AUPRC values inferred via bootstrap (1000 iterations).

**e,** Correlation (Spearman's ρ) between predictor scores for combined CRISPR E-G pairs.

**f,** Correlation (Pearson's ρ) of predictor scores vs observed CRISPRi effect sizes for combined CRISPR data. Error bars represent 95% confidence intervals.

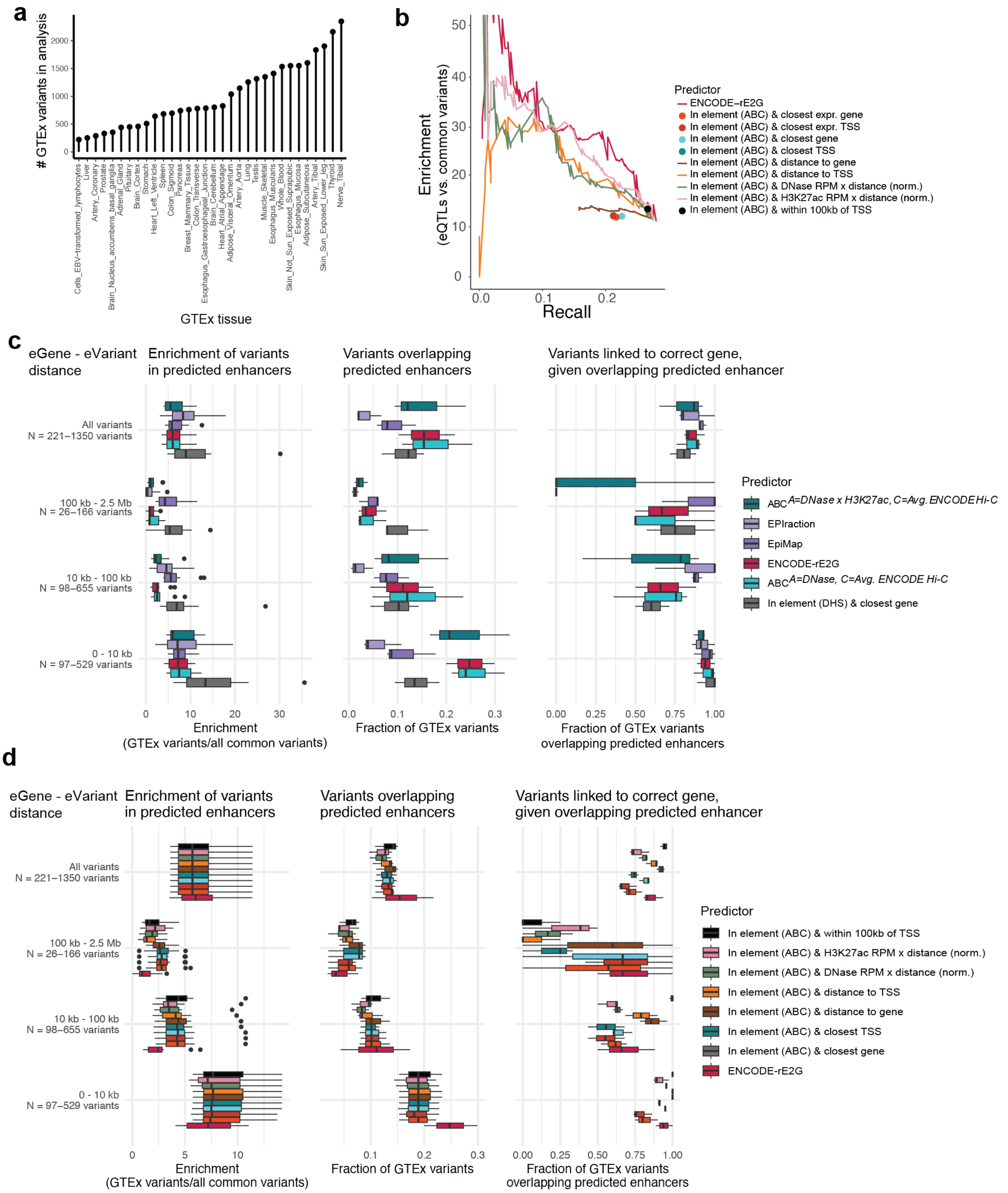


#### Extended Data Fig. 3 | eQTL Benchmarking

**a,** Number of unique fine-mapped GTEx variants used in eQTL benchmarking analysis for each tissue. Variants were filtered to those in distal noncoding regions, PIP>0.5, in a credible set, and linked to a gene expressed in the corresponding tissue.

**b,** Enrichment – recall curves showing the enrichment of fine-mapped distal noncoding eQTLs with a PIP>0.5 in EBV-transformed lymphocytes in predicted enhancers compared to distal noncoding common variants versus fraction of variants overlapping enhancers in GM12878 cells across different score thresholds for enhancers predicted by baseline models computed on candidate ABC enhancers. Numerical results are reported in **Table S16**.

**c,** Boxplots showing enrichment (fraction of variants overlapping any enhancer/fraction of common variants overlapping any enhancer), recall (fraction of variants overlapping any enhancer), and fraction of variants overlapping an enhancer linked to the correct gene given that they overlap an enhancer for predictive models across 12 tissue-biosample pairs. Results are stratified by distance between eVariant and eGene. See Methods, 'Data visualization' section for definition of box plot elements.
**d,** Same as (**c**), except applied to baseline predictors computed on candidate ABC enhancers. See Methods, 'Data visualization' section for definition of box plot elements.

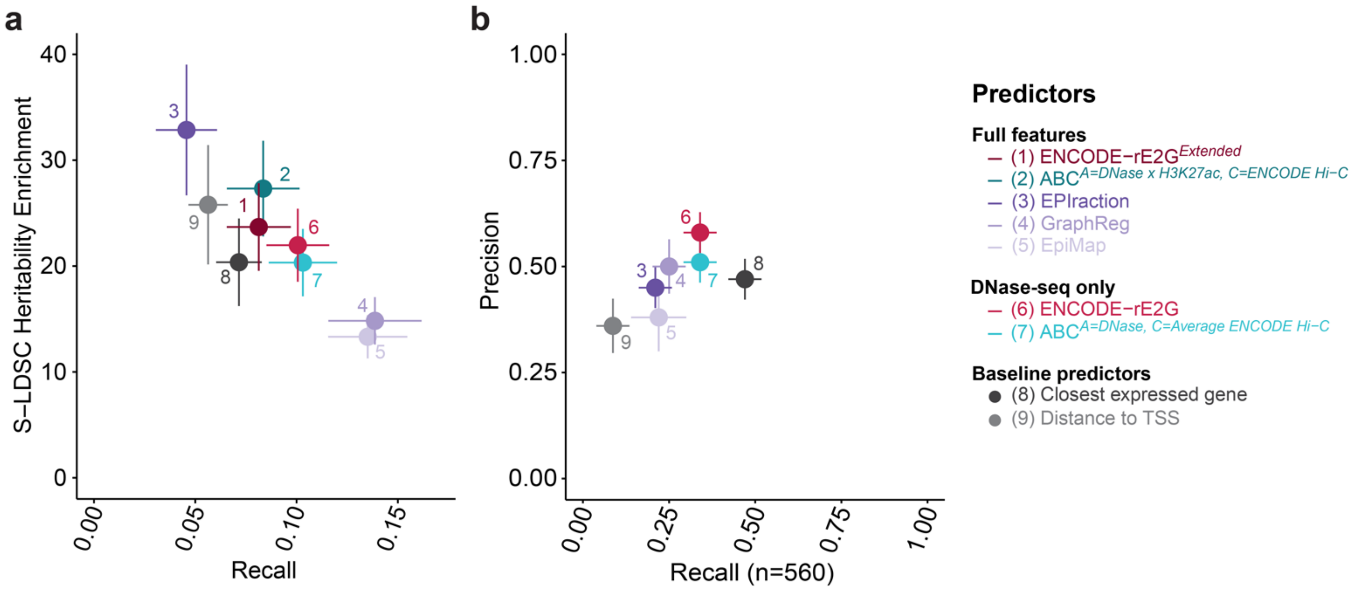


#### Extended Data Fig. 4 | GWAS Benchmarking

**a,** S-LDSC Heritability enrichment (Y-axis) and against recall of fine-mapped SNPs (X-axis) for all element-gene predictions in K562 for three RBC-related traits and GM12878 for two Lymphocyte related traits respectively.
**b,** A re-analysis of the benchmarking test in Figure 2f, where we consider all 560 non-coding credible sets corresponding to 31 traits that are linked to exactly one "putatively causal" gene with a coding fine-mapped variant within 2Mb on either side of the lead variant. We compute the precision and recall in linking it to this gene for each of the element-gene predictions in blood biosamples (see Methods). Error bars represent 95% confidence intervals. Numerical results are reported in **Table S13**.

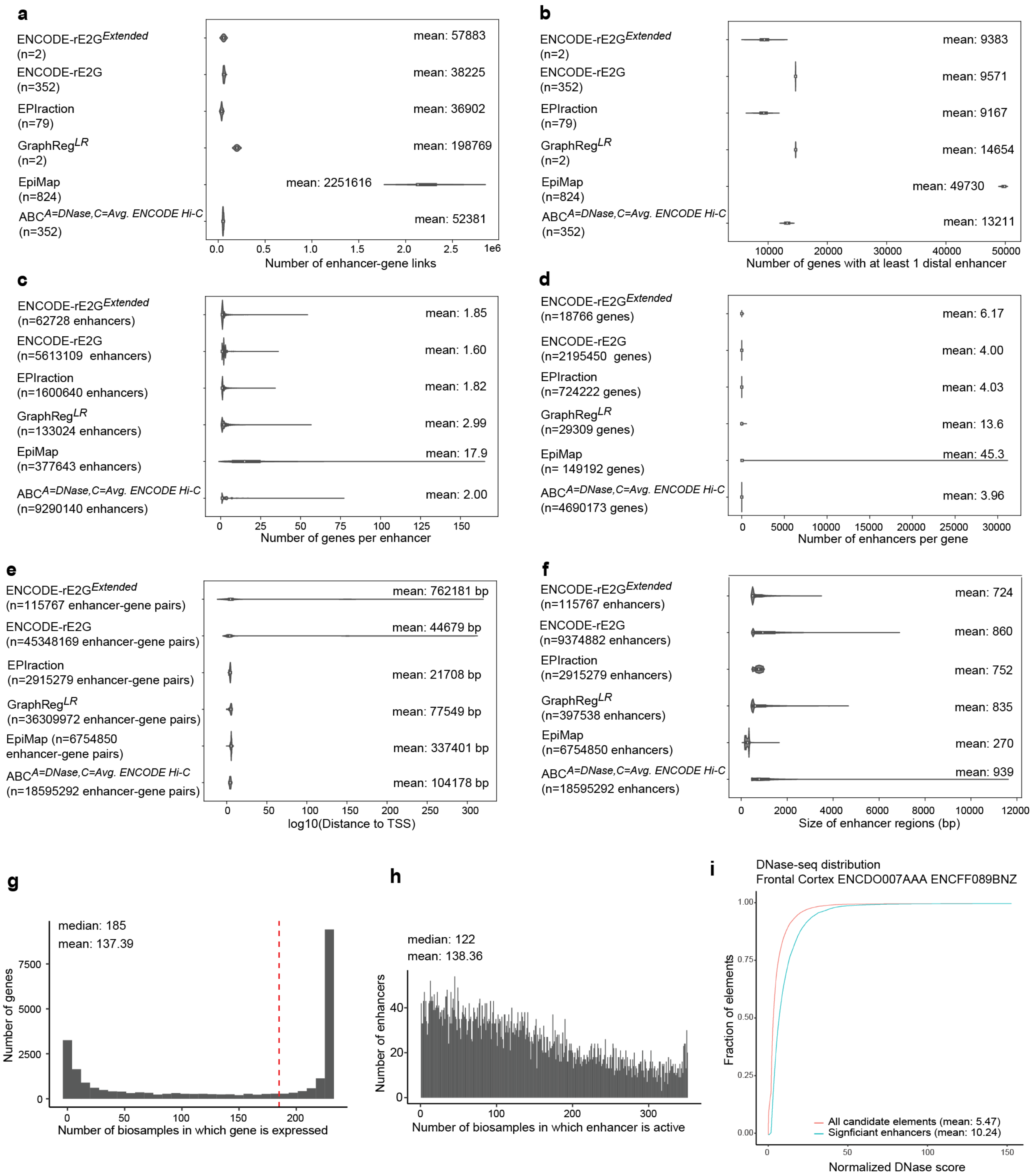


**Extended Data Fig. 5 | Properties of enhancer-gene regulatory interactions for different models**

**a,** Distribution of “enhancer-gene links per biosample” across all biosamples

**b,** Distribution of number of genes that is “regulated by at least one enhancer in a given biosample” across all biosamples

**c,** Distribution of “number of genes per enhancer in a given biosample”, including all genes in all biosamples that have at least 1 enhancer in that biosample

**d,** Distribution of “number of enhancers per gene in a given biosample”, including all genes in all biosamples that have at least 1 enhancer in that biosample

**e,** Distribution of “distance to transcription start site (TSS)” for all enhancer-gene pairs in all biosamples

**f,** Distribution of “enhancer region size” across all biosamples
**g,** Gene Expression Across Biosamples. Genes were considered expressed in a given biosample if TPM > 1.
**h,** ENCODE-rE2G enhancer activity across biosamples. To account for shifting peak boundaries across biosamples, every element that was called as a significant enhancer element in at least one biosample was intersected with all other significant enhancer elements using bedtools intersect. Enhancers were considered the same if their boundaries were at least 50% overlapping in different biosamples and their biosample lists were aggregated to get the final biosample count.
**i,** DNase-seq score for all ENCODE-rE2G candidate elements and all significant ENCODE-rE2G enhancers in the frontal cortex.

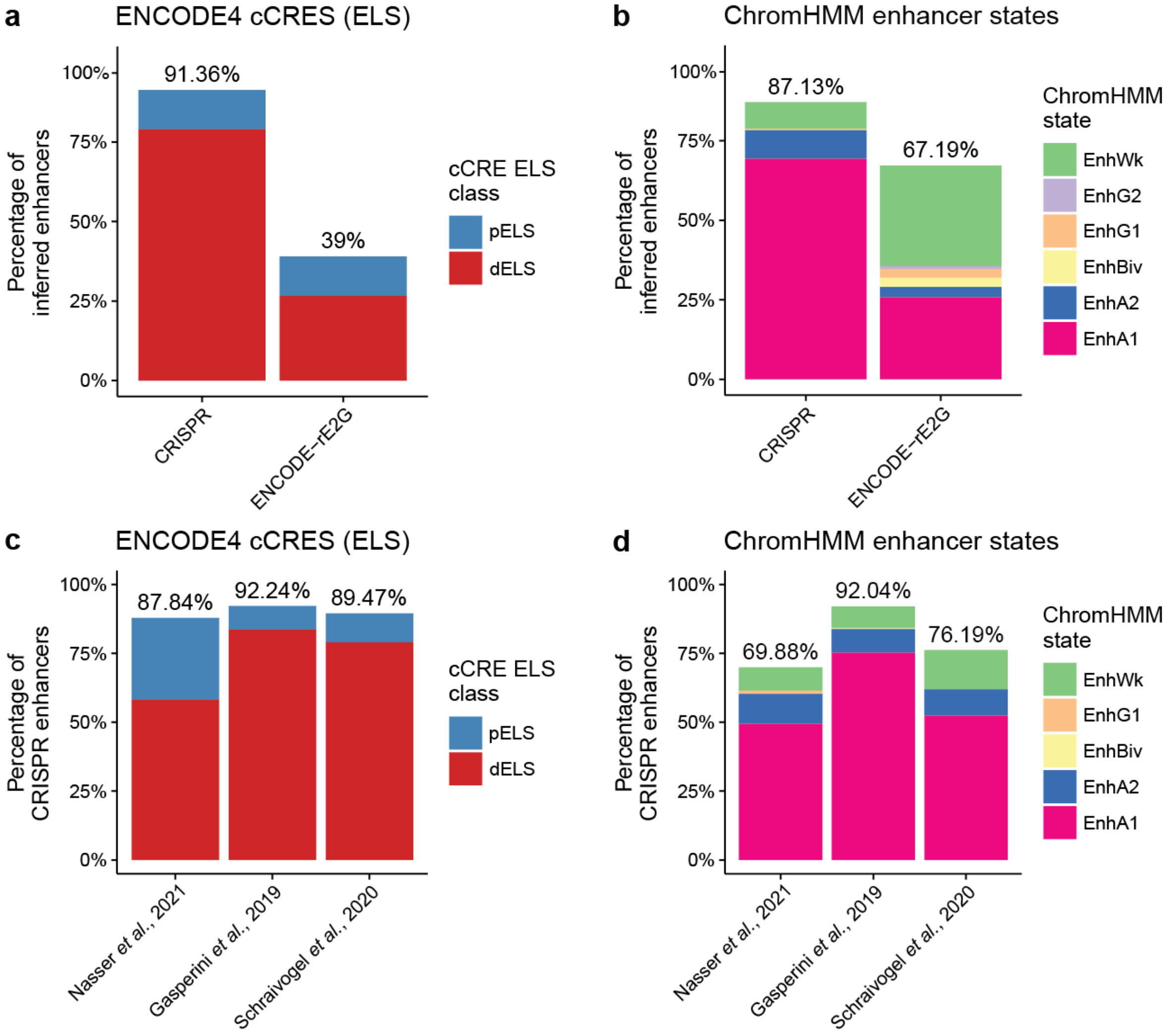


**Extended Data Fig. 6 | Comparison of ENCODE-rE2G to other definitions of enhancer regions**
**a,** Percentage of K562 enhancers predicted by ENCODE-rE2G or inferred from CRISPR data that overlap K562 ENCODE4 proximal or distal enhancer-like signatures (pELS, dELS).
**b,** Like (**a**), but for K562 ENCODE ChromHMM enhancer states.
**c,** Percentage of enhancers identified in different CRISPR experiments that overlap K562 ENCODE4 proximal or distal enhancer-like signatures (pELS, dELS).
**d,** Like (**c**), but for K562 ENCODE ChromHMM enhancer states.

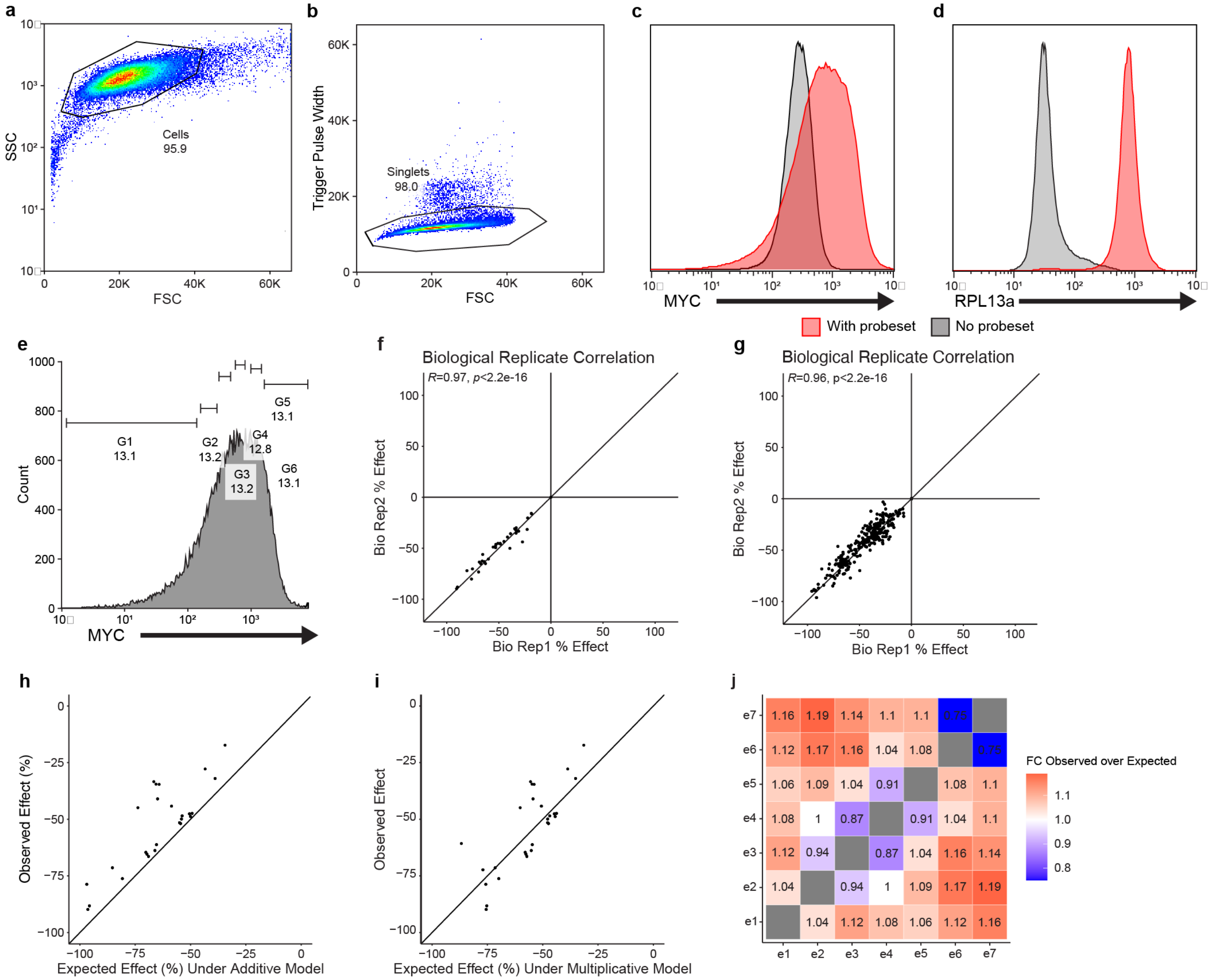


**Extended Data Fig. 7 | CRISPRi-FlowFISH for *MYC* to quantify super-additive enhancer interactions.**

**a,b,** We performed a CRISPRi screen at the *MYC* locus targeting the enhancer elements in pairs. RNA-FISH was used to measure subsequent effects on *MYC* expression. Cells were filter strained prior to flow sorting. Cells were then sorted from debris and from doublet populations.

**c,d,** To measure background cellular fluorescence and confirm that our probe-sets are enriched for signal over noise, a sample was generated without introducing RNA-FISH probes (*gray*). This was compared to the average fluorescent signal from cells with probes (*red*).

**e,** Representative sample of a typical sorting schema; cells were sorted into 6 bins of *MYC* expression.

**f,g,** Effects on *MYC* expression were computed by comparing the non-targeting guide mean expression to the targeting guide mean expression. Biological replicate effect correlations were generated per perturbation (**f**) and biological replicate effect correlations for every guide (**g**) with a significant Pearson R-value > 0.95 in both instances.

**h,** Observed perturbation effects of each element pair vs. the expected effect under an additive model of enhancer-gene regulation, where the expected effect (x-axis) is generated by summing the mean effects of individually perturbing each enhancer.

**i,** In this case we now multiplied the effects of the individual enhancers on gene expression to generate an expected multiplicative model of enhancer-gene regulation versus the observed effect.

**j,** Heatmap of the observed effects divided by the expected effects under a multiplicative model.

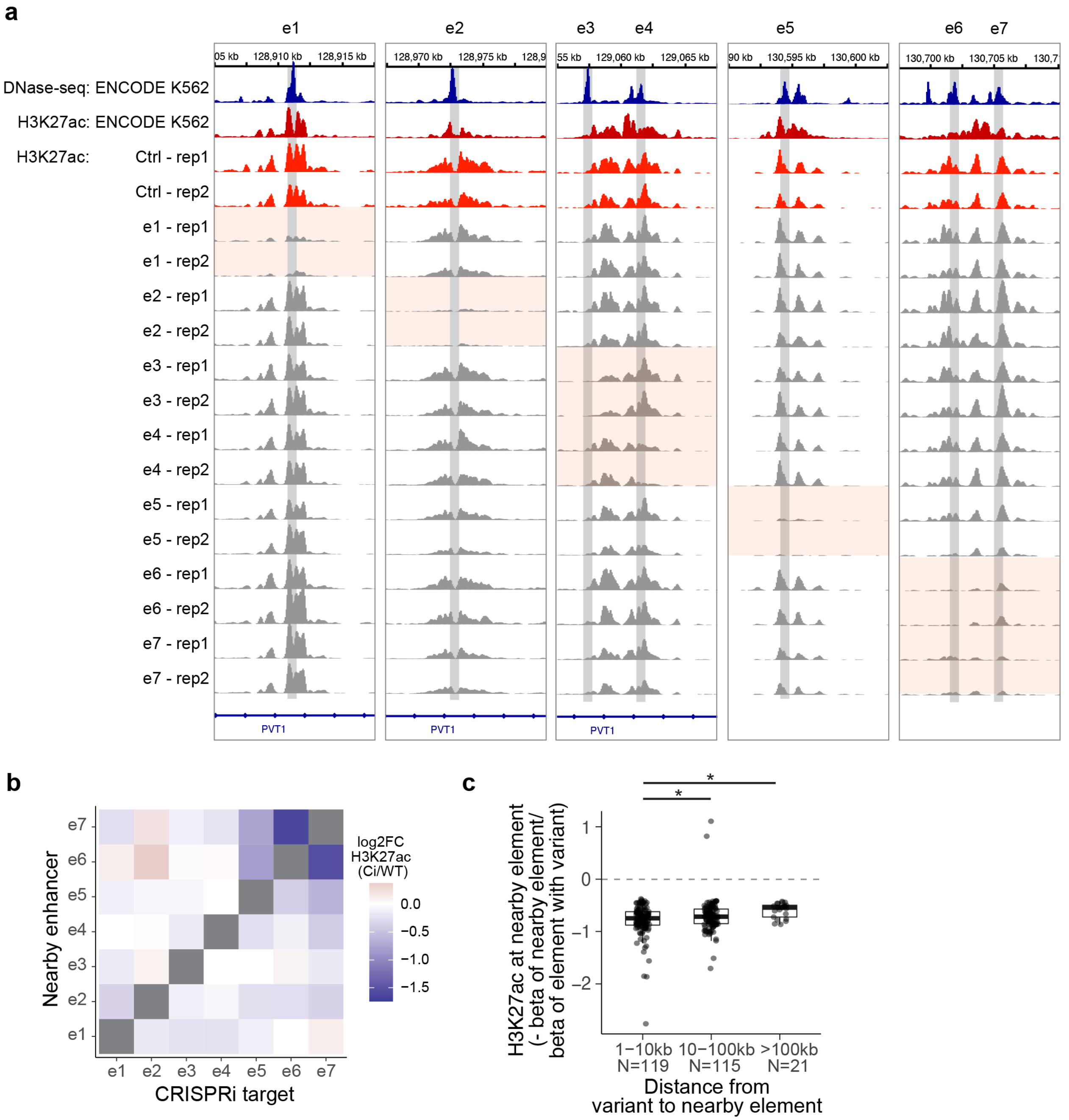


**Extended Data Fig. 8 | Impact of enhancer perturbations on H3K27ac at nearby elements**

**a**, We performed CRISPRi-ChIP-seq at each of 7 *MYC* enhancers (e1-e7): we perturbed each enhancers in K562 using individual CRISPRi gRNAs and performed H3K27ac ChIP-seq to assess the effect on H3K27ac at other nearby enhancers. Signal tracks at top show DNase-seq (blue) and H3K27ac ChIP-seq (dark red) in wild-type ENCODE K562 datasets. Signal tracks below show H3K27ac ChIP-seq in K562 cells perturbed with a negative control gRNA (red) or a gRNA targeting one of the 7 *MYC* enhancers (gray tracks and gray vertical highlights). 2 replicate ChIP-seq experiments are shown for each enhancer. Perturbations to each enhancer successfully deplete H3K27ac at the intended target (orange highlight), as well as lead to quantitative reductions at other nearby elements. Genomic coordinates are shown in hg19.

**b**, Heatmap of log2 fold-change in H3K27ac ChIP-seq signal at each *MYC* enhancer after targeting each other *MYC* enhancer from the experiment described in (**a**).

**c,** Box plots showing effect of chromatin QTLs on a nearby H3K2ac ChIP-seq peak relative to effect on a primary peak stratified by distance between QTL and nearby peak[1]. There is a significant difference in effect size between the QTLs with peaks within 1 and 10 kb and with peaks between 10 and 100 kb (two-sided Wilcoxon rank sum test, p=0.00685), and with peaks greater than 100 kb apart (p = 0.0009174). See Methods, 'Data visualization' section for definition of box plot elements.

#### Supplementary Tables

**Table S1. Collected enhancer-gene predictions**
A list of enhancer-gene linking methods collected as part of this study. This table also contains parameters used for benchmarking analyses.
*Table S1. Collected enhancer-gene predictions.xlsx*

**Table S2. Input datasets for ENCODE-rE2G model and accessions on ENCODE portal**
ENCODE metadata of input data for ENCODE-rE2G model and accession ids for final predictions on the ENCODE portal.
*Table S2. ENCODE-rE2G DNase-only Metadata.xlsx*

**Table S3. Features of enhancers, promoters, and enhancer-promoter pairs**
A list of all molecular features of enhancers, promoters and enhancer-promoter pairs collected as part of this study.
*Table S3. Features of enhancers, promoters, and enhancer-promoter pairs.xlsx*

**Table S4. Input datasets for Average Hi-C**
Metadata and ENCODE accession ids for all Hi-C datasets used to compute average Hi-C interaction frequency across diploid cell types. See Methods 'Generating average Hi-C contact maps' section.
*Table S4. Hi-C files for computing average interaction frequency.xlsx*

**Table S5. Powerlaw values for Hi-C datasets**
Slope (gamma) and intercept (c) values for powerlaw fitted to different Hi-C datasets (see Methods 'Estimating the power-law relationship between 3D contacts and distance' section).
*Table S5. Powerlaw gamma, scale values.xlsx*

**Table S6. Input files for DNase-DNase correlation**
ENCODE accession ids for all DNase-seq files used as input data to compute correlation metrics between DNase-seq signal at cCREs and promoters. See Methods 'Correlation of enhancer-promoter activity across cell types' section.
*Table S6. Input DNase-DNase correlation.xlsx*

**Table S7. Input files for DNase-RNA correlation**
ENCODE accession ids for all DNase-seq and RNA-seq files used as input data to compute correlation metrics between DNase-seq signal at cCREs and RNA-seq expression levels from genes. See Methods 'Correlation of enhancer-promoter activity across cell types' section.
*Table S7. Input DNase-RNA correlation.xlsx*

**Table S8. Input datasets for GraphReg and accessions to predictions on ENCODE portal**
ENCODE accession ids for features and input data that were used as input for the GraphReg model and resulting predictions. See Methods 'GraphReg' section.
*Table S8. GraphReg_LR input data.xlsx*

**Table S9. Metadata for EPIraction**
Metadata for EPIraction predictions in 85 different tissues and cell types, including links to files containing predictions. See Methods 'Epiraction' section.
*Table S9. EPIraction metadata.xlsx*

**Table S10. Input datasets for ENCODE-rE2G$^{Extended}$ model and accessions on ENCODE portal**
ENCODE accession ids or urls for data used as features to compute and apply ENCODE-rE2G$^{Extended}$ in K562 and GM12878. Also includes ENCODE accession ids for final predictions on ENCODE portal.
*Table S10. ENCODE-rE2G extended input data.xlsx*

**Table S11. Input data for baseline predictors and links to predictions on synapse.org**
ENCODE accession ids for files used as input data to compute baseline predictors in 89 biosamples. Also contains urls for resulting baseline predictor files on synapse.org. See Methods 'Generating baseline predictors' section.
*Table S11. Baseline predictors input data and predictions.xlsx*

**Table S12. Performance of predictive model on the CRISPR benchmark**
Performance of predictive models of enhancer-gene regulatory interactions at predicting the experimental results of the combined CRISPR data in the CRISPR benchmarking analyses. Columns show area under the Precision-Recall curve (AUPRC), Precision and threshold values at 70% Recall and 95% confidence intervals for AUPRC and Precision at 70% Recall obtained via bootstrapping (1000 iterations).
*Table S12. CRISPR benchmark performance summary.xlsx*

**Table S13. GWAS benchmarking and guidelines**
Numerical results from benchmarking predictive models of enhancer-gene regulatory interactions against fine-mapped GWAS variants.
*Table S13. GWAS benchmarking and guidelines.xlsx*

**Table S14. GTEx tissue and biosample pairs used in eQTL benchmarking**
GTEx tissues matched to ENCODE biosamples used to benchmark predictive models of enhancer-gene regulatory interactions against fine-mapped eQTL variants.
*Table S14. GTEx tissue and biosample pairs used in eQTL benchmarking.xlsx*

**Table S15. GM12878 enrichment recall table**
Enrichment-Recall values for predictive models of enhancer-gene regulatory interactions benchmarked against fine-mapped eQTL variants in GM12878.
*Table S15. GM12878 enrichment-recall table.xlsx*

**Table S16. Baseline predictors GM12878 enrichment recall table**
Enrichment-Recall values for baseline predictors of enhancer-gene regulatory interactions benchmarked against fine-mapped eQTL variants in GM12878.
*Table S16. Baseline predictors GM12878 enrichment-recall table.xlsx*

**Table S17. Enhancer activity ABC input files**
ENCODE accession ids for files used as input data to compute Activity-By-Contact modes using different chromatin assays to measure enhancer activity. See Methods 'Additional analyses: Assaying enhancer activity' section.
*Table S17. Enhancer activity ABC input files.xlsx*

**Table S18. ENCODE-rE2G summary statistics per biosample**
Summary statistics for binary ENCODE-rE2G predictions of regulatory enhancer-gene interactions 352 biosamples.
*Table S18. ENCODE-rE2G summary statistics per biosample.xlsx*

**Table S19. ENCODE-rE2G summary statistics per gene**
Summary statistics per gene from binary ENCODE-rE2G predictions of regulatory enhancer-gene interactions in 352 biosamples.
*Table S19. ENCODE-rE2G summary statistics per gene.xlsx*

#### Supplementary Text

##### Note S1 | Generation of the integrated CRISPR dataset

We collected and reprocessed 3 previously published CRISPRi enhancer screen datasets from Nasser *et al.*, 2021[2] (mostly CRISPRi FlowFISH), Gasperini *et al.*, 2019[3] (Perturb-seq) and Schraivogel *et al.*, 2020[4] (TAP-seq) (**Fig. S1a**). These different experiments applied different candidate element selection strategies, leading to different characteristics and possible different performance of E-G predictive models (**Fig. S1e**). The CRISPRi FlowFISH experiments in Nasser *et al.*, 2021[2] applied a tiling approach where all DHS sites within five loci were perturbed with a median of 55 guide RNAs, and also incorporated data from other published studies. Gasperini *et al.*, 2019[3] targeted a selection of active enhancers across the genome with 2-4 guide RNAs. Lastly, Schraivogel *et al.*, 2020[4] targeted DHS sites within two genomic regions on chromosomes 8 and 11 and with four guide RNAs each. In an attempt to standardize the analysis of these datasets, we analyzed all experiments on the level of DHS sites, e.g. individual guide RNA perturbations were aggregated per targeted DHS.

To analyze the single-cell RNA-sequencing based datasets from Gasperini *et al.*, 2019 and Schraivogel *et al* 2020, we developed an analysis pipeline centered around performing differential expression tests of perturbed vs. non-perturbed cells for each target enhancer (see Methods section 18, https://github.com/argschwind/ENCODE_CRISPR_data). Depending on the sensitivity of the molecular readout, the number of perturbed cells per targeted element and the expression levels of target genes, CRISPRi screens can have differing statistical power to detect the effect of CRISPRi perturbations on gene expression levels. In practice this can lead to tested element-gene pairs with insufficient statistical power to detect changes in expression of enhancer perturbations. This introduces false negatives, which will negatively impact benchmarking, where E-G pairs are treated as either true positives or true negatives. To filter out any potential false negatives from the collected CRISPR screens, we implemented a simulation-based power analysis that estimates the statistical power of each tested E-G pair to detect specific effect sizes on gene expression levels upon perturbation (e.g. 25% reduction in expression). A similar approach was previously applied in the analysis by Nasser *et al.*, 2021[2]. For effect sizes below 25%, we observed less than 80% statistical power for the majority of tested E-G pairs in both Gasperini *et al.*, 2019[3] and Schraivogel *et al.*, 2020[4] **(Fig. S1b,c)**. For our benchmarking analyses we filtered out any negatives with less than 80% power to detect an effect size of 15% for Gasperini *et al.*, 2019[3] and Schraivogel *et al.*, 2020[4], and 25% for Nasser *et al.*, 2021[2].

In the re-analyzed and power-filtered datasets we observed a total of 472 positive and 9,903 negative E-G pairs (**Fig. S1d),** covering a total of 3,949 candidate elements and 2,121 nearby candidate target genes (**Fig. S1d**). We observed a wide-range of measured effect sizes for detected enhancer-gene links ranging from 1% - 93% reduction of target gene expression levels upon CRISPRi perturbations (**Fig. 1c, Fig. S1f)**, with stronger effects observed for enhancers closer to the target gene TSS. The re-analyzed CRISPR data is available on the ENCODE portal under the accession number ENCSR998YDI and as part of the CRISPR benchmasking pipeline under: https://github.com/EngreitzLab/CRISPR_comparison/blob/main/resources/crispr_data/EPCrisprBenchmark_ensemble_data_GRCh38.tsv.gz

Of note, CRISPR screen experiments do not discriminate between *cis*-acting effects of enhancers and other indirect effects. For example, perturbations of elements regulating 3D contacts (e.g. CTCF binding sites) or *trans*-acting regulatory effects of enhancer-controlled genes can impact downstream gene expression[5]. We therefore cannot exclude that some of the observed regulatory connections are caused by non-enhancer or trans-acting effects, which might bias benchmarking analysis and training models using these data. However, given close proximity to transcription sites for most CRISPR hits, we expect the bulk of these to be real enhancer-gene regulatory connections (**Fig S1.1f**).

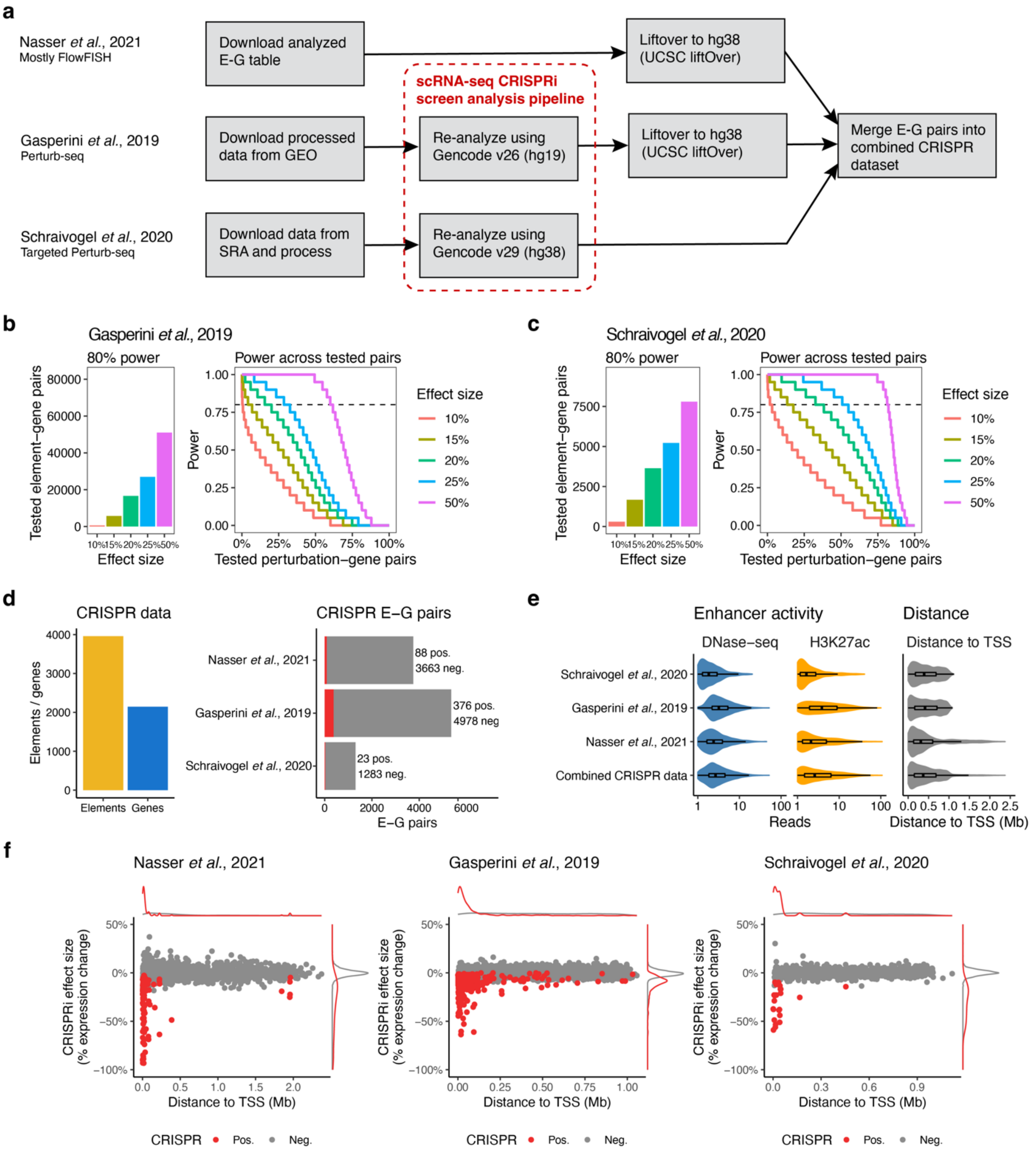


**Figure S1.1 | Generation of the integrated CRISPR dataset**

**a,** Overview of approach to re-analyze and integrate three published CRISPRi screens from Nasser *et al.*, 2021[2], Gasperini *et al.*, 2019[3] and Schraivogel *et al.*, 2020[4].

**b,** Distribution of statistical power to detect specific effect sizes across all experimentally tested element-gene (E-G) pairs and number of E-G pairs with at least 80% power to detect specific effect sizes for the Gasperini *et al.*, 2019 dataset [3].

**c,** Same as b for Schraivogel *et al.*, 2020 [4]

**d,** Number of tested elements and target genes in the combined CRISPR dataset and number of positive and negative E-G pairs per individual dataset.

**e,** Properties of the collected CRISPRi datasets. Shown are activity metrics (DNase-seq and H3K27ac reads overlapping enhancers) for all tested enhancers and distance to TSS distributions for all tested enhancer-gene pairs. See Methods, 'Data visualization' section for definition of box plot elements, outlying points are not shown individually.

**f,** Observed CRISPRi effect sizes as a function of distance to target gene TSS for each of the three datasets.

**Note S2 | Evaluating features of 3D genome organization**

The regulatory functions of an enhancer on a target promoter are thought to be influenced by 3D genome organization, but the extent to which features of 3D contact maps can predict regulatory interactions has been debated[6]. To study such effects, the ENCODE Consortium has measured and defined various features of the 3D genome — including 3D loops (from Hi-C or ChIA-PET), 3D contact frequencies (from Hi-C), contact domains (from Hi-C), and CTCF occupancy (with ChIP-seq) (**Table S2**). Here, we systematically assessed how individual features of 3D genome organization correlated with regulatory effects of enhancers on target genes and contributed to the accuracy of predictive models.

We first characterized the relationship of genomic distance with each of these features, as well as with measured and predicted regulatory enhancer-gene pairs (**Fig. S2.1a-c**). As expected, each feature showed an inverse relationship with distance, with different distributions (**Fig. S2.1a-c**). For example, the 3 types of measured regulatory enhancer-gene relationships each had different decreasing frequencies with distance (median distance = 16.1, 33.2, and 54.9 kb for eQTL, CRISPR, and GWAS enhancer-gene pairs, respectively; 87.1%, 74.1%, and 63.0% of regulatory pairs were located within 100 kb) (**Fig. S2.1a**). The predictive models, including ENCODE-rE2G and ABC, showed similar relationships with distance (**Fig. S2.1c**).

We next assessed the extent to which these features of 3D contact maps, either alone or in combination with other features, contributed to predicting regulatory interactions identified in CRISPR experiments:

Features from measurements of 3D physical interactions, each considered alone, showed poor prediction accuracy (**Fig. S2.1d-e**). For example, assigning enhancers to promoters on the basis of quantitative 3D contact frequency performed better than binary loop or domain calls (recall = 38% at 70% precision, for cell-type specific ENCODE Hi-C at 2 billion reads), but only slightly better than genomic distance (recall = 32% at 70% precision, **Fig. S2.1e**).

Strong 3D contact frequency was neither necessary nor sufficient for enhancer regulation (**Fig. S2.1d**). Among the tested E-G pairs in the top 1% of contact frequencies, only 66 of 102 showed regulatory effects in the CRISPR experiments. The remaining 36 E-G pairs that did not affect gene expression had very low enhancer activity (as estimated from DNase-seq and H3K27ac signals), consistent with these elements being accessible but without strong activating functions (**Fig. S2.1d**). Conversely, while 99% of the tested E-G pairs in the bottom decile of 3D contact frequencies did not have regulatory effects, 7 pairs did. These pairs tended to have very strong activity, consistent with a model where enhancer effects depend on both enhancer activity and 3D contact frequencies.

Accordingly, we next combined different estimates of 3D contact frequency with information about enhancer activity using the Activity-by-Contact framework[5], and found that models using contact frequencies from Hi-C outperformed models that used genomic distance alone. Specifically, we constructed ABC models in which we estimated 3D contact frequency with either: (i) cell-type specific ENCODE Hi-C data (K562, 2 billion reads) at 5 kb resolution, either coverage-normalized (scale-invariant, normalizing for 1D coverage) or unnormalized; (ii) CTCF or RNA Pol II ChIA-PET counts between the element and target gene promoter, or for loops spanning or crossing the E-G pair; (iii) a cell-type-average Hi-C "megamap", in which we averaged contact frequencies from ENCODE Hi-C datasets in 35 tissues to capture cell-type-invariant features of genome organization; or (iv) a power-law function of genomic distance fit to Hi-C data ($1/\text{distance}^{1.03}$). All of these methods performed far better when combined with enhancer activity than they did alone (**Fig. S2.1e-g**). Among these methods, ABC using cell-type-specific ENCODE Hi-C data (5-kb resolution, normalized) performed best (AUPRC = 0.61), cell-type-averaged ENCODE Hi-C (AUPRC = 0.57) and the power-law function of distance (AUPRC = 0.53) (**Fig. S2.1f**). These differences were especially pronounced for E-G pairs located >100 kb apart, where both cell-type specific and cell-type-averaged Hi-C measurements substantially outperformed the power-law (AUPRC = 0.183, 0.145, and 0.0987, respectively; **Fig. S2.1g**). Together, these observations confirm the importance of pairwise enhancer-promoter contact frequencies in predicting enhancer-promoter regulation, and show that cell-type-specific features of 3D contact maps contribute to long-range enhancer regulation beyond a simple function of genomic distance. Notably, the cell-type-averaged ENCODE Hi-C map explained 94% of the variance in the cell-type-specific map (at 5-kb resolution, **Fig. S2.1k**), supporting the use of this generic map for making predictions in cell types in which Hi-C data are not yet available.

Finally, we found that combining pairwise E-G contact frequency with information about CTCF-bound loops that cross or span the enhancer-promoter pair improved regulatory predictions. We computed two quantitative scores based on CTCF ChIA-PET data: the PET counts spanning the E-G pair ("Spanning CTCF loop counts"), and the PET counts crossing the range of the E-G pair (where one PET anchor is inside the E-G pair, and one is outside; "Crossing CTCF loop counts") (**Fig. S2.1h**). Regulatory E-G pairs from CRISPR experiments were 1.9-fold enriched for having at least one spanning CTCF loop count, compared to non-regulatory E-G pairs ($P$ = 0.00000721, **Fig. S2.1i**), and tended to have lower crossing CTCF loop counts (odds ratio = 0.8, $P$ =0.000649). We tested combining these features with the ABC score or with a set of baseline features in logistic regression models, and found that these features led to a modest but significant increase in performance (AUPRC = 0.601 vs 0.625 for ABC alone versus ABC plus spanning and crossing CTCF loop counts, $P_{bootstrap}$ < 0.001; **Fig. S2.1j**). In contrast to these CTCF looping features, incorporating CTCF binding strength or CTCF ChIA-PET counts directly at or between the enhancer or promoter did not improve performance in logistic regression models. Indeed, of the ~500 regulatory E-G pairs in the CRISPR dataset, only 88 (17.3%) had CTCF bound at or within 5 kb of the enhancer, only 49 (9.6%) had CTCF bound at or within 5 kb of the promoter. Together, these observations indicate that, while only a few regulatory enhancer-promoter pairs may be directly mediated by loops with CTCF anchors at the enhancer or promoter, information about CTCF loops spanning or crossing the E-G pair can inform enhancer-gene regulation beyond pairwise enhancer-promoter contact frequencies.

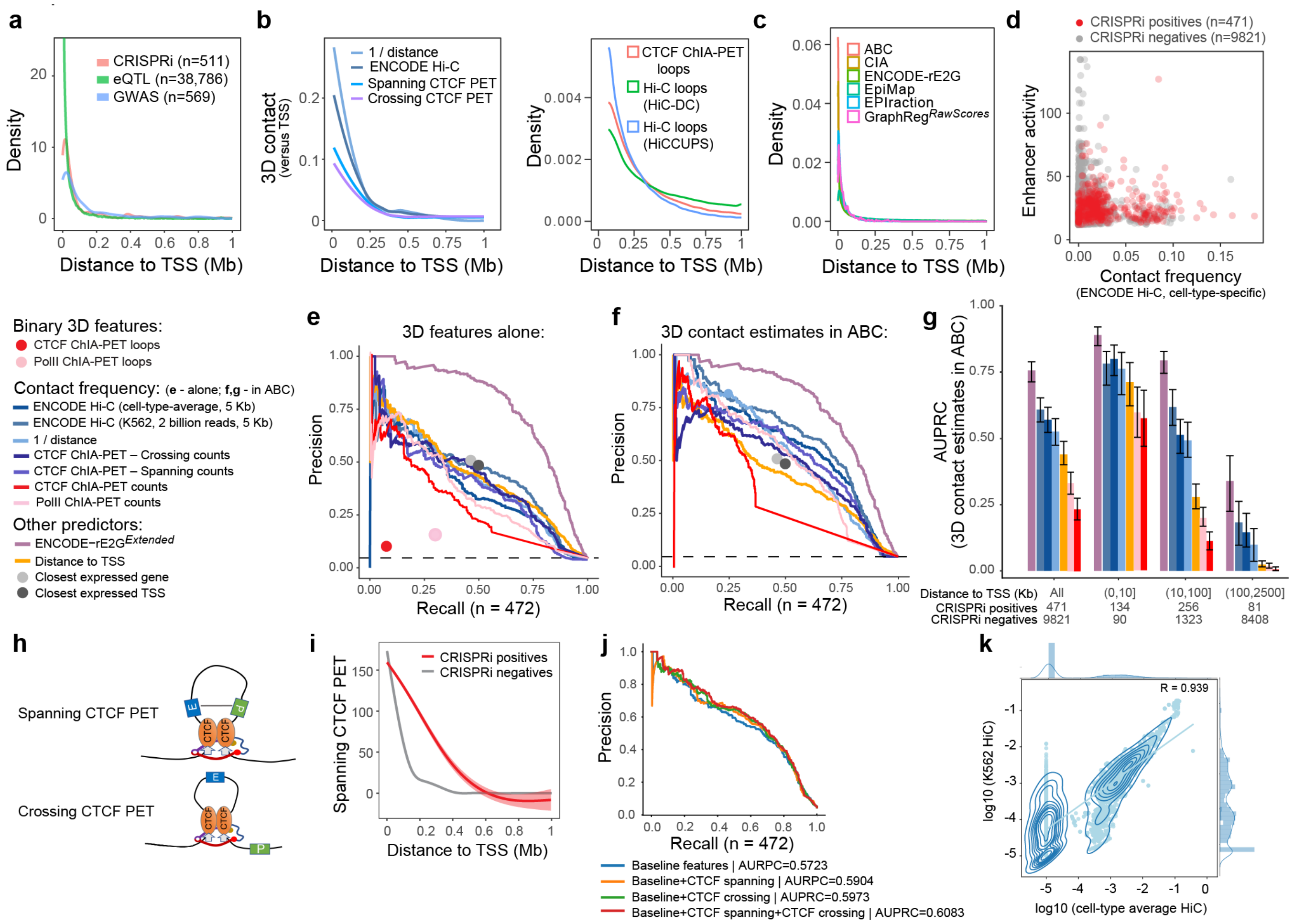


**Fig. S2.1 | Impact of 3D contacts on enhancer regulation**

**a.** As a function of distance, frequency of E-G regulatory interactions from CRISPRi experiments (red), eVariant-eGene pairs (green), and known GWAS variant-gene pairs (blue)

**b.** As a function of distance, frequency of various 3D contact measurements

**c.** As a function of distance, frequency of ABC and ENCODE-rE2G positive predictions

**d.** Relationship between enhancer activity and 3D contact frequency for regulatory (red) and non-regulatory (gray) E-G pairs from the K562 CRISPRi dataset. *X*-axis: Enhancer-promoter contact frequency, from cell-type-specific Hi-C data (5-kb), normalized so that contact frequency at the promoter is 1. *Y*-axis: Enhancer activity as estimated by the ABC model (geometric mean of DNase-seq and H3K27ac ChIP-seq signals).

**e.** Precision-recall plot showing the performance of different 3D features at classifying E-G regulatory interactions from CRISPRi perturbations in K562 cells ($n$ = 10,292 pairs). Lines represent continuous predictors. Dots represent binary predictors. Dotted line: rate of experimental positives. Blue and purple lines represent different measurements or estimates of 3D contact frequencies used directly for prediction.

**f.** Similar to **e**, with blue and purple lines representing ABC models incorporating each 3D contact measurement.

**g.** AUPRC for ABC models using different estimates of 3D contact frequency, binned by the distance from the element to the target gene TSS.

**h.** Illustrating two ways CTCF loop can impact enhancer-promoter (E-P) contacts: a CTCF loop containing E-P pairs facilitates their interaction; a CTCF loop crossing E-P pairs limits their interaction

**i.** Spanning CTCF loop counts from CTCF ChIA-PET as a function of distance for CRISPRi positive (red) and negative (gray) E-G pairs.

**k.** Incorporating PET count of CTCF loops spanning and crossing the E-G pairs improves performance of logistic regression models. “Baseline features” include activity, 3D contact, distance, and others (see Methods).

**l.** Scatterplot showing the relationship between K562 cell type specific Hi-C data and Average Hi-C data across 30 diploid cell types for all E-G pairs in K562.

**Note S3 | Evaluating features related to the correlation between enhancer and promoter activity across cell types**

Many previous approaches to link enhancers to their target genes have involved the assumption that the activities of enhancers and their regulatory target promoters should correlate across cell types or states[7–9]. Such correlations in enhancer and promoter activities depend on the compendium of cell types included in the analysis, and their utility in predicting regulatory interactions has not been systematically compared to other approaches. To test this, we computed 3 metrics: (i) a modified correlation between quantitative DNase-seq signals at enhancer-promoter pairs across 89 biosamples, in which we used generalized least squares to adjust the correlation for particular pair to account for the genome-wide covariance structure among cell types in the analysis ("GLS coefficient", see Methods); (ii) a standard uncorrected DNase-DNase Pearson correlation across the same 89 biosamples; and (iii) a "DNase-RNA GLS coefficient", in which we conducted a similar analysis correlating DNase-seq signals at a distal enhancer with RNA expression level of a nearby gene across 96 matched PolyA+ RNA-seq and DNase-seq datasets.

Overall, the GLS coefficient varied with genomic distance (**Fig. S3.1a**), with higher values at shorter distances similar to the distance distributions of CRISPR-validated enhancers and eQTLs (**Fig. S3.1b**). We compared GLS coefficients to experimental data on enhancer-gene regulatory interactions, and found that regulatory pairs in CRISPRi experiments indeed showed a significantly higher GLS coefficient than distance-matched non-regulatory pairs (mean = 0.3209 vs. 0.1656, respectively; $P = 8.42 \times 10^{-11}$; **Fig. S3.1c**). However, we did not observe similar results for fine-mapped eQTL variants. We overlapped GTEx lymphoblastoid eQTLs with active enhancers in GM12878 cells and compared them with non-significant eQTLs. The GLS coefficient, on average, was lower for GM12878 enhancer-promoter pairs supported by GTEx eQTLs than the ones overlapping the distance-matched non-significant eQTLs (mean = 0.2777 vs. 0.3232, respectively; P = 0.00821; **Fig. S3.1d**). Furthermore, the distributions of GLS coefficients for both CRISPRi enhancers and eQTL variants were highly overlapping with their respective controls (**Fig. S3.1b-d**). As such, the GLS coefficient alone was a poor predictor of regulatory interactions in the CRISPRi dataset (AUPRC = 0.174; **Fig. S3.1e**). The two other correlation-based metrics showed even worse performance (AUPRC = 0.103 and 0.112 for Pearson correlation and RNA GLS coefficients, respectively).

We tested whether the correlations in enhancer-promoter activity across cell types might have independent predictive information from cell-type specific measurements of enhancer-promoter activities or 3D contact frequencies. To do so, we trained logistic regression models in which we supplemented the ABC model or other baseline feature sets with the GLS coefficient, and found that inclusion of the GLS coefficient led to a modest improvement in predictive performance (**Fig. S3.1e**). However, excluding the GLS coefficient from the ENCODE-rE2G$^{Extended}$ model did not significantly reduce performance (**Fig. S3.1e, Fig. S6.3**).

Inspection of the variation of enhancers and promoters across cell types provided a simple explanation for this limited predictive power. We separated the genes from the CRISPRi dataset into two groups: the genes expressed in all 89 biosamples (broadly active promoter) and the ones expressed in several samples (variably active promoters). Similarly, we separated enhancers into 3 groups based on their pattern of activity across cell types: broadly active, active specifically in K562, and inactive (**Fig. S3.1f**). Genes paired with either broadly active or specifically active enhancers showed similar distributions of enhancer-promoter distances (**Fig. S3.1g**). In cases where both the enhancer and promoter were active only in specific cell types, the GLS coefficient performed well. In most cases, however, promoters are active in a much wider array of cell types than the enhancer is active, leading to reduced correlation and worse predictive power (**Fig. S3.1h-k**).

In summary, correlations in enhancer and promoter activity across cell types were insufficient to accurately identify enhancer-gene regulatory interactions, and had only modest/no independent predictive power that could not be captured by features derived from the single specific cell type of interest.

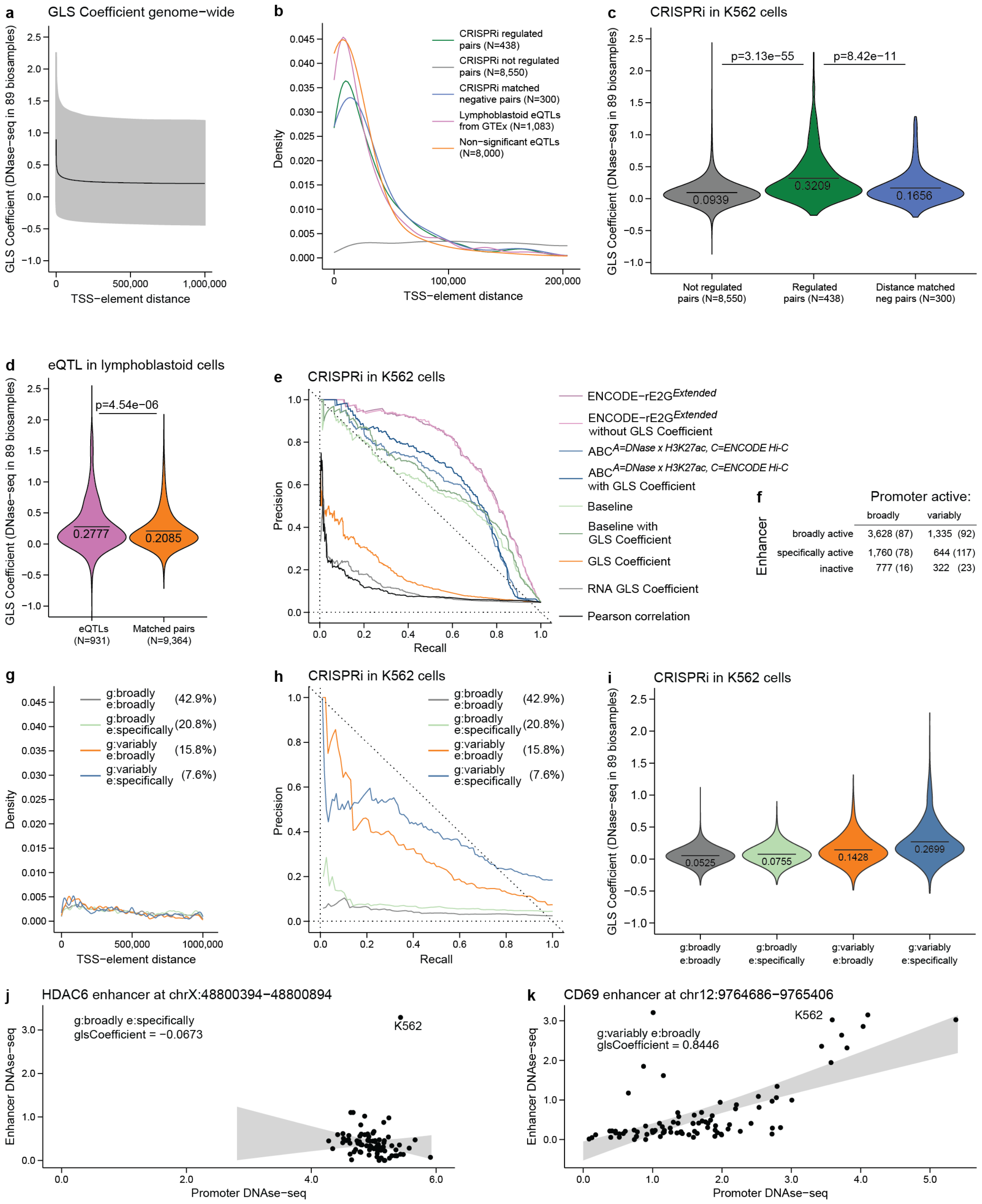


**Fig. S3.1 | Informativeness of correlations in enhancer-promoter activity across cell types.**

**a.** Distance distribution of GLS coefficient, 3.7 million observations were sorted by distance, means and 95% intervals across 10,000 consecutive observations are shown.

**b.** Distance distribution of CRISPRi pairs and GTEx eQTLs.

**c.** GLS coefficient in CRISPRi positive and negative enhancer-promoter pairs.

**d.** GLS coefficient for enhancer-promoter pairs supported by GTEx eQTLs from lymphoblastoid cells.

**e.** GLS coefficient performance predicting CRISPRi positive interactions and its contribution to other models.
**f.** Classification of CRISPRi pairs into different categories based on the amount of cell types corresponding promoter and enhancer are active in, the number in brackets correspond to the amount of CRISPRi positive pairs. **g.** No substantial difference in distance distribution for enhancer-promoter pairs with different spectra of activity.
**h.** GLS coefficient performance predicting CRISPRi positive interactions for different groups of enhancer and promoter activities.
**i** GLS coefficient distribution in corresponding groups.
**j.** Example of the broadly expressed gene regulated by K562-specific enhancer in K562 cells. Biosamples with no active enhancer dominate for GLS coefficient calculation, resulting in non-informative random estimation.
**k.** Example of gene, expressed in several biosamples, regulated by broadly active enhancer. GLS coefficient is able to efficiently predict such interactions.

**Note S4 | Evaluating feature importance in ENCODE-rE2G models**

We used several analysis strategies to evaluate the importance of features in the ENCODE-rE2G and ENCODE-rE2G$^{Extended}$ models.

We performed an initial feature attribution analysis by computing SHAP scores for all used features (**Fig. S4.1, Fig. S4.2**). For ENCODE-rE2G$^{Extended}$, the top feature is enhancer activity measured by EP300 ChIP-seq. The top 10 features contributing to a good performance of ENCODE-rE2G$^{Extended}$ include those from diverse categories, including from ABC (enhancer and promoter activities measured by DNase and H3K27ac), GraphReg (gradient features), genomic features (distance to TSS, number and summation of nearby enhancers), and CTCF (number of CTCF PET counts crossing E-P pairs) (**Fig. S4.1**). For ENCODE-rE2G, the top contributing feature is DNase signal at the enhancer, and among the top 10 features there are DNase-only ABC scores (which are derived from average Hi-C and DNase used for enhancer activity), genomic features (distance to TSS), and promoter class (ubiquitously expressed genes) (**Fig. S4.2**).

Because many individual features are correlated with one another, we performed a groupwise ablation analysis of the ENCODE-rE2G and ENCODE-rE2G$^{Extended}$ models. To this end, we defined categories of related features (some features are assigned to multiple categories) and computed logistic regression models leaving out each category of features. We calculated the difference between the performance (AUPRC and precision at the recall value of 0.7) of the ablated model and the full-featured model to define the delta performance (Delta_AUPRC = AUPRC_ablated - AUPRC_full, and Delta_Precision = Precision_ablated - Precision_full). To assess the significance of differences in performance, we performed bootstrap sampling of the E-G pairs (1000 samples, with replacement). In order to see the effects of each feature group as a function of E-G distance, we also performed similar analyses after stratifying E-G pairs to three distance ranges [0, 10kb), [10kb, 100kb), and [100kb, 2.5Mb] (**Fig. 5a, Fig. S4.3, Fig. S4.4**).

For both ENCODE-rE2G and ENCODE-rE2G$^{Extended}$, the two most important categories were 3D Contacts/Distance and Enhancer Activity (**Fig. S4.3, Fig. S4.4**) — as expected based on the good performance of previous models like ABC, which uses only these two components. Notably, ablation of the “Contact” category, which removes all features derived from Hi-C and ChIA-PET but preserves features related to 1D genomic distance, significantly reduced performance. This supports observations that 3D contact measurements include information about enhancer-promoter regulation that is not captured by genomic distance alone (see also **Note S2**). Interestingly, other categories of features beyond those included in ABC had smaller but significant effects on model performance, including promoter class and nearby enhancer activity for both ENCODE-rE2G and ENCODE-rE2G$^{Extended}$ (see also **Note S6**, **Fig. 5**, **Fig. 6**).

The importance of feature categories varied across bins of genomic distances in the ablation analysis (**Fig. S4.3**, **S4.4**). For example, while ablation of the “enhancer activity” feature category significantly affects model performance at all distance ranges, categories of features involving 3D contacts were only important in the longer distances ranges (>10 kb), as expected.

Finally, we performed forward sequential feature selection on both models (**Fig S4.5)**. For models using features from ENCODE-rE2G, performance reached a local maximum in AUPRC with 8 of the 13 features, although it achieved highest performance including all features (**Fig. S4.5a**). Interestingly, the first three features selected were the ABC$^{A=DNase,\ C=Average\ Hi\text{-}C}$ score, whether the regulated gene was ubiquitously expressed, and the number of DNase peaks within 5 kb of the enhancer, consistent with the importance of these features in other analyses (**Fig. 5c-e**). For ENCODE-rE2G$^{Extended}$, the maximum AUPRC was achieved with just 16 of the 47 features (**Fig. S4.5b**). These 16 features also spanned many categories, including ABC score, promoter class, correlation, PET counts, activity at nearby enhancers, promoter class, and GraphReg gradients.

Altogether, these results show that multiple categories of features are complementary and important for optimal performance of both the ENCODE-rE2G and ENCODE-rE2G$^{Extended}$ models.

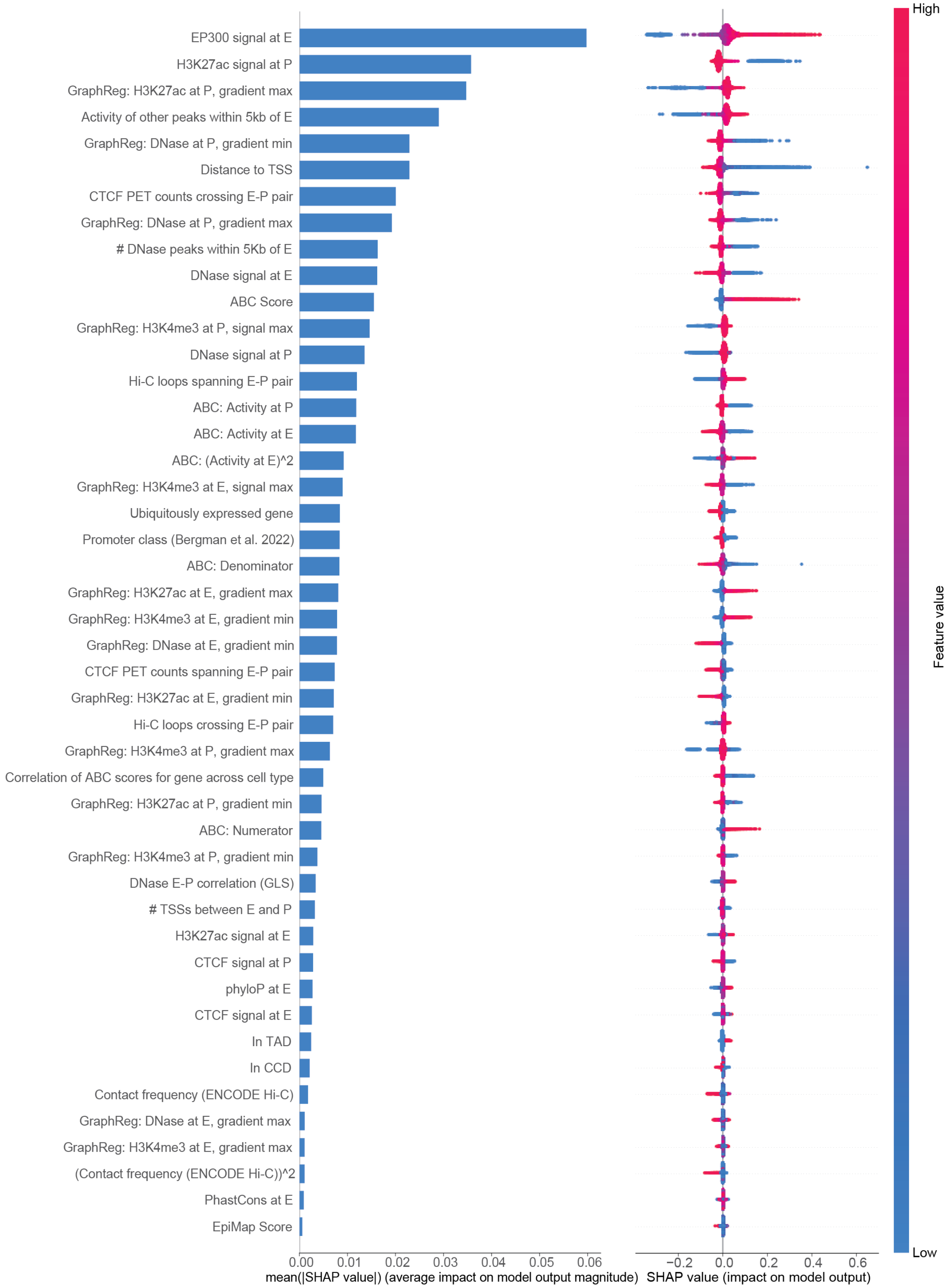


**Fig. S4.1 | SHAP scores of ENCODE-rE2G$^{Extended}$ model**

SHAP scores of all 46 features used by ENCODE-rE2G$^{Extended}$ model in a decreasing order. Left bar plot shows the mean of absolute SHAP scores for all the predicted EG pairs. Right dot plot shows the SHAP scores of each predicted EG pair color coded by the feature values.

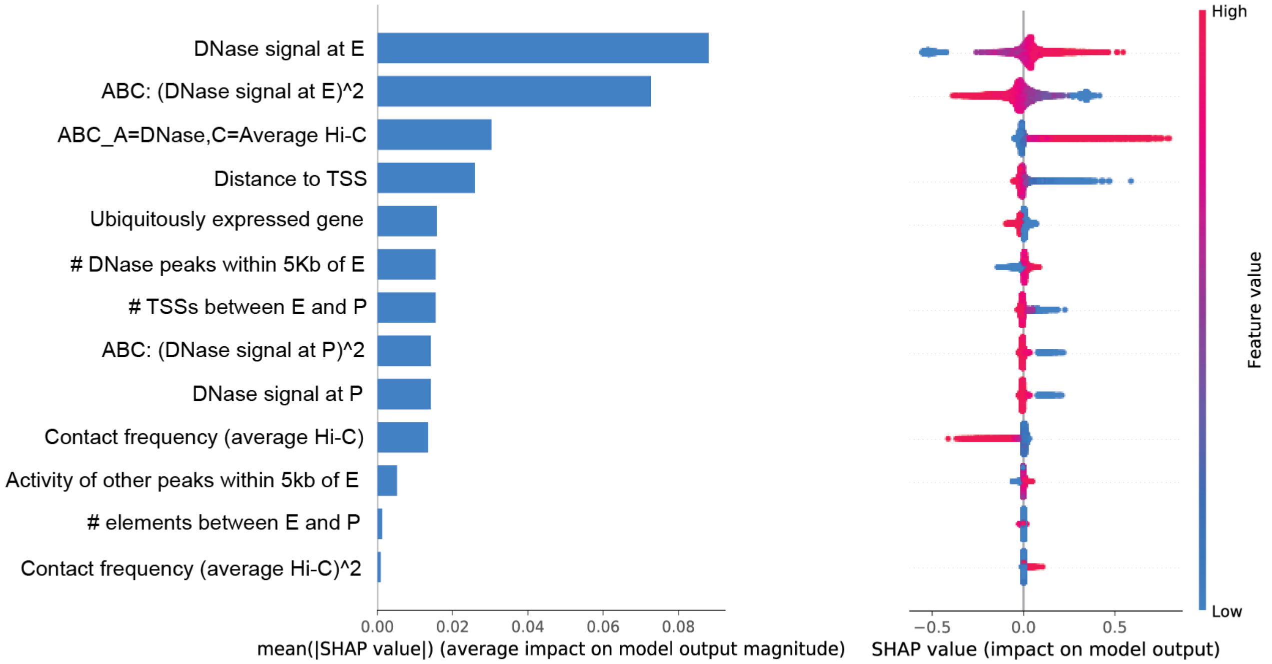


**Fig. S4.2 | SHAP scores of ENCODE-rE2G model**

SHAP scores of all 13 features used by ENCODE-rE2G model in a decreasing order. Left bar plot shows the mean of absolute SHAP scores for all the predicted EG pairs. Right dot plot shows the SHAP scores of each predicted EG pair color coded by the feature values.

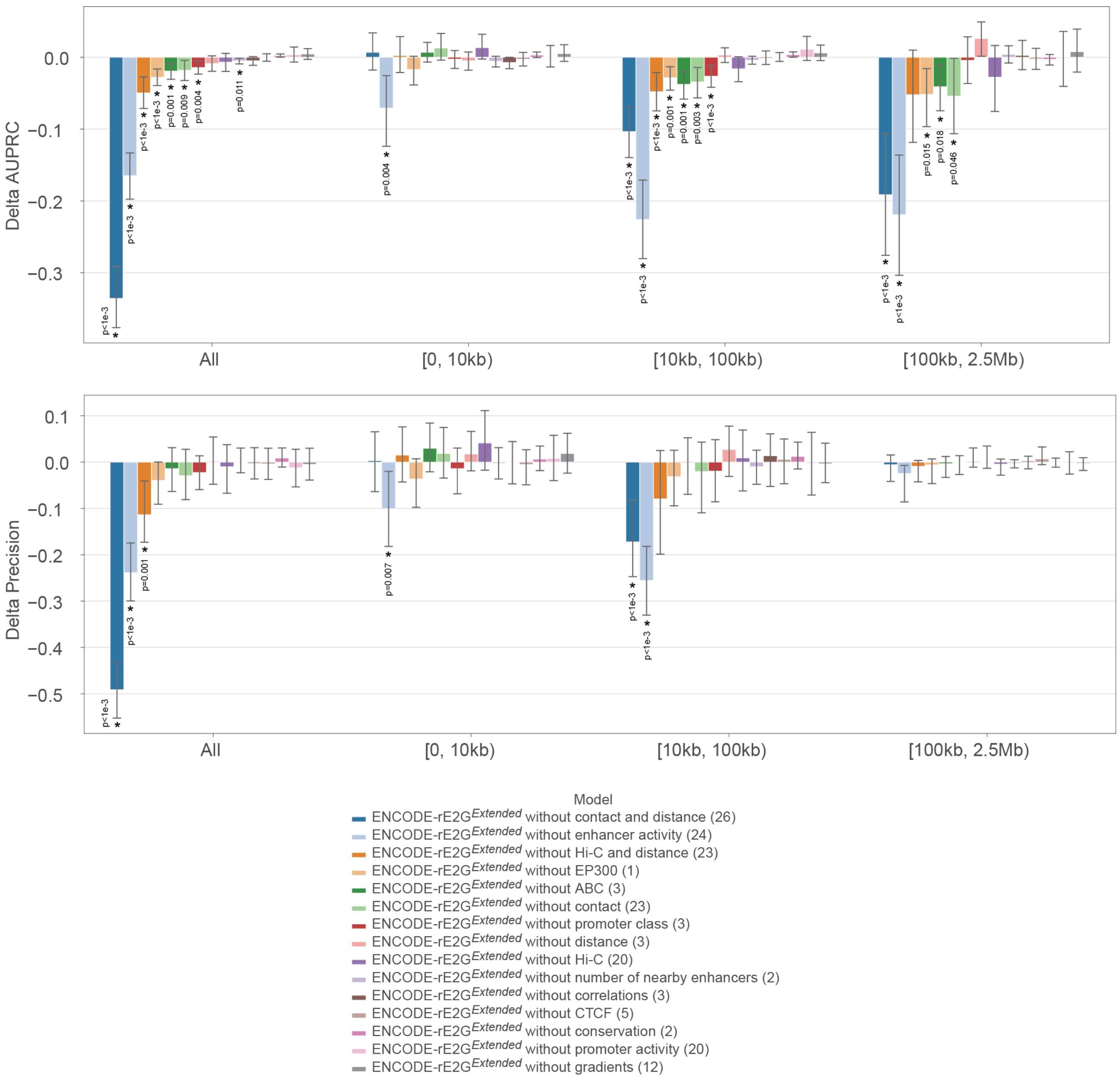


**Fig. S4.3 | Performance of ENCODE-rE2G$^{Extended}$ after removing categories of features.**
Effect of different feature categories on EG prediction task in ENCODE-rE2G$^{Extended}$ model. Each feature category is removed from the feature set used by ENCODE-rE2G$^{Extended}$ to see the amount of performance reduction. The numbers inside the parentheses show the numbers of the deleted features. Bar plots of delta AUPRC and delta precision (at recall value 0.7) in distance-stratified ranges. Bar plots and error bars show the mean of delta (AUPRC and precision) and 95% confidence intervals when randomly subsampling the EG pairs with replacement (bootstrapping) 1000 times.

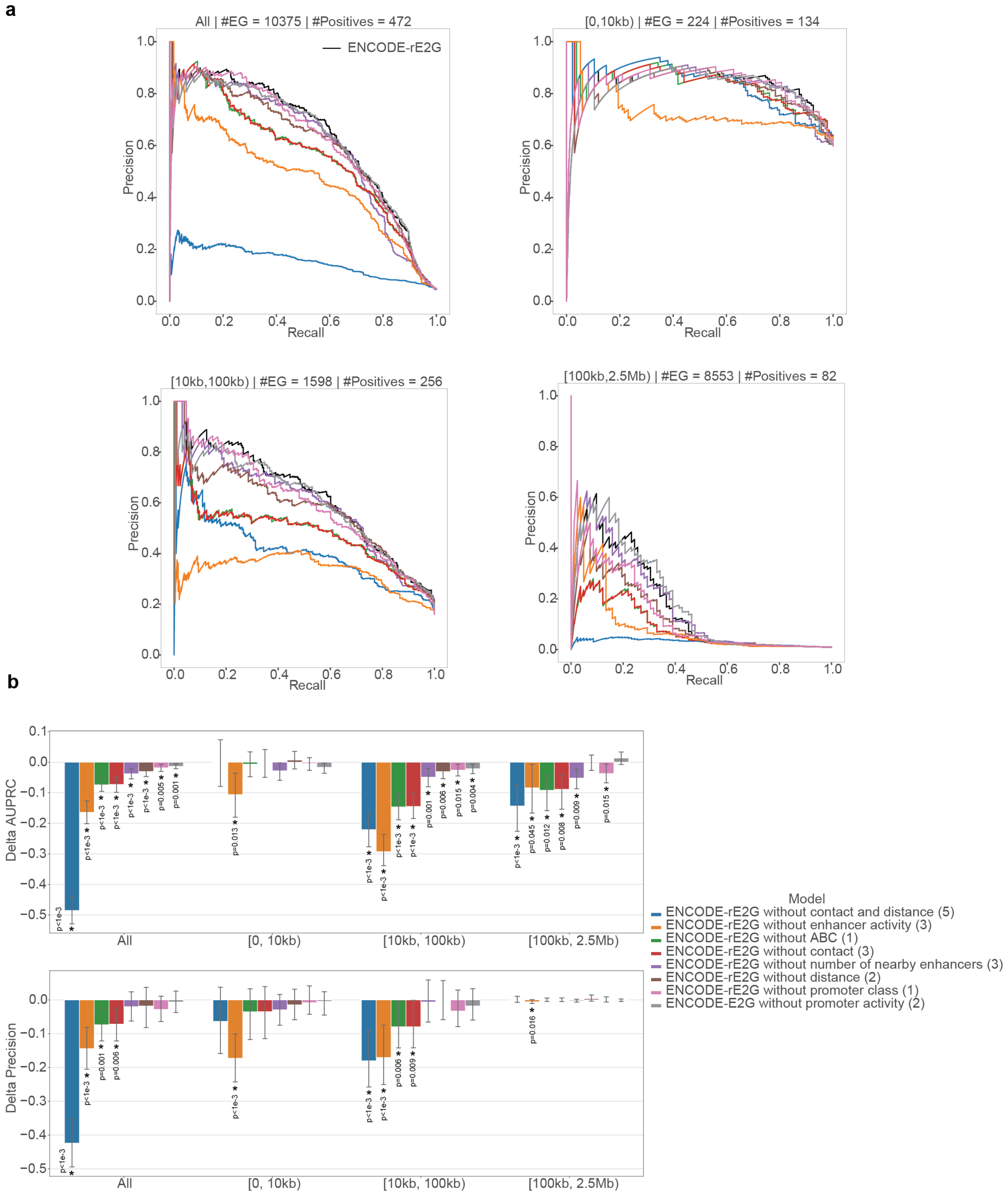


**Fig. S4.4 | Performance of ENCODE-rE2G after removing categories of features.**
Effect of different feature categories on EG prediction task in ENCODE-rE2G model. Each feature category is removed from the feature set used by ENCODE-rE2G to see the amount of performance reduction. The numbers inside the parentheses show the numbers of the deleted features. **a,** Precision-recall curves of ENCODE-rE2G and ablated models in all and distance-stratified ranges. **b,** Bar plots of delta AUPRC and delta precision (at recall value 0.7) in all and distance-stratified ranges. Bar plots and error bars show the mean of delta (AUPRC and precision) and 95% confidence intervals when randomly subsampling the EG pairs with replacement (bootstrapping) 1000 times.

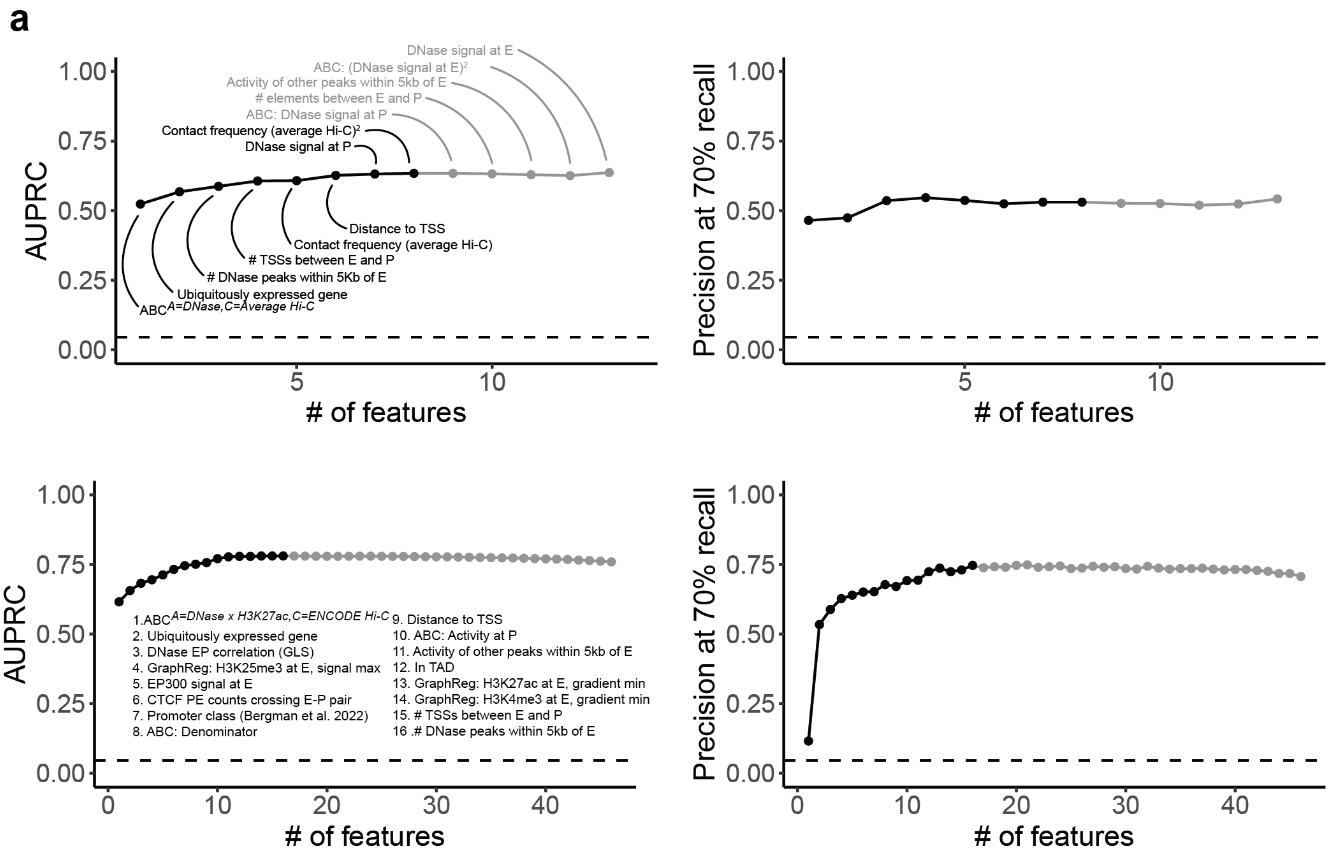


**Fig. S4.5 | Sequential feature selection on ENCODE-rE2G and ENCODE-rE2G$^{Extended}$.**
Features were sequentially added to each model based on maximum increase in AUPRC by adding a given feature. **a.** AUPRC of ENCODE-rE2G reached a local maximum at 8 features, after which performance did not significantly increase until reaching the maximum AUPRC with all 13 features. Precision similarly plateaued after 8 features. b. AUPRC of ENCODE-rE2G$^{Extended}$ reached its absolute maximum with the 16 listed features. Precision with additional features also did not increase significantly.

#### Note S5 | Evaluating GraphReg gradient features in ENCODE-rE2G$^{Extended}$

We recently developed GraphReg, a graph neural network approach that involves predicting RNA expression from measurements of enhancer and promoter activity (DHS, H3K27ac, and H3K4me3) and information about element-gene physical interactions from Hi-C loop calls [10]. In this study, we recomputed GraphReg using ENCODE data and used the predictions in several ways. In an initial approach, we used the raw element-gene weights from the GraphReg model as a predictor of enhancer-gene regulatory interactions, and found that this score performed relatively poorly (GraphReg$^{Raw\ Scores}$ in **Fig. 2a**). As a second approach, we incorporated intermediate features from the GraphReg model — specifically, gradients describing how changing enhancer or promoter activity (DHS, H3K27ac, or H3K4me3 signals) should affect gene expression — into the ENCODE-rE2G$^{Extended}$ model and other logistic regression models.

In this second analysis, we first trained a logistic regression model incorporating only features derived from the GraphReg model (including estimates of enhancer and promoter activity and gradients of how changing enhancer or promoter activity) plus genomic distance ("GraphReg$^{LR}$", see Methods). We then ablated this model by removing all the 12 gradient features, of which 6 corresponds to each enhancer and 6 to each promoter (see Methods). We stratified all the 10,375 E-G pairs in the ensemble CRISPRi dataset based on the distance to three non-overlapping distance-stratified ranges: [0, 10kb), [10kb, 100kb), and [100kb, 2.5Mb). **Fig. S5.1** shows the Precision-Recall (PR) curves, area under the PR curves (AUPRC), and the precision values at the recall value of 0.7 in the entire dataset and the distance-stratified ranges. The first observation is that the gradient features help the performance of the GraphReg$^{LR}$ model. At all distances, removing the gradient features leads to a performance reduction in AUPRC (0.6716 to 0.6629) and precision (0.5651 to 0.5272), though the reductions are not significant based on bootstrapping. The second observation is that the gradient features help significantly in prediction of the long-range enhancers and are of no importance for the short-range enhancers. If we look at the performance of the GraphReg$^{LR}$ model in the two categories [0, 10kb) and [10kb, 100kb), we can see that there is not a performance drop in AUPRC and precision by removing the gradients in the short-range category [0, 10kb); however, in the long-range category [10kb, 100kb), there is a significant drop (16.2%) in precision from 0.6172 to 0.5173 in the GraphReg$^{LR}$ model by removing these gradient features ($P_{Bootstrap}$ = 0.012) (**Fig. S5.1b)**. Furthermore, in the very long-range category [100kb, 2.5Mb), there is a significant drop (40.2%) in AUPRC from 0.1750 to 0.1047 in the GraphReg$^{LR}$ model by removing these gradient features ($P_{Bootstrap}$ = 0.001) (**Fig. S5.1b)**.

To illustrate how gradients can help enhancer predictions, we show the gene locus of *RAB7A* and a candidate enhancer from CRISPRi dataset about 45kb upstream of the gene (**Fig. S5.2b**). There is an interaction between the promoter of this gene and the candidate enhancer based on Hi-C data. Both ABC (score=0.1203) and GraphReg$^{LR}$ model without gradients (score=0.1744) predict a high score for this E-G pair. However, the GraphReg$^{LR}$ model predicts a lower score of 0.0431 for that E-G pair. The reason is that since the graph used in the GraphReg model does not have any E-G interaction between *RAB7A* and the candidate enhancer, the gradients are zero and the predicted score is smaller, showing correctly that this is not a true enhancer of the gene.

We note that, while in the GraphReg$^{LR}$ model the gradients have a significant effect on performance, ablating the GraphReg gradients from the ENCODE-rE2G$^{Extended}$ model did not significantly affect performance (**Fig. S5.2a** and **Fig. S4.3**). This is likely because the ENCODE-rE2G$^{Extended}$ model includes other correlated features such as 3D contact information and the ABC score that are correlated with the GraphReg gradient features (**Extended Data Fig. 1**).

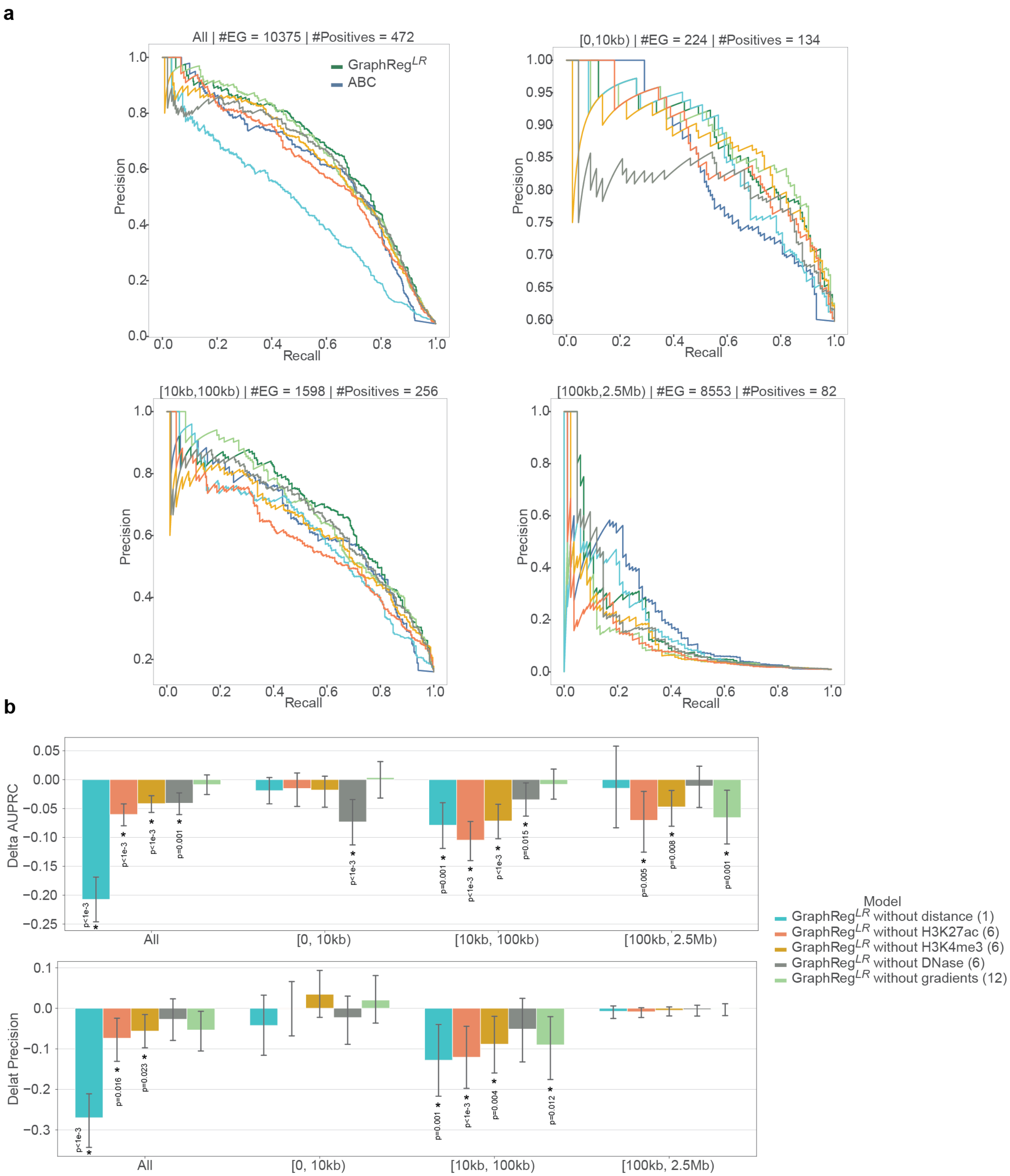


**Fig. S5.1 | GraphReg Ablation in a simplified logistic regression model.**
Effect of different feature categories on EG prediction task in the GraphReg$^{LR}$ model, which includes genomic distance, estimates of enhancer and promoter activity, and 12 gradient features from the GraphReg model. Each feature category is removed from the feature set used by GraphReg$^{LR}$ to see the amount of performance degradation. The numbers inside the parentheses show the number of the deleted features. **a,** Precision-Recall curves show the EG prediction performance of the ABC, the full GraphReg$^{LR}$, and the ablated GraphReg$^{LR}$ models in the ensemble CRISPRi dataset, all and stratified by non-overlapping distance ranges [0, 10kb), [10kb, 100kb), and [100kb, 2.5Mb). **b,** Bar plots of delta AUPRC and delta precision (at recall value 0.7) in all and distance-stratified ranges. Bar plots and error bars show the mean of delta (AUPRC and precision) and 95% confidence intervals when randomly subsampling the EG pairs with replacement (bootstrapping) 1000 times. See Methods, 'CRISPRi benchmark' section for description of statistical test.

**a**

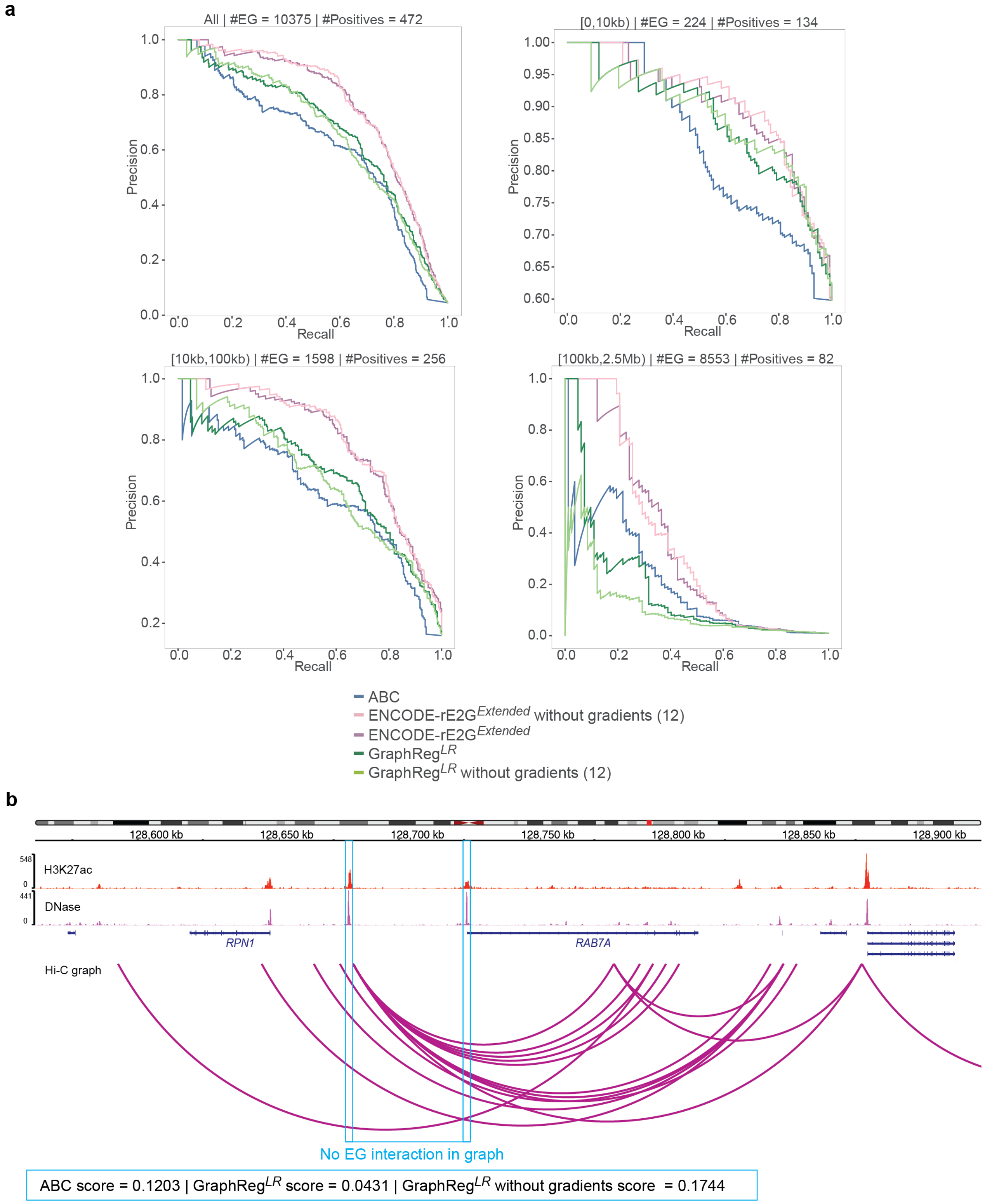


**Fig. S5.2 | Effect of GraphReg gradient features on enhancer prediction.**

**a,** Precision-Recall curve shows the enhancer prediction performance of ABC, ENCODE-rE2G$^{Extended}$, and GraphReg$^{LR}$ models with and without GraphReg's gradient features in the full CRISPRi dataset.

**b,** An example of a candidate enhancer for the gene *RAB7A* about 45kb upstream of its TSS from the Nasser *et al.*, 2021[2] CRISPR dataset which is not a positive (regulating) enhancer. Both ABC and GraphReg$^{LR}$ without gradient predict it as a positive enhancer (false positive) with the scores of 0.1203 and 0.1724, respectively, while GraphReg$^{LR}$ model (which has the gradient features) predicts it correctly as a negative enhancer with the score of 0.0413. This is because of zero gradients for that enhancer as a result of no EG interaction in the graph used by the GraphReg model.

**Note S6 | Evaluating differences between ubiquitously and specifically expressed genes**

Recent studies suggest that enhancer-promoter regulation in the genome may be influenced by differences in the architecture of different promoters, and in particular that ubiquitously expressed (housekeeping) genes might be less sensitive to distal enhancers[11,12]. We evidence to support this effect across our dataset:

We first classified promoters via three metrics (see Methods). (1) The enhancer correlation score - in which we give genes a score based on how consistent their enhancers are across cell types, as calculated by correlating the ABC scores of enhancers across our previous atlas[2]. (2) Promoter class - in which, as previously defined[11], "P1" promoters have weaker intrinsic activity and are more strongly regulated by enhancers and "P2" promoters are intrinsically stronger, less sensitive to enhancers and more ubiquitously expressed across cell types. (3) Expression uniformity- in which we stratify genes based on the ubiquitousness and uniformity of their expression across cell types (see Methods).

We found that for each of these three metrics, genes in the more constitutive gene categories (those with enhancer correlation scores in the top 25% of genes, genes predicted to have P2 promoters, and ubiquitously and uniformly expressed genes) were less likely to be linked to eQTLs in predicted enhancers (**Fig S6.1a**) and predicted enhancers for these genes were less likely to be true regulators in CRISPRi datasets (**Fig S6.1b**).

We tested whether promoter class could be used to improve our predictive models. We found that when we add each of these features to a logistic regression model with just the ABC Score all three lead to significant increases in the AUPRC relative to ABC Score alone (**Fig S76.1c-d**). P2 and ubiquitous expression in particular significantly improve upon the precision of the models (**Fig S6.1e**) and reduce the number of false positive predictions **(Fig S6.1f)**. Furthermore, ubiquitous expression and P2 promoter class are both shown to be predictive in our extended models ranking respectively 19 and 20 out of 47 features in the ENCODE-rE2G$^{Extended}$ model, and with ubiquitous expression ranking 5th out of 13 features in the ENCODE-rE2G model (**Fig. S4.1-2)**. Thus, incorporating information about promoter class can improve predictions of enhancer-gene regulation.

Together, these observations are consistent with a model in which sequence-intrinsic features of the promoters of ubiquitously expressed genes make them insensitive to distal enhancers.

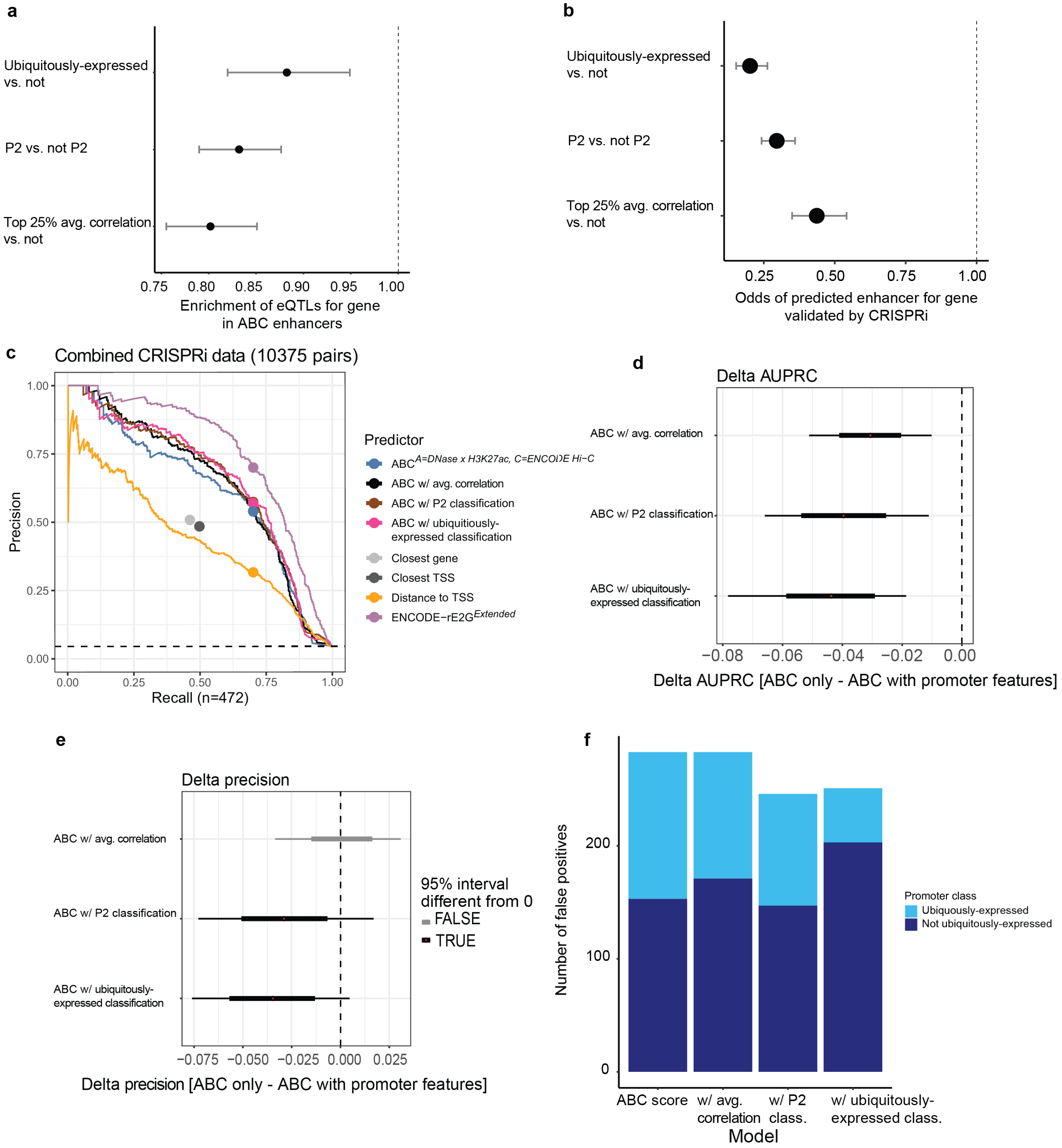


**Fig. S6.1 | Differences in distal regulation between gene/promoter classes**

**a.** Likelihood of eQTL falling within an ABC predicted enhancer element for genes across 3 promoter class features. Ubiquitously expressed genes, P2 promoter class, and genes with enhancer correlation scores in the top 25% of scores are respectively 88.2% (p = 0.006, Fisher's Exact Test) , 83.2% (p = 2.18 * $10^{-12}$, Fisher's Exact Test), 80.2% (p = 1.18* $10^{-13}$, Fisher's Exact Test) as likely as their counterparts to have an eQTL for that gene in a predicted ABC enhancer. The error bars denote 95% confidence intervals.

**b.** Likelihood of a predicted enhancer for a gene being validated by CRISPRi genes across 3 promoter class features. Ubiquitously expressed genes, P2 promoter class, and genes with enhancer correlation scores in the top 25% of scores are respectively 20.2% (p = 6.20 x $10^{-45}$, Fisher's Exact Test) , 29.6% (p = 1.15 x $10^{-37}$, Fisher's Exact Test), 43.7% (p = 7.58 x $10^{-16}$, Fisher's Exact Test) as likely as their counterparts to have an eQTL for that gene in a predicted ABC enhancer. The error bars denote 95% confidence intervals.

**c.** Precision recall curves showing that the addition of the promoter class features improve upon performance of the ABC model alone (blue).
**d.** Delta AUPRC for ABC score alone vs ABC with ubiquitously expressed gene information (delta AUPRC = -0.044, p = 0.0001), P2 promoter class information (delta AUPRC = -0.040, p = 0.0001), enhancer correlation information (delta AUPRC = -0.031, p = 0.0001). Error bars represent 95% range of AUPRC values inferred via bootstrap (10000 iterations).
**e.** Delta precision at 70% recall for ABC score alone vs ABC with ubiquitously expressed gene information (delta precision = -0.029, p = 0.009), P2 promoter class information (delta precision = -0.035, p = 0.0017), enhancer correlation information (delta precision = 0.0008, p = 0.932). Error bars represent 95% range of AUPRC values inferred via bootstrap (10000 iterations).
**f.** Number of false positives for ABC alone compared with ABC with each of the promoter class features for non-ubiquitously expressed genes vs ubiquitously expressed genes. Total false positives: ABC only - 283, ABC with ubiquitous expression information - 251, ABC with P2 information- 246, ABC with enhancer correlation information - 283.

**Note S7 | Using ENCODE-rE2G maps to interpret GWAS signals**

We assessed how to interpret ENCODE-rE2G predictions when applied to link noncoding variants associated with common, complex diseases to target genes and cell types. To this end, we analyzed fine-mapped distal noncoding variants from GWAS studies for 94 traits from the UK Biobank (PIP > 0.1, *N*=29.1K) and expression QTLs for 49 tissues from GTEx (PIP > 0.1 for >=1 gene, *N*=247K). To assess enrichment of these variants in enhancers identified by ENCODE-rE2G, we compared these fine-mapped variants to a control set: all distal noncoding variants detected in the 1000 Genomes project (*N*=9.2 million).

In total, we identified that 46% of fine-mapped GWAS variants and 41% of eQTL variants overlapped an ENCODE-rE2G enhancer in one or more of the 352 biosamples, compared to only 19% of control variants (2.3 and 2.1-fold enrichment, respectively) (**Fig. S7.1a**). In comparison, repeating this for all DNase peaks, 74% of GWAS variants and 69% of eQTL variants overlapped with DNase across all biosamples, compared to 55% of control variants (1.3 and 1.2-fold enrichment, respectively). Blood-related traits showed a 22% higher fraction of overlap with ENCODE-rE2G predictions than an average trait, while brain-related traits showed 39% lower fraction of overlap with ENCODE-rE2G predictions (**Fig. S7.1a**). This can be attributed to both (i) poor representation of brain-related cell types compared to immune cell types among the ENCODE-rE2G biosamples, and (ii) high polygenicity of brain-related traits[13].

Our GWAS analyses expectedly showed that variants were enriched for overlapping ENCODE-rE2G enhancers in expected tissues (**Fig. 4c**). However, in some cases, we also observed significant enrichment for variants for a given trait in enhancers in other biosamples not likely to be relevant, presumably due to sharing of enhancers across multiple cell types or tissues. For example, fine-mapped variants for atrial fibrillation trait were most strongly enriched in skeletal muscle, instead of the likely causal cardiac myoblasts, which was ranked 3rd based on enrichment; this may be attributed to 52% sharing of enhancer variants in cardiac myoblasts with skeletal muscle. At the level of individual fine-mapped variants, we observed similar trends: For any given variant that overlapped an enhancer in 1 biosample, that same variant overlapped enhancers in an average of 26 other biosamples providing multiple candidate biosamples in which the variant might act. At the same time, 21% of variants showed strong cell-state specificity, in that they were identified in only one biosample, often excluding other related biosamples.

Next, towards guiding the design of future data collection, we considered the marginal value of adding biosamples to the compendium. For each trait, we examined the fraction of prioritized fine-mapped variants that were uniquely captured by biosample with the greatest genome-wide enrichment for that trait — *i.e.*, the fraction of variants that would have been missed if one did not have this biosample in the dataset. This ranged from 0-14% across traits (**Fig. S7.1b**). For example, only 1.1% of fine-mapped variants that overlap an ENCODE-rE2G enhancer for RBC count (5 variants) were uniquely identified by hematopoietic progenitor cells, and the remaining 451 were included in enhancers in other biosamples. For the Lymphocyte count GWAS trait, removing top prioritized T-cell biosample (T helper cells) and all T-cell biosamples resulted in a drop of 1 (0.2%) and 16 variants (3.4%) captured respectively. We conclude that putatively disease-critical cell types tend to show the highest global enrichment of fine-mapped variants while for the disease, a large fraction of these fine-mapped variants are mapped by ENCODE-rE2G in other likely related biosamples; as a result, adding cell types closely related to biosamples already included in this ENCODE-rE2G resource may yield only limited additional discoveries with respect to predicting the target gene of noncoding variants.

As a further validation of biosample-specificity of traits, we examined the extent of overlap of GTEx eQTLs for 28 tissues in ENCODE-rE2G maps for 28 groups of biosamples matched to GTEx tissues (**Table S13**). For 43% tissues, the GTEx eQTLs in a tissue showed the highest enrichment in ENCODE-rE2G for the matched set of biosamples; for 68% cases, the matched set of biosamples were among the top 3 most enriched (**Fig. S7.1c**). For example we observed the highest enrichment of fine-mapped eQTLs from GTEx cerebellum and cerebellar hemisphere in ENCODE-rE2G predictions of cerebellum-related biosamples (**Fig. S7.1c**). These results highlight that using ENCODE-rE2G maps to identify candidate causal cell types for variants causally associated with diseases and molecular phenotypes should consider which biosamples are globally enriched for disease variants and consider sharing of enhancers across related cell types. We also expect that identification of causal cell types will likely benefit from additional information about the cell-type specificity of the effects of the variant on enhancer activity (Abramov et al 2021 *Nat Comm*, Vierstra et al 2020 *Nature*), which are not captured by ENCODE-rE2G.

Finally, we examined the degree of redundancy in the ENCODE-rE2G variant-to-gene linking predictions across biosamples by focusing our analyses to a subset of top globally enriched biosamples for a trait. Genes linked to GWAS fine-mapped variants for a trait in the top 10 enriched biosamples showed on average 19% higher PoPS prioritization score for the trait compared to those linked in all biosamples. Of these, 37 of 94 traits showed significantly higher (FDR < 10%) average PoPS prioritization across linked genes between top 10 and all biosamples (**Fig. S7.1d**). We evaluated different combinations of ENCODE-rE2G and PoPs beyond the one highlighted in the main text; we observed that focusing on the one or two nearby genes with the strongest ENCODE-rE2G scores, rather than including all nearby genes, greatly increased the precision of gene predictions (58% for the top gene; 35% for the top 2 genes; versus 4% for all genes linked to a variant), at the cost of some reduction in recall (38% for the top gene; 46% for the top 2 genes; versus 66% for all genes linked to a variant) (**Fig. S7.1e**).

We present 3 example GWAS causal variants for mean corpuscular hemoglobin (MCH) that are linked target genes by ENCODE-rE2G, in potentially causal cell types (**Fig. S7.2**). The GWAS causal variant *rs4927708* for MCH (PIP = 1) is linked by ENCODE-rE2G to the gene *TFRC,* specifically in hematopoietic multipotent progenitor cells. The gene *TFRC* is known to play an important role in iron import and metabolism, and in erythroblast development[14,15]. *TFRC* is not only the top PoPS prioritized gene in the locus, but also is a top PoPs prioritized gene for MCH genomewide (rank 27). The variant *rs218265* (PIP=1) is linked to the KIT gene in K562 and hematopoietic progenitors. KIT is a proto-oncogene that has been linked to anemia and erythroid differentiation[16,17]. For the intronic GWAS causal variant, *rs7599488* (PIP = 0.89), we see an ENCODE-rE2G link to the overlapping gene, *BCL11A*. This variant is located in a well-known clinically severe intronic enhancer for *BCL11A* that possesses erythroid-restricted, adult-stage-specific enhancer activity[18,19].

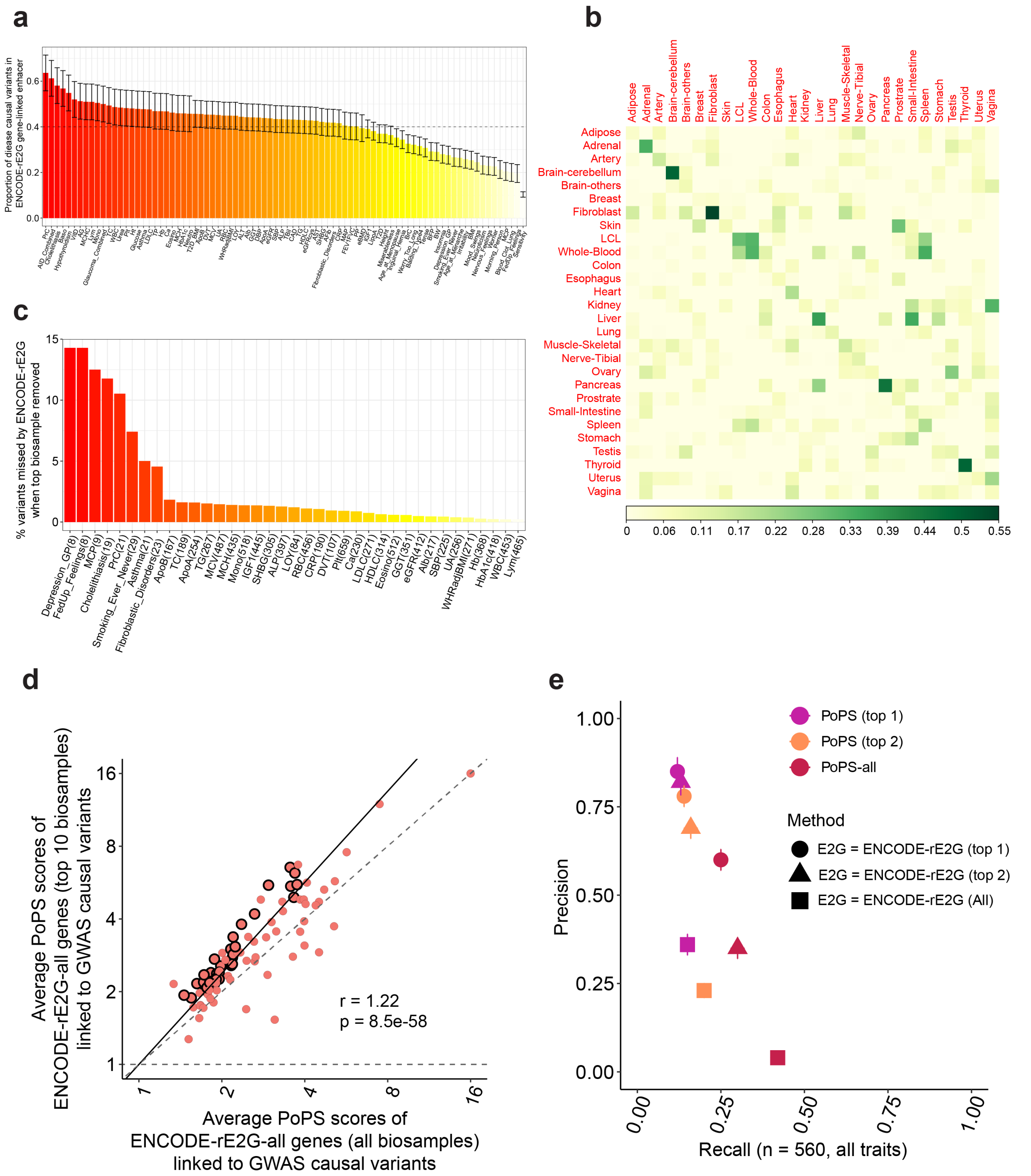


**Fig. S7.1 | Linking common variants fine-mapped from GWAS data by ENCODE-rE2G.**

**a,** Fraction of GWAS fine-mapped variants (PIP > 0.10) for each of GWAS trait that overlap ENCODE-rE2G predictions in at least one of 352 biosamples. The traits are ordered based on the fraction of ENCODE-rE2G overlapping GWAS variants. The error bars denote 95% confidence intervals based on a Jack-knife standard errors over chromosome blocks.

**b**, Heatmap representing the enrichment of GTEx fine-mapped eQTLs (maximum PIP across genes > 0.10) for 28 GTEx tissues in ENCODE-rE2G predictions in sets of biosamples that can be matched to each of these GTEx tissues. For a sparse representation, the enrichments are mean-adjusted across rows (GTEx tissues) and then

across columns (ENCODE-rE2G biosample groups). All adjusted enrichments are thresholded below at 0, and are colored from white to green based on increasing magnitude of adjusted enrichment.
**c**, Barplot representing percentage of GTEx fine-mapped variants missed by ENCODE-rE2G predicted elements when the top globally enriched biosample for the trait is removed from the analysis. Traits are ordered based on the magnitude of this percentage. Results are reported only for traits with at least one variant missed when the top biosample is removed.
**d**, Comparison of the average PoPS scores of genes linked to GWAS fine-mapped variants for a trait by (**Y axis**) ENCODE-rE2G predictions in top 10 globally enriched biosamples based on Panel (b) and (**X axis**) ENCODE-rE2G predictions in all 352 biosamples. Circled dots denote traits with significant (FDR<10%) difference between these averages, the solid line denotes y=x, and the dashed line denotes the regression slope. We report the slope of the regression line.
**e,** For 560 non-coding credible sets corresponding to 94 blood-related traits that are linked to exactly one "putatively causal" gene with a coding fine-mapped variant within 2Mb on either side of the lead variant, we compute the precision and recall in linking it to genes by the top 1, top 2 and all ENCODE-rE2G scored in the window computed across top 10 biosamples globally enriched in GWAS fine-mapped variants for each trait. Additionally, we assess precision and recall when these ENCODE-rE2G predictions in the window are intersected with top 1 or top 2 PoPS scored genes in the window for the given trait. Error bars represent 95% confidence intervals.
Numerical results are reported in Table S13.

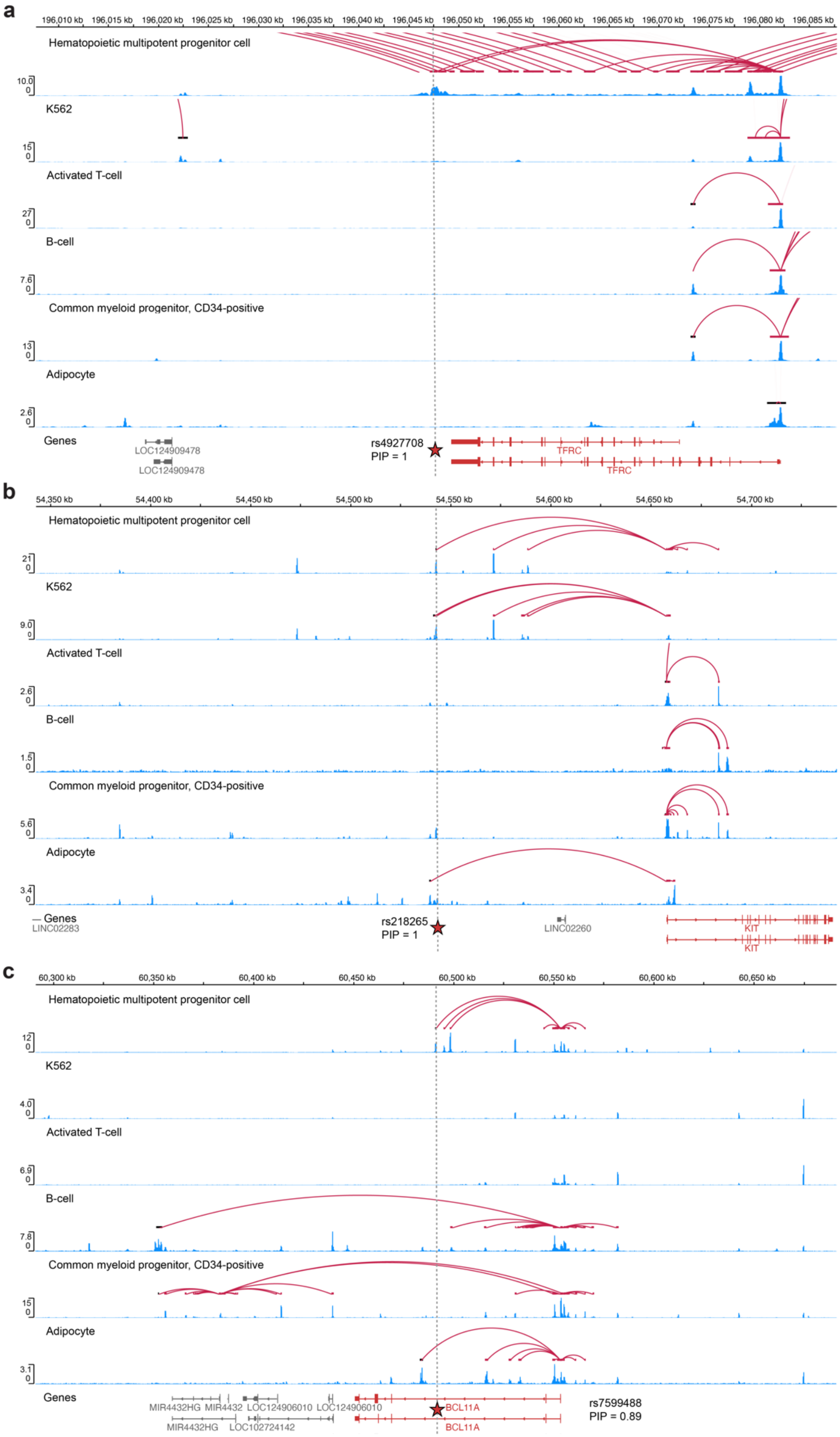


**Fig. S7.2 | Example illustrative GWAS loci linked to target genes by the ENCODE-rE2G + PoPS.** ENCODE-rE2G + PoPS links two confidently annotated GWAS causal variants (PIP > 0.7) for mean corpuscular hemoglobin to target genes - **(a)** *rs4927708* to target gene *TFRC* in hematopoietic multipotent progenitor cells, **(b)** rs218265 to target gene KIT in hematopoietic progenitors and K562 and **(c)** intronic variant *rs7599488* to BCL11A gene specifically linked in hematopoietic progenitors.

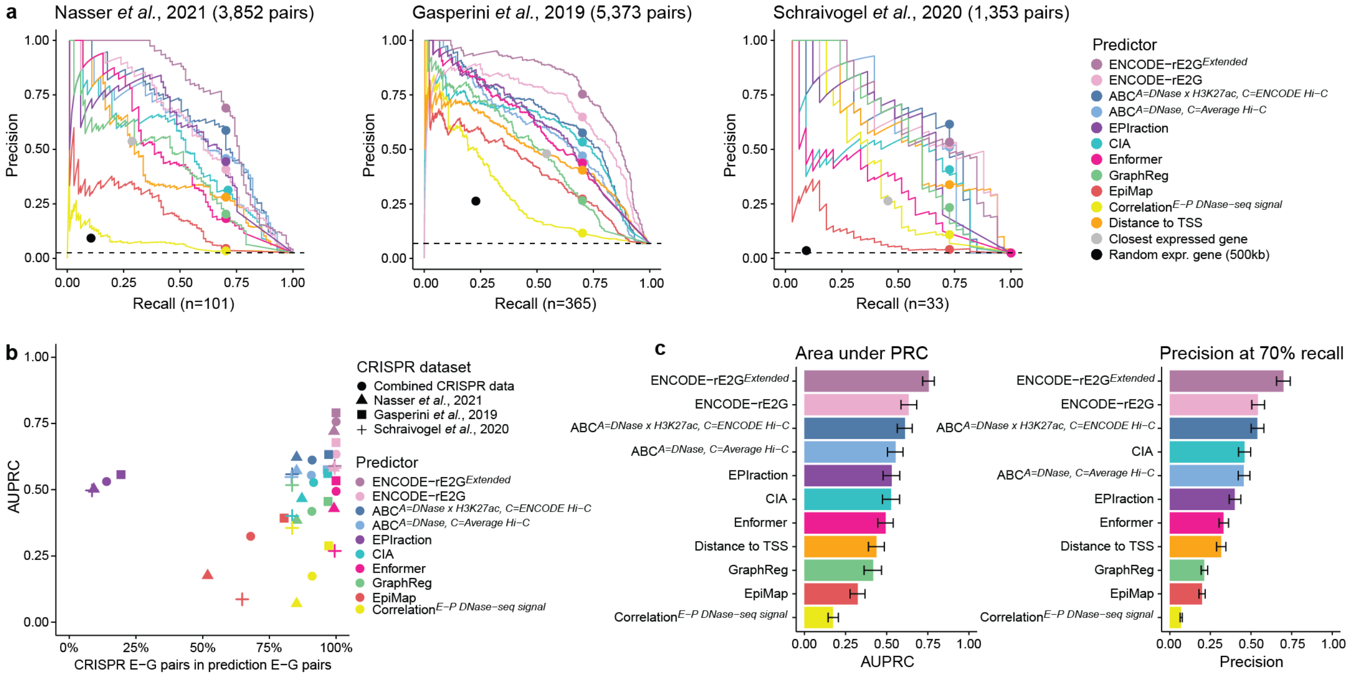


**Fig. S8 | Additional CRISPR benchmarks**

**a,** Precision-Recall curves showing the performance of predictive models benchmarked against each collected CRISPRi dataset separately.

**b,** Performance (AUPRC) of predictive models versus the fraction of element-gene pairs in the combined CRISPR dataset also found in the universe of element-gene pairs in predictions. A smaller fraction indicates that fewer CRISPR E-G pairs are also present in the predictions of a given model. The minimum score for each predictor was used as fill-in value for these missing cases, generally lowering performance of predictive models with low overlap with CRISPR benchmarking dataset.

**c,** Area under the PRC (AUPRC) and precision and inferred cutoff at 70% recall for quantitative predictors benchmarked against the combined CRISPRi dataset. Error bars represent 95% confidence intervals inferred via bootstrap (1000 iterations)

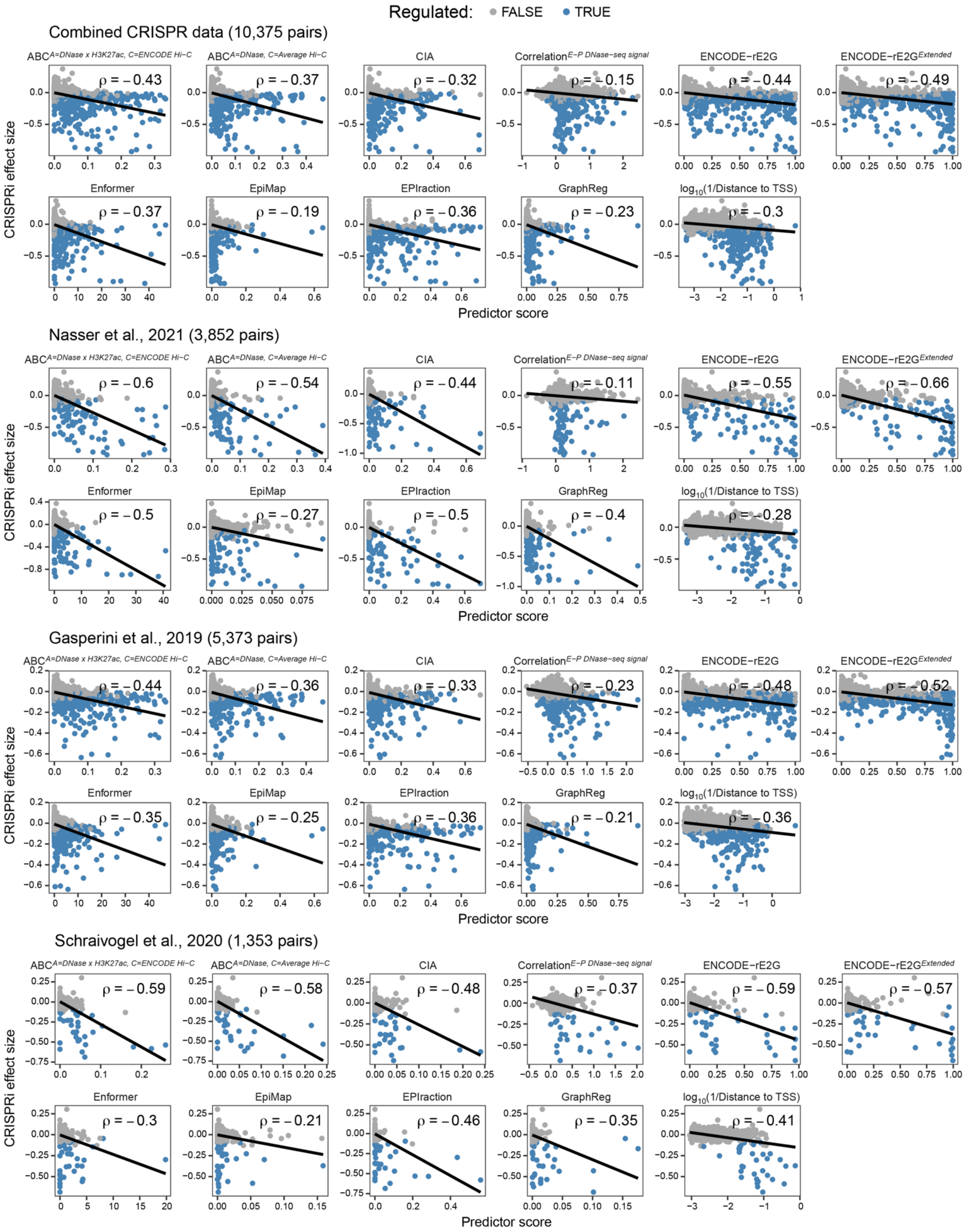


**Fig. S9 | Correlation between predictor scores and CRISPR effect sizes**

Predictor scores versus observed effect size upon CRISPR perturbation for all E-G pairs in the different CRISPR datasets. Effect sizes are percent changes in target gene expression and Pearson correlation coefficient (ρ) between predictor scores and CRISPR effect sizes is provided for each comparison.

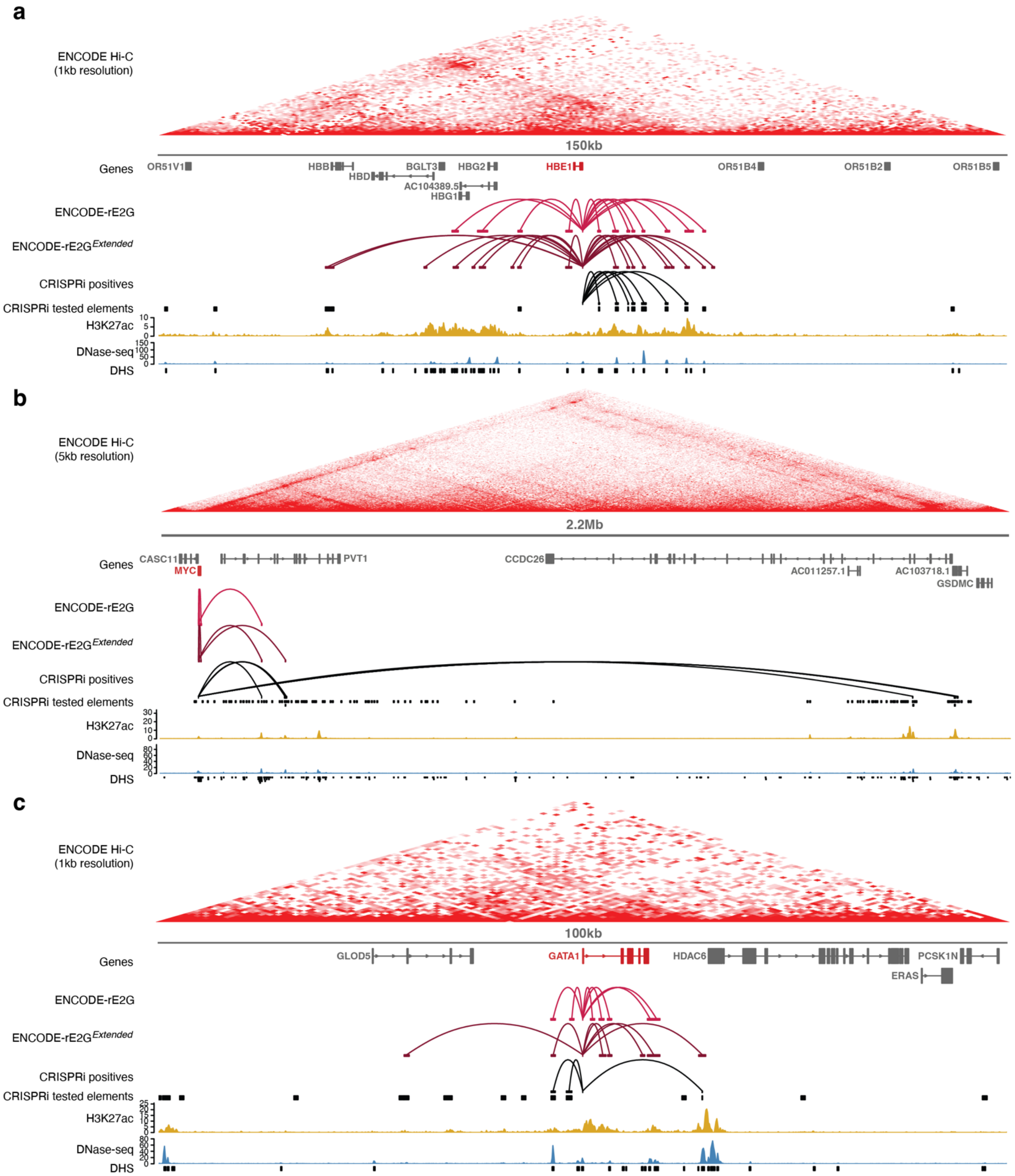


**Fig. S10 | K562 gene locus plots**

Locus plots showing 3D contact (K562 ENCODE Hi-C), genes, predicted and CRISPR positive enhancer-gene regulatory connections, CRISPRi tested elements, H3K27ac and DNase-seq data in K562 for *HBE1* (**a**), *MYC* (**b**) and *GATA1* (**c**). Genes for which enhancer-gene regulatory connections are shown are highlighted in red.

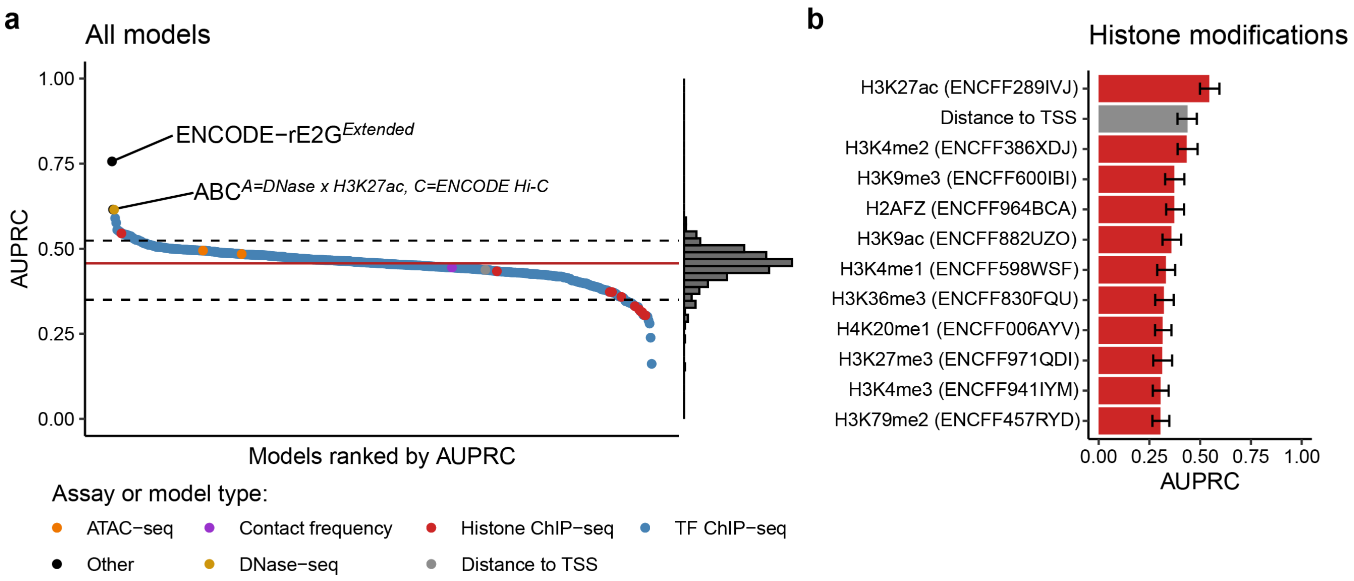


**Fig. S11 | Features of enhancer activity**

**a.** Area under Precision-Recall Curve (AUPRC) performance of Activity-By-Contact (ABC) models using different ENCODE chromatin assays to estimate enhancer activity at predicting experimental results of CRISPRi data in K562 cells. Red line represents the median AUPRC across all models and dashed lines show 90% range of AUPRC. Performance of ENCODE-rE2G$^{Extended}$ and ABC$^{A=DNase\ x\ H3K27ac,\ C=ENCODE\ Hi\text{-}C}$ is shown as reference.

**b.** AUPRC of models using ChIP-seq from different histone modifications to estimate enhancer activity. Error bars represent 95% range of AUPRC values inferred via bootstrap (1000 iterations).

#### References supplementary material
