## Supplementary material for "An encyclopedia of enhancer-gene regulatory interactions in the human genome": Methods


**Methods** **1**

1. Genome build 2
2. Gene reference file for ENCODE-rE2G 2
3. Defining a set of biosamples for ENCODE-rE2G and ABC enhancer-gene predictions. 2
4. Downloading and pre-processing DNase-seq and H3K27ac CHIP-seq data for ENCODE-rE2G and ABC predictions 2
5. Defining candidate elements and element-gene pairs for predictive models and annotations 4
6. Annotating enhancer-promoter pairs with epigenomic measurements and genomic features 4
7. Correlation of enhancer-promoter activity across cell types 8
8. Applying the Activity-by-Contact model to ENCODE4 data 9
9. Applying EpiMap to ENCODE4 data 11
10. EPIraction 11
11. GraphReg 11
12. Enformer 12
13. CTCF loop-constrained Interaction Activity (CIA) 13
14. Training ENCODE-rE2G logistic regression models 13
15. Collection of previously published enhancer-gene predictions 14
16. Generating baseline predictors 15
17. Defining classification thresholds for predictive models 15
18. CRISPRi benchmark 15
19. GWAS benchmark 17
20. eQTL benchmark 19
21. Additional analyses: Assaying enhancer activity 21
22. Additional analyses: Linking variants to genes analyses 21
23. Combinatorial enhancer perturbations at *MYC* locus 21
24. Additional analyses: Enhancer synergy analyses 24
25. Additional analyses: Enhancer-promoter correlation 26
26. Data visualization 26

**References methods** **27**

1. **Genome build**
All coordinates in the human genome are reported using build GRCh38 unless otherwise specified.

2. **Gene reference file for ENCODE-rE2G**
We used the following list of genes and gene promoters for analysis with ENCODE-rE2G and the ABC model: ENCFF672JOP. This file contains one promoter (500-bp region centered on the TSS) per gene symbol, selected to use the RefSeq TSS with the largest number of coding isoforms. This set of promoters was generated by lifting over to hg38 the hg19-based promoter file we previously developed for the ABC model (https://github.com/broadinstitute/ABC-Enhancer-Gene-Prediction/blob/master/reference/RefSeqCurated.170308.bed.CollapsedGeneBounds.TSS500bp.bed) [1]. (Empirically, we found that using this TSS set, as opposed to other approaches such as regenerating this file from hg38 RefSeq annotations or using other gene references such as Ensembl, led to improved accuracy of the ABC model in comparison with CRISPR data in K562 cells, data not shown).

3. **Defining a set of biosamples for ENCODE-rE2G and ABC enhancer-gene predictions.**
We calculated ENCODE-rE2G and ABC predictions across up to 352 biosamples, corresponding to a set of unique biosamples that had DNase-seq data generated by ENCODE and that passed a set of filters on DNase-seq data quality and additional filters we defined based on the quality of the enhancer-gene predictions (**Table S2**). Specifically, we (i) downloaded DNase-seq data and generated an initial list of biosamples as described in Methods Section 4 below; (ii) re-called peaks using MACS2 (Section 5 below); (iii) computed ABC scores (Section 8 below); and (iv) removed biosamples that were outliers in the properties of their peak calls (<120,000 candidate enhancer regions identified) or ABC predictions (<30,000 enhancer-gene pairs predicted by $\mathrm{ABC}^{\mathit{Activity=DNase,\ Contact=Average\ Hi\text{-}C}}$).

4. **Downloading and pre-processing DNase-seq and H3K27ac CHIP-seq data for ENCODE-rE2G and ABC predictions**

   *4.1. Downloading Metadata.* Using the ENCODE API, we downloaded all available DNase-seq and H3K27ac ChIP-seq bam files from the ENCODE Portal. These commands can be found in our encode_v1 branch of the ABC-Enhancer-Gene-Prediction github repository: https://github.com/broadinstitute/ABC-Enhancer-Gene-Prediction/blob/encode_v1/Snakefiles/workflow/scripts/download_log.sh.

   We filtered to files aligned to the GRCh38 genome assembly, files annotated as “released” and files processed using the ENCODE4 processing pipeline. Because ENCODE stores multiple versions of files for each experiment accession, we selected the most up-to-date version (as denoted by “Date added” per biological and technical replicate. For each experiment accession, we pooled biological and technical replicates (seen as comma-delimited files in the metadata table provided).

   For predictions involving both DNase-seq and H3K27ac-seq, we merged DNase-seq and H3K27ac ChIP-seq metadata files based on the following columns: 'Biosample term name', 'Biosample organism', 'Biosample treatments', 'Biosample treatments amount', 'Biosample treatments duration', 'Biosample genetic modifications methods', 'Biosample genetic modifications categories', 'Biosample genetic modifications targets', 'Biosample genetic modifications gene targets', 'File assembly', 'Genome annotation', 'File format', 'File type', 'Output type'. This generated 2686 combinations of every DNase-seq experiment with every H3K27ac ChIP-seq for each of 94 unique biosamples available on ENCODE. We provide a metadata list of the different combinations and assign unique IDs to each prediction. Files to download DNase-seq and H3K27ac ChIP-seq metadata files can be located at https://github.com/broadinstitute/ABC-Enhancer-Gene-

Prediction/blob/encode_v1/Snakefiles/workflow/scripts/download_log.sh. Filtering steps used to merge metadata files can be located at https://github.com/broadinstitute/ABC-Enhancer-Gene-Prediction/blob/encode_v1/Snakefiles/workflow/scripts/grabDownload.py.

For predictions involving DNase-seq only, we merged DNase-seq metadata files based on the following columns: 'Biosample term name', 'Biosample organism', 'Biosample treatments', 'Biosample treatments amount', 'Biosample treatments duration', 'Biosample genetic modifications methods', 'Biosample genetic modifications categories', 'Biosample genetic modifications targets', 'Biosample genetic modifications gene targets', 'File assembly', 'Genome annotation', 'File format', 'File type', 'Output type'. This generated 1535 entries of DNase-seq experiments for 352 unique biosamples available on ENCODE. We provide a metadata list of the available combinations and assign unique IDs to each prediction. For the final 352 unique biosamples used to generate ENCODE-rE2G predictions, we prioritized samples that were the most recently deposited to the ENCODE portal and focussed on samples that were part of the “ENCODE 4” processing pipeline. In cases where there were multiple entries of similar biosamples, we ranked biosamples based on the labs that generated these datasets, mainly: UW > Duke > UMass. Finally, if after these filtering steps, there were still multiple entries of similar biosamples, we arbitrarily picked the first among all duplicates to get a unique set of 352 biosamples.

4.2. *Data preprocessing.* We processed the bam files differently depending on whether the bam files were paired-ended or single-ended. Information on the run type (single-ended or paired-ended) of bam files was obtained by mapping ENCODE experiment accessions in the fastq metadata with the bam metadata, because this information was not readily available in the bam metadata provided by ENCODE. For ambiguous files that were annotated as both single-ended and paired-ended, we manually corrected these entries using `samtools view -f 0x1.`

For single-ended bam files, we removed reads that were unmapped, mate unmapped, not primary alignment, or failing platform (-F 780), and further filtered for multi-mapped reads with MAPQ < 30 (-q 30). This filtering was performed using the command `samtools view -F 780 -q30 -u single-ended.bam`. Single-end bam files were not further filtered to remove duplicate fragments, because duplicate fragments are difficult to detect with single-end data and running removal of duplicate reads would lead to undercounting of the true number of duplicate fragments giving the high complexity and sequencing depth of these DNase-seq datasets.

For paired-ended bam files, we directly downloaded the “filtered” alignments from ENCODE. These bam files were previously processed by the encode atac-seq-pipeline and encode chip-seq-pipeline filtering steps. Filtering steps are as follows: We removed reads that were unmapped, mate unmapped, not primary alignment, failing platform and PCR or optical duplicates (-F 1804), as well as multi-mapped reads with MAPQ < 30 (-q 30). We retained properly paired reads (-f 2). This filtering was performed using the command `samtools view -F 1804 -q 30 -f2 -u paired-ended.bam`, (samtools version 1.7). We marked and filtered PCR duplicates using Picard’s MarkDuplicates (Picard version 1.126).

### 5. Defining candidate elements and element-gene pairs for predictive models and annotations

We defined a set of candidate elements in each biosample based on ENCODE DNase I hypersensitivity data using an approach similar to that we previously described for the ABC model[1]. We used this same set of candidate elements to construct the following predictive models: ENCODE-rE2G, ABC, GraphReg-LR, and baseline models.

5.1. *Calculating candidate elements.* For experiment accessions that have multiple biological and technical replicates, we call peaks using MACS2 on all replicates of either DNase-seq or ATAC-seq as a measure of chromatin accessibility. In situations where peak calling failed when using multiple biological and technical replicates due to poor data quality, we used just one replicate. We initially considered all peaks with P<0.1 and then removed any peaks overlapping regions of the genome that have been observed to accumulate anomalous number of reads in epigenetic sequencing experiments (blacklisted regions, downloaded from https://sites.google.com/site/anshulkundaje/projects/blacklists). We then counted DNase-seq (or ATAC-seq) reads overlapping these peaks and kept the 150,000 with the highest number of read counts. We then resized these peaks to 500-bp in length centered on the peak summit. To this peak list, we added 500-bp regions centered on the transcription start site of all genes. Any overlapping regions resulting from these additions or extensions were merged.

We define these extended and merged peaks as candidate elements. We classified each candidate element as a promoter, genic or intergenic element. Promoter elements are those that are within 500 bp of any TSS contained in the promoter reference file described in Methods Section 2. Genic elements are those contained within any annotated gene body. Intergenic elements are all other candidate elements. We denote any genic or intergenic element as a 'distal' element.

5.2. *Computing a list of candidate element-gene pairs.* We generated a list of candidate element-gene pairs to consider in predictive models using the list of elements described in Methods S5.1 and the list of promoters described in Methods S2, including all pairs located within 5 Mb of one another.

### 6. Annotating enhancer-promoter pairs with epigenomic measurements and genomic features

We systematically computed features of enhancer-promoter pairs, including using measurements of chromatin state, measurements/estimates of 3D contact frequencies, and other features including genomic distance and element density (**Table S3**).

6.1. *Chromatin state at enhancers and promoters.*

6.1.1. *Calculation of DNase, H3K27ac, and other ChIP-seq signals.* We first utilized the bam files for the corresponding assays from the ENCODE portal. For information on any preprocessing we did, please refer to S4.2 on data preprocessing. Once the bam files have been preprocessed, we proceed to use `bedtools coverage -counts` to count reads for candidate element regions, gene bodies and gene promoter regions. We calculated cell-type specific enhancer activity for all candidate enhancers by computing the geometric mean of DNase-seq (and, if applicable, H3K27ac ChIP-seq count reads) because we expect that strong enhancers and strong promoters would have strong signals for both. Once DNase-seq (and, if applicable H3K27ac ChIP-seq count reads) have been computed for candidate element regions, gene bodies and gene promoter regions, we quantile normalized ("rank normalized") these values with a reference located here:

https://github.com/broadinstitute/ABC-Enhancer-Gene-Prediction/blob/encode_v2/Snakefiles/workflow/scripts/EnhancersQNormRef.K562.txt. This quantile normalization reference was generated using K562 predictions made in hg19 for the original ABC Paper[1]. We first distinguish quantile normalization references for promoters and non-promoter regions respectively. Generally, we ranked the promoters in K562 by their magnitude, calculated the average value for all promoters with the same rank and substituted the values of all promoters occupying the same rank. This was repeated for non-promoter regions. This quantile normalization step is key in ensuring that we can compare read counts, and any downstream features (such as the ABC Score), across different cell types. Additionally, quantile normalization is utilized to minimize technical variability that might have been introduced in the experimental setup. We've found that using this reference works well across hg19 and hg38 genome builds. Code for these calculations can be found by running the `run.neighborhoods.py` script in the ABC codebase.
For our run specifically, we used:

```
python run.neighborhoods.py --
candidate_enhancer_regionPeaks/macs2_peaks.narrowPeak.sorted.can
didateRegions.bed
--genes
reference/hg38/RefSeqCurated.170308.bed.CollapsedGeneBounds.hg38
.bed --DHS
example_chr22/input_data/Chromatin/wgEncodeUwDnaseK562AlnRep1.ch
r22.bam
--chrom_sizes example_chr22/reference/chr22
--ubiquitously_expressed_genes
reference/UbiquitouslyExpressedGenesHG19.txt
--cellType K562 --qnorm reference/EnhancersQNormRef.K562.txt
--outdir Neighborhoods/
```

6.2. *3D contact frequencies from Hi-C.*

6.2.1. *Calculating contact frequency from cell-type-specific Hi-C data*. For 3D contact values from Hi-C used for predictions with ENCODE-rE2G$^{Extended}$ and the ABC model, we used our Hi-C preprocessing approach previously described for the ABC model[1], with slight modifications. Specifically, we used normalized ENCODE Hi-C contact maps (at 500-bp, 1-kb, or 5-kb resolution), and processed these maps in two steps:

For rows and columns from ENCODE Hi-C experiments corresponding to SCALE normalization factors < 0.25, we did not use SCALE normalization (these typically correspond to 5-kb bins with very few reads). Instead, we linearly interpolated the Hi-C signal in these bins by calculating an expected value based on power-law fit (S6.2.3 for more information on how this power law relationship is calculated).

Each diagonal entry of the Hi-C matrix was replaced by the maximum of its four neighboring entries. As previously described[1], this step is taken because the diagonal of the Hi-C contact map corresponds to the measured contact frequency between a 5-kb region of the genome and itself. The signal in bins on the diagonal can include restriction fragments that self-ligate to form a circle, or adjacent fragments that re-ligate, which are not representative of contact frequency. Empirically, we observed that the Hi-C signal in the diagonal bin was not well correlated with either of its neighboring bins and was influenced by the number of restriction sites contained in the bin.

We then computed Contact for an element-gene pair by rescaling the data as follows:

- We set the Contact of the element-gene pair to the Hi-C signal at the bin of this row corresponding to the midpoint of E. For element-gene pairs that do not have a corresponding contact value (i.e., NaN), we set contact to zero.
- For distances greater than 5 kb, we added a small adjustment (pseudocount) based on the power law expected count at a given distance threshold (as predicted by the power-law relationship between contact frequency and genomic distance). The distance threshold is usually determined by the resolution of the Hi-C data used. Distances less than 5 kb were given the pseudocount computed at 5 kb. We compute the power law by utilizing the steps in Section 6.3.
- We found that different Hi-C datasets have slightly different power-law parameters. To weight all cell types equally in generating an average Hi-C profile, we scale the Hi-C profile in a given cell type by the cell-type specific gamma parameter from the power law relationship in that cell type. The scaling factor at distance d is given by d ^ ($gamma_{ref}$ – $gamma_{celltype}$), where $gamma_{ref}$ is the reference gamma parameter. For this study, we used gamma = 0.86 as a reference.

These processing steps are implemented in: https://github.com/broadinstitute/ABC-Enhancer-Gene-Prediction/blob/5c024fe29d19c06ee47eacdf0634e0beaf7ac235/Snakefiles/workflow/scripts/predictor.py#L55.

We performed similar steps to calculate normalized ENCODE Hi-C contact maps at 1-kb and 500-bp resolution.

For some analyses, we also explored using unnormalized ENCODE Hi-C data as an estimate of 3D contact frequency. In these cases, these steps were repeated to calculate unnormalized ENCODE Hi-C contact maps at 500-bp, 1-kb and 5-kb resolution. The only difference is instead of using the normalized Hi-C values, we directly used the quantitative signal observed in the bin containing the center of enhancer and TSS of the gene. This is illustrated here: https://github.com/broadinstitute/ABC-Enhancer-Gene-Prediction/blob/encode_v1/Snakefiles/workflow/scripts/hic.py#L104 where no normalization file was used to normalize the contact maps (hic_norm_file == None).

*6.2.2.* *Generating average Hi-C contact maps.* We previously found that, for most genes, using an average Hi-C profile in the ABC model yields similar performance to using a cell-type specific Hi-C profile[1], which we confirmed here (**Fig. S2.1f**). To facilitate making predictions in a large panel of cell types, including those without cell type-specific Hi-C data, we have provided an average Hi-C matrix (averaged across 35 cell lines and tissues, at 5kb resolution). We used a total of 35 diploid primary cell lines and tissues for averaging listed (**Table S4**). We extracted normalized Hi-C values across all 35 cell types and tissues, and proceeded to merge them based on similar contact bins (i.e. binX, binY). For each contact bin, we calculated an average of non-zero values to produce the final average Hi-C matrix. Code to generate an average Hi-C matrix is located at https://github.com/broadinstitute/ABC-Enhancer-Gene-Prediction/blob/master/src/makeAverageHiC.py. The average Hi-C contact map can be downloaded from: ENCFF134PUN.

6.3. *Estimating the power-law relationship between 3D contacts and distance.* It has been previously shown that Hi-C contact frequencies generally follow a power-law relationship (with respect to genomic distance). We computed a power-law fit for Hi-C datasets by first examining the relationship between genomic distance (binned at 5-kb resolution) and normalized 3D contact values. For each bin, we calculated a value for each distance bin by normalizing the hic contact value (normalized over total number of values in that distance bin). Finally, we log-transformed both the distance bins and mean hic contact values and fitted the data using linear regression. Once a line of best fit is obtained using the Hi-C data $(Contact \sim (hic\ distance\ bin) \cdot \gamma + c)$, we extracted the slope and intercept values denoted as gamma and c (**Table S5**). We utilized the power-law relationship between 3D contact and distance described by these learned parameters (slope and intercept values) to impute missing 3D contact values as described in Methods S6.2.1 and to conduct certain analyses using the power-law in place of 3D contact data (*e.g.*, see **Note S2**).

6.4. *Detecting CTCF Contact Domains (CCD)*
We first computed CTCF loop coverage for each base by summing up the PET count of all CTCF loops from in situ ChIA-PET that span the base. This base-pair resolution track was binned at 200-bp resolution by using the maximum signal in the bin. Then, we applied a Hidden Markov Model (HMM) with two hidden states on this track to segregate the genome into high-coverage and low-coverage domains based on their levels of CTCF interaction frequency.  We call the domains with high CTCF loop coverage as CTCF contact domains (CCD). The HMM model is imported from the python sklearn.hmm package.

6.5. *Computing CTCF ChIA-PET features: "Loop Cross" and "Loop Contain"*
CTCF loops for K562 and GM12878 from ChIA-PET were downloaded from the ENCODE portal (ENCSR184YZV, ENCSR597AKG). 'Loop Contain' is given by the ratio of total PET count of all CTCF loops containing both the enhancer and the promoter over total PET count of CTCF loops containing the promoter. 'Loop Cross' is given by the ratio of total PET count of all CTCF loops across the enhancer and the promoter pair over total PET count of CTCF loops containing the promoter. All CTCF loops have a minimum PET count of 3 and maximum length of 500kb.

6.6. *ENCODE Hi-C loops and domains.* For measurements involving binary loop calls and domain calls from ENCODE Hi-C data, we obtained data from ENCODE Accessions ENCFF134HIZ and ENCFF173VDJ respectively. Binary loop and domain calls were converted to contain a start region, end region and contact value associated with each entry. This was then intersected with the CRISPR perturbation data to obtain binary predictions.

6.7. *Features related to promoter class.*  We used 3 correlated metrics to define classes of promoters. (i) An enhancer "complexity" score, in which we previously calculated the degree to which ABC enhancers for a given gene are correlated across cell types[2]. Higher correlations indicate that the enhancer landscape for a gene is similar across cell types, whereas lower correlations indicate that the enhancer landscape differs. (ii) Promoter class as defined by Bergman and Jones et al.[3], in which a subset of "P2" promoters were measured to be less sensitive to distal enhancers, and then additional P2 promoters were identified across a genome based on logistic regression of sequence and chromatin features. (iii) Is this a ubiquitously expressed housekeeping gene? We defined ubiquitously expressed data based on FANTOM5 gene expression data as previously described[3].

6.8. *Features of genomic position*

6.8.1. *Distance to TSS.* Distance to TSS for each candidate element-gene pair was computed as the absolute distance between the TSS and the center of the candidate element.

6.8.2. *How many protein-coding TSSs away is the candidate element from the promoter? (i.e., how many protein-coding gene TSSs are located between the enhancer and promoter? 0 = closest TSS).* Utilizing the full list of enhancer-gene predictions, we generated a bed file by starting at the midpoint of each enhancer and extending it to end at the promoter of the target gene. Each bed file now consists of the chromosome, the midpoint of the enhancer and the promoter of the target gene. Using *bedtools intersect,* we intersected this bed file with the list of gene transcription start sites and counted the total number of promoters that intersected each enhancer-gene connection, excluding the target gene promoter.

6.8.3. *How many other nearby enhancers/DNase peaks are within 5 kb? (count or quantitative sum of Activities, cell-type specific).* For each candidate element, we annotated the element with either the count or summed enhancer Activity of all other nearby candidate elements within 5 kb. (An element is counted if both its start and end coordinate are within 5 kb of the midpoint of the query element). For enhancer Activity, we used the definition of the corresponding ABC model (see Methods Section 8 below), *i.e.* using quantitative H3K27ac and DNase signals for certain models including ENCODE-rE2G$^{extended}$ and only DNase signals for the DNase-only models including ENCODE-rE2G.

6.9. *Sequence conservation.* For each candidate element, we created 100bp windows across the element and calculated the mean conservation across each window using PhastCons scores for multiple alignments of 29 genome sequences to the human genome and PhyloP scores for multiple alignments of 99 vertebrate genomes to the human genome. We then calculated the mean, min, max of these per window scores as the score per candidate element.

### 7. Correlation of enhancer-promoter activity across cell types

cCRE-TSS pairs were generated using ENCODE4 cCREs obtained from http://users.wenglab.org/moorej3/Registry-cCREs-WG/V4-Files/GRCh38-cCREs.V4.bed.gz and the curated RefSeq TSS list. DNase-seq reads were counted within 100bp from the center of each cCRE and within a 500bp window centered on each TSS across 89 ENCODE biosamples (**Table S6**). DNase-seq reads were normalized for library size to 1 million reads per biosample across all cCREs, respectively all TSS, and log transformed with adding one pseudocount. Pearson and Spearman correlation coefficients were calculated for each cCRE - TSS pair within 1Mb using R (4.1.1). Generalized Least Square (GLS) regression models were used to model DNase-seq reads at TSSs as a function of DNase-seq reads at cCREs within 1Mb. To account for global correlation among biosamples, a correlation matrix between all samples was computed from normalized DNase-seq reads in all cCREs. The gls function from the nlme (3.1-162) R package was used to fit regression models using the correlation matrix as correlation structure. The cCRE coefficients were extracted from the fitted models and used as measurement of correlation between cCRE and TSS activity.

For DNase-RNA correlations, RNA-seq data for 96 ENCODE PolyA+ RNA-seq samples (**Table S7**) was downloaded and normalized to 1 million reads per sample. The same correlation approach as described above was applied, but instead of promoter accessibility, DNase-seq accessibility at cCREs was correlated with library size normalized and log transformed RNA-seq TPM values from genes with promoters within 1Mb of cCREs.

### 8. Applying the Activity-by-Contact model to ENCODE4 data

*8.1. Updates to the Activity-by-Contact Model.* We applied the Activity-by-Contact model using the same formula as previously described[1], with updates and improvements to the processing code, data inputs, and quality control metrics:

*8.1.1. Enabling biological and technical replicates when defining candidate elements.* We enabled the calling of peaks using MACS2 on all replicates of either DNase-seq or ATAC-seq as a measure of chromatin accessibility (previous versions of ABC used just one replicate to call peaks and define candidate elements). We then counted DNase-seq (or ATAC-seq) reads overlapping these peaks from all replicates, and report the average reads overlapping these peaks. We kept the 150,000 peaks with the highest number of read counts and annotated these as candidate regions. Here, we used 1 or more DNase-seq replicates as described above.

*8.1.2. Enabling biological and technical replicates when calculating enhancer activity from DHS and H3K27ac ChIP–seq signals.* We enabled the computation of the geometric mean of DNase-seq (and, where applicable, H3K27ac ChIP-seq) signals from all biological and technical replicates. For every enhancer element, we calculated enhancer "Activity" as the averaged read coverage from all replicates for DNase-seq (and, where applicable, H3K27ac ChIP-seq) signals. Here, we used 1 or more replicates as described above.

*8.1.3. Reproducibility.* We improved overall reproducibility of running ABC by packaging the code using Snakemake, allowing users to download metadata files, preprocess files and generate ABC predictions using one line of code.

*8.1.4. Reproducibility of enhancer-gene connections across experiment accessions and biosamples.* We generated global statistics of enhancer-gene connections predicted by ABC across different experiment accessions for each biosample to measure the variance in using different experiments as input into ABC. For further downstream analysis, we utilized these global statistics to filter high-quality predictions based on the number of significant enhancer-gene connections, enhancer-gene distance and other metrics. Poor predictions could be a result of low coverage signal files from DNase-seq or H3K27ac ChIP-seq input files, and were thus not included in further analysis. Here, we used the following statistics to remove low-quality biosamples: <120,000 peaks called from DNase bam files, <30,000 candidate enhancer regions, <30,000 significant enhancer-gene links.

*8.1.5. Additional updates.* We updated the codebase to use ENCODE Hi-C data, including to adjust the Hi-C preprocessing steps as described in Methods Section S6.2.1. We updated the ABC pipeline to support analyses in hg38, including by adding an hg38 reference promoter file (see Methods Section S2).

*8.2. Applying ABC pipeline to ENCODE4 data*. We applied this updated ABC pipeline to compute various versions of ABC scores, using different combinations of input datasets. Importantly, different ABC score thresholds are used for each of these types of ABC models, selected based on obtaining 70% recall in comparison to CRISPRi perturbation data (see **Table S1**):

$\text{ABC}^{\textit{Activity=DNase, Contact=Average Hi-C}}$. We computed one set of ABC predictions for 352 biosamples with DNase only, in which we used DNase to estimate activity, and average ENCODE Hi-C data to estimate 3D contact frequency (**Table S1**). The threshold used for $\text{ABC}^{\textit{Activity=DNase, Contact=Average Hi-C}}$ scores was 0.014157.

$ABC^{Activity=DNase+H3K27ac,\ Contact=Average\ Hi\text{-}C}$. We computed one set of ABC predictions for 330 biosamples with both DNase and H3K27ac, in which we used DNase and H3K27ac to estimate activity, and average ENCODE Hi-C data to estimate 3D contact frequency (**Table S1**). The threshold used for $ABC^{Activity=DNase+H3K27ac,\ Contact=Average\ Hi\text{-}C}$ scores was 0.014157.

$ABC^{Activity=DNase+H3K27ac,\ Contact=ENCODE\ Hi\text{-}C}$. We computed one set of ABC predictions for K562 and GM12878, the two Tier 1 samples, in which we used DNase and H3K27ac to estimate activity, and cell-type specific ENCODE Hi-C data to estimate 3D contact frequency (**Table S1**. The threshold used for $ABC^{Activity=DNase+H3K27ac,\ Contact=ENCODE\ Hi\text{-}C}$ scores was 0.025693.

In K562 cells, we computed various other versions of the model using different assays for Activity (see **Fig. S11**) and Contact (see **Note S2**).

For all ABC models, ENCODE Hi-C data was analyzed at 5-kb resolution unless otherwise specified.

8.3. *Evaluating the sensitivity of ABC model predictions to transcription start site selection.* In Fulco *et al.*, 2019[1], we determined the sensitivity of ABC to promoter and gene annotations. In an attempt to study the effects of using different promoter annotations, we provide ABC predictions generated using 3 types of gene annotation files which we will call: RefSeq Annotated, Gencode V29 Unique, Gencode V29 AltTSS.

We lifted over the hg19 RefSeq gene and TSS annotation file used in the original ABC code to hg38. This set of gene and TSS annotation files considers only one transcription start site per gene. The original RefSeq gene annotations list was curated by selecting the gene isoform that was the most used.

In an attempt to enable consistency with the ENCODE portal, we curated a gene and tss annotation file using GENCODE v29 gene annotation file. For each gene, we selected the longest transcript, level 1 annotations (if not available, we selected level 2 annotations) as determined by GENCODE to pick one unique TSS. We then called transcription start sites by extending the beginning of each transcript by 250-bp on each side to produce 500-bp promoter regions.

Given the knowledge of alternative transcription start sites where a single gene might undergo alternative transcription initiation (ATI), we developed a simple heuristic to select the top two promoter regions for each gene using the GENCODE v29 transcript annotation file from the GENCODE browser (as denoted by GENCODE v29 gene annotation). We started by first using the refGene file from the USCS Genome Browser and filtered the refGene file for only protein-coding regions. We obtained cell-type specific DNase and H3K27ac bam files to obtain counts for each TSS region in the filterted refGene file. These counts were used to calculate promoter activity, which is the geometric mean of the DNase and H3K27ac counts for each region respectively. For every gene, we proceeded to select the top two TSS regions as determined by promoter activity. In addition, the top two TSS regions had to be at least 500 bp apart from each other. We will reference this alternative transcription initiation (ATI) as alternative TSS annotations.

Code and reference files are available at:
https://github.com/EngreitzLab/ABCGeneReference

Comparing RefSeq, alternative TSS annotations and GENCODE v29 gene annotations, we found that there were approximately 15818 TSS annotations that were similar when using these annotations in K562 cells.

**9. Applying EpiMap to ENCODE4 data**

We modified the EpiMap[4] correlation-based peak gene linking methodology to (1) update the genome build to GRCh38, (2) focus on distal enhancers, (3) de-bias compositional changes, and (4) better model peak-gene distance.

To do so, we utilized GRCh38 mappings of the ENCODE4 cCRE index and mapped histone modifications and DNase-seq profiles in EpiMap to these locations. To do this for all 833 EpiMap biosamples, we obtained the corresponding GRCh37 locations of these GRCh38 cCREs via UCSC liftOver. We then used UCSC bigWigAverageOverBed to get the signal associated with these regions over imputed and observed marks in EpiMap. To get active enhancers per biosample, we filter these cCREs to those overlapping at least 50% of one of the following ChromHMM states: EnhA1, EnhA2, EnhWk, EnhG1, EnhG2, EnhBiv (this step is different than the original EpiMap predictions, in which promoter and TSS-proximal ChromHMM steps were also used).

Then, we updated the core EpiMap peak-gene linking methodology. We utilized ZCA whitening to de-bias compositional changes in the EpiMap sample-by-cCRE matrices per-mark as well as in the RNA-seq FPKM matrix. When correlating cCREs to genes, we utilize weighted correlations that use weights from sample-sample correlations generated from the sample-by-cCRE matrix. These weighted correlations weigh similar samples more to provide more accurate cell type specific cCRE-gene links.

Finally, we utilize an ABC-style normalization to better model peak-gene distances. When combining cCRE-gene links across ChromHMM states, we take the probability of links output from the model, multiply these by the ABC distance decay (Power=0.87), and then normalize all links per-gene such that the sum of all link scores per gene equals 1.

**10. EPIraction**

We included in our model the information about H3K27ac epigenetic signal correlation at promoters and enhancers provided by the EPIraction[5]. Briefly, the authors collected H3K27ac data for 1,507 samples from 85 different tissues (**Table S8**). These samples were uniformly processed and jointly normalized. For every enhancer-promoter pair they calculated a weighted Pearson correlation coefficient of H3K27ac signal across all 1,507 samples up-weighting the samples, representing the tissue of interest.

**11. GraphReg**

GraphReg[6] is a chromatin-interaction-aware gene regulation model which uses 1-dimensional epigenomic signals as well as 3-dimensional genomic interaction graphs, extracted from Hi-C data, to predict the gene expression values by graph attention networks. As GraphReg is a distal-enhancers-aware predictive model, its feature attribution values for each gene could be informative for the task of E-G predictions. The methods described in this section are the same as GraphReg paper but the input epigenomic and Hi-C datasets are different. As a result, the trained models are different from the published models. Here we trained GraphReg models in two cell types K562 and GM12878 using three informative epigenomic features at 100bp resolution, DNase, H3K27ac, and H3K4me3, and extracted genomic interaction graphs from ENCODE Hi-C contact maps at the resolution of 5kb (**Table S8**). We used HiC-DC+[7] with the false discovery rate (FDR) of 0.1 to extract the genomic interaction graphs containing the interactions up to 2Mb. For gene expression, we have used CAGE-seq, binned at 5kb to be consistent with the Hi-C graphs, and predicted it at all the bins. We have used chromosome-wise cross-validation for the predictions and held out two chromosomes for each test and validation sets and trained on the remaining chromosomes. As such, we have trained 11 GraphReg models. After training, the models are used for feature attributions. As we need to get the feature attributions of all the genes, we have used only one model and the gradient method for feature attribution to speed up the process. The process of calculating the gradients of the genes with respect to the input features works as follows. We first retain all the genomic bins that contain the TSSs' of the genes. If the minimum distance between TSS and the boundaries of the 5kb bin is more than 200bp, we assign only that bin to that gene, and if the distance between TSS and the right (left) boundary is less than 200bp, we assign that bin plus the right (left) 5kb bin to that gene. Then for each gene, we calculate the gradient of the model output for its assigned bins (1 bin or the sum of two bins) with respect to the 1D epigenomic features in the input of the model using automatic differentiation in TensorFlow. This means that we will have three gradient values (one for each epigenomic feature used) at each 100bp bins for each gene. These gradient values are always non-zero for the promoter bins but could be zero for the distal enhancer regions if there are no E-G interactions.

For an initial set of GraphReg scores (denoted GraphReg$^{Raw\ Scores}$, see **Table S1**), we use our previous approach[6] in which we use the gradient values and input epigenomic features to define a score which could be used for enhancer predictions as follows:

$$s[i,g] = |\,exp(x_{dnase}[i]-1) \times dg/dx_{dnase}[i] + exp(x_{h3k27ac}[i]-1) \times dg/dx_{h3k27ac}[i]\,|,$$

$$score_{e,g} = \Sigma_{k=-L:L} s[i_{mid}+k,g] / \Sigma_k (s[k,g]),$$

where $x_{dnase}$ and $x_{h3k27ac}$ denote log-normalized ($log(x+1)$) DNase, H3K27ac values used in the training of the GraphReg model (they are exponentiated back to their original values here); $dg/dx[i]$ denotes the gradient of gene g w.r.t. the epigenomic feature $x$ at bin $i$; $s[i,g]$ denotes the absolute value of the dot product of the two epigenomic features with their corresponding gradients at the bin $i$ for the gene $g$; $i_{mid}$ denotes the index of the bin related to the summit of the enhancer element $e$; $L$ is the one-sided window size around the enhancer summit; and $score_{e,g}$ is the score of enhancer $e$ for the gene $g$. This unsupervised score worked well in the GraphReg paper where only the genes with at least 10 candidate enhancers and at least one true functional enhancer were considered, as the main goal was per-gene enhancer rankings. However, if the goal is to have a global score for every E-G pair in the genome, this unsupervised score performs suboptimally for the task of enhancer prediction (**Fig. 2b,c**).

To remedy this issue and in line with the logistic regression approach used in this paper, we also suggested supervised GraphReg scores which uses all the features extracted from the trained GraphReg models and trains a logistic regression model using those features to predict the CRISPR data in K562 cell, as for ENCODE-rE2G models (denoted GraphReg$^{LR}$, see **Table S1**). This supervised model uses 19 features: three epigenomic features in the enhancers (Dnase, H3K27ac, and H3K4me3) and three in the promoters, six gradient features in the enhancers and six in the promoters, and we also include distance between enhancers and genes as another feature because we have observed a strong effect of the distance on the performance. The reason that we have six gradients instead of three in each enhancer and promoter is that we use both maximum and minimum values of the gradients around the element summit spanning $2L+1$ bins. We have used $L=8$, which means that the spanning length around each enhancer or promoter element is 1700bp.

### 12. Enformer

Enhancer-gene predictions using Enformer were created for sets of element-gene pairs. For benchmarking the performance against CRISPR data, Enformer predictors were generated for all element-gene pairs in the combined CRISPR dataset. For benchmarking predictions against *eQTL variants in GM12878* two lists of DNase peak-gene pairs were used: 1) a list of all MACS2 DNase peaks in GM12878 overlapping the filtered set of LCL eQTLs, linked to the overlapping variant's eGene (N=2187); 2) a list of all MACS2 DNase peaks in GM1278 overlapping a random selection of 10,000 distal, noncoding background SNPs, each linked to every gene within 1 Mb (N=74,340).

For each set of element-gene pairs, Enformer predictions were made for pairs where the distance between the element and the gene was less than 100kb (due to Enformer's input sequence length of ~200kb). We focused the analysis on Enformer outputs corresponding to CAGE K562 tracks, and we considered the genomic position corresponding to each gene's primary TSS, as chosen by the Enformer model. For each gene, we computed gradients back to the input reference nucleotides for a 200kb region centered at the TSS and took the absolute value to score each single nucleotide. For each element, we calculated the gaussian-weighted mean with sigma = 200 of nucleotide scores in a 2kb window centered on the element. Finally, to normalize for different gene expression levels and compare sites across genes, we divided the element-gene pair scores by the mean absolute value of the nucleotide scores across the entire 200kb region for each gene.

### 13. CTCF loop-constrained Interaction Activity (CIA)

We implemented a "CTCF loop-constrained Interaction Activity" (CIA) model that combines enhancer activity with a measure of 3D contact estimated from CTCF ChIA-PET data. This model has similarities to the ABC model, in that it combines chromatin activity of enhancers and the 3D contact between enhancers and promoters. In the CIA model, enhancer activity ($A_E$) was computed from the geometric mean of normalized DNase-seq and H3K27ac ChIP-seq signal, as defined in the ABC model. Because spatial interactions between enhancers and promoters are largely constrained by CTCF loops, 3D contact was characterized by CTCF Constraint ($CC_{E,P}$), *i.e.* the difference between total PET count of CTCF loops that contain the enhancer-promoter pair ($\Sigma_{\text{CTCF loop i contains E-P}}$ $PET_i$ ) and the total PET count of CTCF loops that cross the enhancer-promoter pair ($\Sigma_{\text{CTCF loop j crosses E-P}}$ $PET_j$). This term was normalized by the sum of PET count of CTCF loops that span the promoter pair ($\Sigma_{\text{CTCF loop i contains P}}$ $PET_i$+1) to allow cross-gene comparison. We filtered out CTCF loops greater than 500kb because most of them were weak and can be noisy. The CIA model implemented in this manuscript is different from Luo *et al.*[8] in some respects, including that, in Luo *et al.*[8], only CTCF loops containing enhancer-promoter pairs are considered.

### 14. Training ENCODE-rE2G logistic regression models

We trained multiple logistic regression models using different combinations of input features, including ENCODE-rE2G and ENCODE-rE2G$^{Extended}$ (**Table S3**). For ENCODE-rE2G, we used 13 features that can be computed from cell-type specific DNase-seq data, cell-type averaged Hi-C data, and genome annotations alone in 352 ENCODE cell types (**Table S2**). For measurements of enhancer activity and 3D contact frequency, we included both first-order and squared values to capture possible non-linear effects of these features (**Table S3**). For ENCODE-rE2G$^{Extended}$, we used an extended set of 47 features available in ENCODE K562 and GM12878 cell lines. (**Table S10**). Our training dataset is a list of 10,411 element-gene pairs tested by CRISPR in K562 cells (see Methods Section 18). We first apply a variance-stabilizing transformation $log(|x| + \epsilon)$, with $\epsilon = 0.01$, to all the features before feeding them to a standard logistic regression model. For the predictions, we use a chromosome-wise cross-validation, meaning that we hold out one chromosome as the test set and train the models on the remaining chromosomes. As our ensemble CRISPR dataset contains the E-G pairs in the chromosomes 1-22 and X, we do the cross-validation 23 times to get the predictions in all the E-G pairs across the genome. To apply the models to new cell types, we generate a similar feature table as described above and apply the model to each candidate element-gene pair using the pre-trained weights learned in K562 cells.

For analyses of the importance of specific features, we also trained variations on the logistic regression model with different subsets of features. For some of these analyses, we defined a "baseline" feature set consisting of the following features (see also **Table S3**): DNase-seq signal at enhancer, DNase-seq signal at promoter, H3K27ac ChIP-seq signal at enhancer, H3K27ac ChIP-seq signal at promoter, 3D contact (ENCODE Hi-C, 5-kb resolution), and distance to TSS.

15. **Collection of previously published enhancer-gene predictions**

We collected the following additional enhancer-gene predictions that have been previously published (**Table S1**). If predictions were provided in hg19, we used UCSC tools LiftOver to lift them over to GRCh38.

*15.1.* *ENCODE2012 E2G.* Distal accessible elements were previously linked to gene promoters by looking at correlation of DNase I hypersensitivity across 125 cell and tissue types from ENCODE[9]. We downloaded these links from: ftp://ftp.ebi.ac.uk/pub/databases/ensembl/encode/integration_data_jan2011/byDataType/openchrom/jan2011/dhs_gene_connectivity/genomewideCorrs_above0.7_promoterPlusMinus500kb_withGeneNames_32celltypeCategories.bed8.gz.

*15.2.* *Anderson 2014.* The transcriptional activity of enhancers and TSSs was previously linked using the FANTOM5 CAGE expression atlas[10]. We downloaded these predictions from: http://enhancer.binf.ku.dk/presets/enhancer_tss_associations.bed.

*15.3.* *Liu 2017*. Gene expression was previously correlated with five active chromatin marks (H3K27ac, H3K9ac, H3K4me1, H3K4me2 and DNase I hypersensitivity) across 56 biosamples, and these correlation links were then used to make predictions for the predicted enhancers (regions with the '7Enh' ChromHMM state) in 127 biosamples from the Roadmap Epigenome Atlas[11]. We downloaded these predictions from www.biolchem.ucla.edu/labs/ernst/roadmaplinking and made predictions using the confidence score.

*15.4.* *Granja 2019 (healthy)*. Single-cell ATAC-seq and RNA-sequencing data in peripheral blood and bone marrow mononuclear cells, CD34+ bone marrow cells and cancer cells from patients with leukaemia were previously analyzed and the ATAC-seq signal in accessible elements was correlated with the expression of nearby genes[12]. We downloaded these predictions from https://github.com/GreenleafLab/MPAL-Single-Cell-2019 and used the correlation in samples from healthy individuals as the quantitative score. Cell-type-specific links were not reported.

*15.5.* *Gao 2020*. EAGLE was previously used to predict enhancer–gene interactions across a number of human tissues and cell lines[13]. The method calculates a score based on six features obtained from the information of enhancers and gene expression: correlation between enhancer activity and gene expression across cell types, gene expression level of target genes, genomic distance between an enhancer and its target gene, enhancer signal, average gene activity in the region between the enhancer and target gene, and enhancer–enhancer correlation. We downloaded enhancer annotations for 104 cell types from http://www.enhanceratlas.org.

*15.6.* *Sheffield 2013*. The DNase I signal and gene expression levels were previously correlated using data from 112 human samples representing 72 cell types to identify regulatory elements and to predict their targets[14]. We downloaded these predictions from http://dnase.genome.duke.edu and used the correlation as the quantitative score. Cell-type-specific links were not reported.

*15.7.* *Cao 2017*. Correlations between gene expression and various enhancer features (for example, DNase1 and H3K4me1) were previously computed across multiple cell types to identify a set of putative enhancers[15]. Then, a sample-specific model is used to predict the enhancer gene connections in a given cell type. We downloaded the lasso-based JEME predictions in all ENCODE+Roadmap cell types from http://yiplab.cse.cuhk.edu.hk/jeme. We used the JEME confidence score as a quantitative score.

*15.8.* *TargetFinder*. A model was previously generated to predict whether nearby enhancer–promoter pairs are located at anchors of Hi-C loops[16].

*15.9.* *ABC (Nasser 2021)*. Full ABC prediction files for 131 cell types[2] were obtained from ftp://ftp.broadinstitute.org/outgoing/lincRNA/ABC/Nasser2021-Full-ABC-Output and lifted over from hg19 to GRCh38 using UCSC tools LiftOver.

*15.10.* *Schraivogel 2020.* A random forest model trained on CRISPRi enhancer perturbation data to predict enhancer-gene regulatory interactions from molecular features[17]. The published model trained on the targeted Perturb-seq (TAP-seq) CRISPRi enhancer screen in the chromosome 8 and 11 regions was taken to make genome-wide predictions in K562 cells. Candidate enhancer elements were defined using the same strategy as for the TAP-seq enhancer screen, *i.e.*, DHS overlapping active enhancer chromatin states from GenoSTAN[18]. Candidate element-gene pairs were created with all protein-coding genes within 300 kb of elements and molecular features for each pair were computed in the same as the original publication: https://github.com/argschwind/TAPseq_manuscript/blob/master/scripts/chromatin_annotated_etps/genome_wide_etps.R

### 16. Generating baseline predictors

We generated two sets of simple baseline predictors in 89 ENCODE4 biosamples (**Table S11)**. Each set utilized a different universe of elements: The first set was based on ENCODE4 DNase-seq pipeline narrowPeak calls (ENCPL848KLD) available through the ENCODE portal (www.encodeproject.org). The second set was based on candidate elements defined in this study using MACS2 peak-calling (see Methods Section 5.1). For each set, all element-gene pairs were created by pairing candidate TSSs with all candidate elements within 1Mb. For each of the 2 sets, 7 types of baseline predictors were generated: 1) Distance from elements to the TSS calculated as the absolute distance between the center of the element and the TSS (“distance to TSS”). 2) Distance from elements to the gene body calculated as the absolute distance between the center of the element and the gene body. If the element overlapped the gene, distance was set as 0. 3) Assign each element to the nearest TSS or gene (“nearest TSS or gene”). 4) Assign each element to the nearest expressed TSS or gene (“nearest expressed TSS or gene”), where “expressed” was approximated using promoter activity in the ABC model as previously described[1]. 5) Assign each element to every TSS within 100kb (“within 100kb of TSS”). 6) Multiply the read-depth normalized DNase-seq or H3K27ac ChIP-seq signals within the element by 1/distance to TSS (“reads by distance to TSS”; a simplified version of Activity x Contact). 7) “reads by distance to TSS” scores normalized to 1 per gene (a simplified version of the ABC score).

Not all candidate elements in the collected CRISPR datasets are part of the baseline predictor element universe and others can have different boundaries. For optimal baseline predictors in the CRISPR benchmarking pipeline, additional versions of the “distance to TSS”, “distance to gene”, “nearest TSS of Gene”, “nearest expressed TSS or gene”, “within 100kb of TSS” were computed based on the universe of all CRISPR element-gene pairs.

### 17. Defining classification thresholds for predictive models

We used performance in the CRISPR benchmarking analysis to define classification thresholds for all generated and collected predictive models. For each predictor, a threshold was chosen that corresponds to 70% recall on the combined CRISPRi data (**Fig 2b, Table S1)**. These thresholds were then applied to create binary predictions of enhancer-gene regulatory connections in all cell types or biosamples to which a given predictor was applied.

### 18. CRISPRi benchmark

We developed a CRISPRi benchmarking pipeline to compare predictive models to perturbation data from previous studies in which CRISPR was used to perturb candidate enhancers and effects on gene expression were measured in K562 cells. We collected and reprocessed 3 previously published CRISPRi enhancer screen datasets from Nasser *et al.*, 2021[2] (itself comprised most of CRISPRi screens from Fulco *et al.*, 2019[1]), Gasperini *et al.*, 2019[19] (Perturb-seq) and Schraivogel *et al.*, 2020[17] (TAP-seq). See **Note S1** for more information on the overall approach.

18.1. *Analyzing CRISPRi Perturb-seq/TAP-seq enhancer screen data.* We have developed an analysis pipeline to identify regulatory enhancer–gene pairs from Perturb-seq experiments. For each perturbed element, differential expression testing between perturbed and unperturbed cells was performed to identify target genes. UMI counts were normalized for a size factor based on total counts per cell, excluding top 10% expressed genes per cell for size factor calculation, and then log-transformed. Cells carrying at least one guide RNA targeting a given element were tested against a specified number of control cells sampled from cells not carrying a perturbation for the element using MAST (v. 1.20.0)[20]. Benjamini-Hochberg false discovery rate was computed to correct for multiple testing for all tests performed across perturbed elements.

In addition, we also implemented a simulation-based power analysis to estimate the statistical power of every tested enhancer-gene pair to detect perturbation effects of specific effect sizes based on the number of perturbed cells and gene expression levels. UMI counts were simulated from negative binomial distributions with mean and dispersion parameters estimated for every gene using DESeq2 (v. 1.34.0)[21]. Mean expression levels across all cells were used as unperturbed expression levels and perturbation effects for each enhancer-gene pair were simulated by injecting a specified decrease in mean expression for perturbed cells. Simulated UMI counts were normalized, and differential expression tests and multiple testing correction were performed using the same approach as for real data. Simulations were repeated 20 times and statistical power for each pair was defined by the proportion of simulations in which the injected perturbation effect was detected as statistically significant.

*18.2.* *Processing of CRISPRi-Perturb-seq data from Gasperini et al., 2019.* Processed data containing UMI counts per cell and gene, and detected guides per cell were downloaded from GEO (accession: GSE120861). We recomputed guide-to-element assignments for our analysis. To do so, genomic coordinates of guideRNA binding sites were inferred using BLAT to align guide sequences to the hg19 genome sequence. Candidate elements were defined by extending the summits of the top 150,000 K562 DNase-seq peaks by 250bp upstream and downstream and merging overlapping regions. Guides were assigned to targeted elements by overlapping guide binding sites with the generated candidate elements. For each targeted element, genes within 2Mb were tested for differential expression between perturbed cells and 5,000 randomly sampled control cells as described above. Power simulations were performed for all candidate element-gene pairs included in the differential expression analysis. Benjamini-Hochberg False Discovery Rate (FDR) was calculated for all tested element-gene pairs and a cutoff of < 5% was applied to identify positive hits for both differential expression and power simulations. The final dataset was compiled by filtering all tested E-G pairs for a distance to TSS between 1kb and 1Mb and any pairs where the enhancer overlapped the gene body of their respective target gene were removed, due to the ability of CRISPRi to repress a gene when targeted to its gene body[1,22]. Negative E-G pairs were further filtered for minimum statistical power of 80% to detect a 15% decrease in target gene expression upon perturbation.

*18.3.* *Processing of CRISPRi-TAP-seq data from Schraivogel et al., 2020.* Raw data was processed as in Schraivogel et al., 2020[17]. For each targeted element, all genes within the same genomic region (chr11 and chr8) were tested for differential expression between perturbed cells and 3,500 randomly sampled control cells with the same distribution across 10x lanes as perturbed cells as previously described[17]. Power simulations were performed for all element-gene pairs included in the differential expression analysis. Benjamini-Hochberg False Discovery Rate (FDR) was calculated for all tested element-gene pairs and a cutoff of < 5% was applied to identify positive hits for both differential expression and power simulations. The final dataset was compiled by filtering all tested E-G pairs for a distance to TSS between 1kb and 1Mb and any pairs where the enhancer overlapped the gene body of their respective target gene were removed. Negative E-G pairs were further filtered for minimum statistical power of 80% to detect a 15% decrease in target gene expression upon perturbation.

18.4. *Processing of CRISPR enhancer perturbation data from Nasser et al.,* 2021[2]. This dataset contains mostly data from CRISPRi FlowFISH DHS tiling experiments[1], with additional E-G pairs added from other published studies as previously described[1,2]. The processed dataset was downloaded from: https://raw.githubusercontent.com/EngreitzLab/ABC-GWAS-Paper/main/comparePredictorsToCRISPRData/comparisonRuns/K562-only/experimentalData/experimentalData.K562-only.txt and reformatted for our benchmarking framework.

18.5. *Combined CRISPR dataset.* We combined the three individual power-filtered datasets into one large, combined dataset. In case a given E-G pair was targeted in more than one dataset, one pair was chosen using the following heuristic: 1) Only positive E-G pairs were retained if there were any. 2) If there were no or more than one positive, one pair was chosen based on the highest statistical power to detect a 25% expression change.

18.6. *CRISPR benchmark.* To benchmark the performance of predictive models, all E-G pairs from predictions were overlapped with the CRISPR E-G pairs using the following algorithm (for each predictive model separately): For a given gene found in both the CRISPR data and predictions (matched by gene symbol) CRISPR elements were overlapped with elements in predictions based on genomic coordinates. If a CRISPR element overlapped a prediction element, the prediction score was assigned to that CRISPR E-G pair. If more than one prediction element overlapped the CRISPR element, the prediction scores were aggregated using a predictor-specific function (*e.g.*, sum or max, see **Table S1**). If no prediction element overlapped a given CRISPR element, this element was considered as “not predicted” by the predictive model and the minimum possible prediction score was assigned to that CRISPR pair. The merged CRISPR and predictions data was then used to compute precision-recall curve metrics (ROCR v. 1.0-11) using the binary experimental outcome whether a CRISPR E-G pair was detected as positive or negative as ground truth.

18.7. *Quantifying model performance and comparing models using bootstrapping.* To quantify and compare model performance, Area Under the Precision-Recall Curve (AUPRC) and precision at 70% recall were computed (**Table S12**). A bootstrapping approach was applied to estimate 95% confidence intervals by re-sampling with replacement (1000 iterations) and computing the range containing 95% of bootstrapped values. The R package boot (v. 1.3-28.1) was used to perform bootstrap sampling. To assess the significance of pairwise comparisons in performance between models, the delta AUPRC or delta precision at 70% recall was computed. The same bootstrap scheme as described above with 10,000 iterations was used to obtain 95% confidence intervals and the approach to compute p-values through inversion of confidence intervals implemented in the boot.pval (v. 0.4.1) R package was applied.

### 19. GWAS benchmark

We developed a "GWAS benchmarking pipeline to assess whether predictive models of enhancer-gene regulatory interactions could (i) identify enhancers enriched for fine-mapped GWAS variants, and (ii) link those enhancers to genes already known to be involved in the GWAS disease/trait, similar to our previous approach[2] (**Table S13**). For this benchmarking pipeline, we analyzed data from the UK Biobank for 94 traits.

19.1. *Fine-mapped GWAS variants.*

We obtained fine-mapping results and summary statistics 94 traits based on an unpublished analysis ( J.C.U., M. Kanai and H.K.F., unpublished data) that analyzed data from the UK Biobank (application 31063; fine-mapping data are available at https://www.finucanelab.org/data). In this analysis, up to 361,194 individuals of white

British ancestry with available phenotypes and variants with INFO > 0.8, minor allele frequency > 0.01%, and Hardy–Weinberg equilibrium P > 1 × 10−10 were included in the GWAS. Covariates for the top 20 principal components, sex, age, age2 , sex × age, sex × age2 and dilution factor, where applicable, were controlled for in the association studies. Quantitative traits were inverse rank transformed and associations were estimated using BOLT-LMM for quantitative traits and SAIGE for binary traits. In-sample dosage linkage disequilibrium was computed using LDStore, and phenotypic variance was computed empirically. Fine-mapping was performed using the sum of single effects (SuSiE) method, allowing for up to ten causal variants in each region. Prior variance and residual variance were estimated using the default options, and single effects (potential 95% credible sets) were pruned using the standard purity filter such that no pair of variants in a credible set could have r 2 > 0.25. Regions were defined for each trait as ±1.5 Mb around the most significantly associated variant (with this window chosen based on the linkage disequilibrium structure in the human population), and overlapping regions were merged. Variants in the MHC region (chr. 6: 25–36 Mb) were excluded as were 95% credible sets containing variants with fewer than 100 minor allele counts. Coding (missense and predicted loss of function) variants were annotated using the variant effect predictor v.85.

For all traits, except where specified, we considered only the 'noncoding credible sets'—that is, those that did not contain any variant in a coding sequence or within 10 bp of a splice site annotated in the RefGene database (downloaded from UCSC Genome Browser on 24 June 2017)

19.2. *Calculating enrichment of GWAS variants in merged enhancer elements.*

We filtered for enhancers where Score > threshold (**Table S1**) for downstream enrichment analysis. For a given trait, we intersected variants with PIP ≥ 10% in noncoding credible sets with enhancers (or other genomic annotations). We merged enhancer regions across cell types to generate a unique set of enhancer regions for each predictor. We then calculated enrichment values by looking at the number of variants with PIP ≥ 10% that overlapped the merged enhancer set over the total number of background variants, defined by common and low frequency variants (Minor allele count >=5) in 1000 Genomes Project, that overlapped enhancers. For calculating enrichment values excluding promoter regions, we performed the same calculations but only considered enhancers that did not overlap promoter regions.

19.3. *Disease Heritability enrichment using Stratified LD score regression.*

Stratified LD score regression (S-LDSC) is a method that assesses the contribution of a genomic annotation to disease and complex trait heritability[23,24]. S-LDSC assumes that the per-SNP heritability or variance of effect size (of standardized genotype on trait) of each SNP is equal to a linear contribution of each annotation. $var(\beta_j) = \sum_j a_{cj}\tau(c)$, where $a_{cj}$ is the value of annotation c for SNP j, and $\tau(c)$ is the contribution of annotation c to per-SNP heritability conditioned on other annotations. S-LDSC estimates the $\tau(c)$ for each annotation using the following equation:

$$E({\chi^2}_j) = N \sum_c l(j,c)\,\tau(c) + 1$$

where $l(j,c) = \sum_k a_{ck}\, {r^2}_{jk}$ is the stratified LD score of SNP j with respect to annotation c and $r_{jk}$ is the genotypic correlation between SNPs j and k computed using data from 1000 Genomes Project (see URLs); N is the GWAS sample size. We assess the informativeness of an annotation c using two metrics. The first metric is enrichment (E), defined as follows (for binary and probabilistic annotations only):

$$E = h^2{}_g(c)/h^2{}_g \times M/\sum_j a_{cj}$$

where $h^2{}_g(c)$ is the heritability explained by the SNPs in annotation c, weighted by the annotation values. The second metric is standardized effect size ($\tau$*) defined as follows:

$$\tau*(c) = \tau(c)\,sd_c\,/\,(h^2{}_g\,/M)$$

where $sd_c$ is the standard error of annotation c, $h^2{}_g$ is the total SNP heritability and M is the total number of SNPs on which this heritability is computed (equal to 5,961,159 in our analyses). $\tau*(c)$ represents the proportionate change in per-SNP heritability associated to a 1 standard deviation increase in the value of the annotation.

*19.4.* *Precision-recall analysis of silver-standard causal genes in GWAS loci.* We evaluated the accuracy of linking noncoding GWAS variants to target genes using a strategy we previously described[25], in which we attempt to link noncoding GWAS signals to nearby genes that carry coding variants independently associated with the same trait. The genes carrying coding variants are very likely relevant to the trait, and are more likely than other genes in the same locus to be the target of the independent noncoding variant[25]. Using fine-mapping for UK Biobank traits as described above, we defined a set of 560 noncoding credible sets (comprised entirely of noncoding variants) corresponding to 31 UKBiobank traits that are located near exactly 1 gene carrying a coding variant, similar to the approach we previously described[25]. First, we used the Variant Effect Predictor[26] to identify protein-truncating variants and damaging missense variants with posterior inclusion probability from fine-mapping (PIP) >= 50%. Second, we examined noncoding credible sets in which exactly 1 gene within 2 Mb carried such a coding variant. Third, we removed duplicated credible sets in which the same noncoding variant was associated with multiple traits (*e.g.*, due to genetic correlation between the traits). We deduplicated such cases by (i) finding 'duplicate' variant-gene pairs where the same noncoding variant had PIP >= 0.05 for more than two traits, and was in the same locus attempting to link to the same gene with a coding variant; and (ii) removing such duplicates, keeping the credible set where the variant had a higher PIP. For the primary benchmarking analysis in **Fig. 2f**, we restricted our analysis to 197 out of the 560 noncoding credible sets corresponding to 11 blood-related traits (Eosinophil count, Lymphocyte count, Monocyte count, Neutrophil count, Platelet count, RBC count, Hemoglobin count, Mean Corpuscular Hemoglobin, Mean Corpuscular Hemoglobin concentration, Mean Corpuscular Volume, All Autoimmune Disease - UKBB). This choice was motivated by the fact that immune cell types had a much more consistent and elaborate representation across different enhancer-gene strategies compared to other cell types. Results for all 560 noncoding credible sets are reported in **Extended Data Fig. 4**.

### 20. eQTL benchmark

We developed an "eQTL benchmarking pipeline" to assess whether predictive models of enhancer-gene regulatory interactions could (i) identify enhancers enriched for fine-mapped eQTL variants, and (ii) link those enhancers to their corresponding eQTL target genes. We analyzed variants from the GTEx V8 resource[27] fine-mapped using the Sum of Single Effects (SuSIE) model[28].

*20.1.* *Fine-mapped eQTL variants.* We obtained the set of GTEx V8 variants fine-mapped using SuSiE[28] by Hilary Finucane and Jacob Ulirsch at the Broad Institute. We considered variants with a posterior inclusion probability greater than 0.5 and that were included in a credible set. We converted the target genes for each variant from their Ensembl ID to HGNC symbol, removing variants without a corresponding HGNC symbol. We mapped the median transcripts per million (TPM), obtained from GTEx (https://storage.googleapis.com/gtex_analysis_v8/rna_seq_data/GTEx_Analysis_2017-06-05_v8_RNASeQCv1.1.9_gene_median_tpm.gct.gz), to each variant depending on the target gene and tissue it was identified in. We included variants linked to genes expressed at levels above 1 TPM from their respective tissues. For each predictive model, GTEx variants and enhancer-gene links were filtered to those linked to genes in their shared gene universe. The GTEx variants and TPM file are available to download on Synapse at the following link: https://www.synapse.org/#!Synapse:syn52264240.

20.2. *Filtering to distal noncoding SNPs.* For eQTL benchmarking analyses, we focused on "distal noncoding SNPs" by filtering out (i) coding sequences, 5′ and 3′ untranslated regions of protein-coding genes, and splice sites (within 10 bp of a intron–exon junction of a protein-coding gene) of protein-coding genes, and (ii) promoters (±250 bp from the gene TSS) of protein-coding genes.

20.3. *Defining tissue - biosample matches for eQTL benchmarking.* For each predictive model that was applied to multiple biosamples, we assigned each GTEx tissue to the closest matching biosample (*e.g.*, GTEx tissue Muscle_Skeletal was paired with ENCODE-rE2G and $ABC^{A=DNase,\ C=Avg.\ ENCODE\ Hi\text{-}C}$ biosample skeletal_muscle_cell_ENCDO094AAA_ENCFF724SPV). The set of 12 tissues represented in all predictive models was used for the multiple biosample benchmarking analysis (**Extended Data Fig. 3c,d**), and the biosamples matching each of the 12 tissues for each predictive model are listed in **Table S14**. To generate enrichment-recall curves (**Fig. 2d**), the GTEx tissue "Cells_EBV-transformed lymphocytes" was paired with enhancer-gene predictions in a particular lymphoblastoid cell line, GM12878.

20.4. *Calculating enrichment of eQTLs in predicted enhancers.* Here we defined enrichment as (fraction of eQTL variants with PIP >= 50% that overlap predicted enhancers) / (fraction of all 1000G SNPs that overlap predicted enhancers) (see **Fig. 2d** and **Extended Data Fig. 3b,c,d**). To calculate this enrichment, we obtained a list of approximately 10 million SNPs from the 1000 Genomes Project from the Price Group (https://alkesgroup.broadinstitute.org/LDSCORE/baseline_v1.1_hg38_annots/), then filtered them to the distal noncoding regions of the genome as described above. For each predictive method, we further filtered the GTEx variants to those with an eGene in the gene universe considered by that predictive method; in absence of a provided gene universe file, our curated GENCODE gene universe was used. This default gene universe was curated from the GENCODE v29 gene annotations by selecting "basic"-tagged protein-coding transcripts (referring to a subset of representative transcripts for each gene) with levels 1 or 2 confidence (verified or manually annotated loci), The code used to generate this file is available here: https://github.com/EngreitzLab/ABCGeneReference. We then calculated the number of variants from each GTEx tissue. Next, we thresholded the predictions for each method based on the threshold that yielded 70% recall from CRISPRi benchmarking, and intersected the predictions from each biosample with the GTEx variants from each tissue, recording the number of variants at each intersection. We also calculated the number of distal noncoding SNPs intersecting thresholded predictions in each biosample. The enrichment for each prediction biosample, B and GTEx tissue, T intersection was then calculated as [number of variants in T overlapping predicted enhancers in B / total number of variants in T]/[number of distal non-coding 1000G SNPs overlapping B / total number of 1000G SNPs]. We used a modified enrichment calculation for Enformer: Because the predicted enhancers for Enformer were generated based on a control set of DNase peaks overlapping a random set of 10,000 1000G SNPs, the enrichment calculation was modified for by defining the background set of distal, noncoding variants as this list of 10,000 variants.

20.5. *Calculating recall of predictive models at linking eQTL variants to target eGenes.* To calculate the recall of prediction methods in linking an eQTL variant to its eGene, GTEx tissues were matched with biosamples from each prediction method to define corresponding pairs (**Table S14**). For each pair, we calculated the 1) total recall, defined as the fraction of eQTL variants overlapping predicted enhancers (**Extended Data Fig. 3c,d**) 2) recall accounting for linking, defined as the fraction of eQTL variants overlapping predicted enhancers that are linked to the variant's eGene (**Fig. 2d**, **Extended Data Fig. 3b**); and 3) the fraction of variants overlapping a predicted enhancer linked to the variant's eGene, given that the variant overlaps a predicted enhancer.

20.6. *Choosing threshold values for enrichment-recall curves.* We calculated a range of predictor score values to construct enrichment-recall curves (**Fig. 2d**, **Extended Data Fig. 3b**) by defining 50 to 100 evenly-spaced steps across the recall for each predictive model. We took the unique set of variants overlapping predicted enhancers where the variant's eGene was linked to the enhancer's target gene, then calculated the score that yielded each interval of the total possible recall. Enrichment and recall were evaluated at each of the threshold values to generate enrichment-recall curves (**Tables S16, S17**).

### 21. Additional analyses: Assaying enhancer activity

To build models for different enhancer activity measurements, element-gene pairs based on candidate elements within 5Mb of candidate promoters were used as the universe of enhancers and genes. Data for 513 1D chromatin assays (**Table S17**) in K562 were downloaded from ENCODE portal as "read-depth normalized signal" (DNase-seq) or "fold change over control" (ATAC-seq, ChIP-seq) bigWig files. Enhancer activities for all candidate elements were calculated for each assay separately using the ABC `run.neighborhoods.py` script, where the different assays were used in place of DNase-seq. The activities for each element and assay were measured by taking the normalized read counts for a given assay. ABC scores for all element-gene pairs and activity measurements were then computed by multiplying activity with the Hi-C contact frequency of the candidate elements and promoters as implemented by ABC.

### 22. Additional analyses: Linking variants to genes analyses

To generate the ENCODE-rE2G + PoPS prediction, we intersected the top two genes with the strongest ENCODE-rE2G prediction score for a peak overlapping a common or low-frequency variant from the 1000 Genomes Project with top two genes in a 1Mb locus around the variant that have the highest Polygenic Priority Score (PoPS)[25]. The PoPS sore prioritizes genes for a disease based on their functional similarity to genes implicated by disease GWAS. PoPS trains a model to predict MAGMA (v1.08)[29] gene scores using 57,543 gene-level functional features based on gene expression data (bulk and single-cell RNA-seq), biological pathways, and PPI networks. PoPS uses a leave-one-chromosome-out framework, such that the PoPS score of a gene on chromosome x is not informed by GWAS data or the ENCODE-rE2G linking data on chromosome x. The ENCODE-rE2G + PoPS score, unlike ENCODE-rE2G, is therefore a disease-specific score. For **Fig. 4b**, we assessed the enrichment of the number of disease-related fine-mapped variants (PIP > 0.1) in ENCODE-rE2G + PoPS implicated variants, in comparison to all annotated variants, and also compared precision and recall against silver standard causal genes in the GWAS loci (see GWAS benchmark section above). For **Fig. 4c**, ENCODE-rE2G + PoPS was computed for each of 352 cell-types and 76 diseases and traits using top two ENCODE-rE2G predicted genes at the locus *for each cell-type/biosample*, and intersecting that with top two PoPS prioritized genes.

### 23. Combinatorial enhancer perturbations at *MYC* locus

23.1. *MYC enhancer paired-sgRNA (pgMYC) library design.* We conducted a CRISPRi-FlowFISH screen to perturb all pairs of 7 enhancers that we previously found to regulate *MYC* in K562 cells[22] (**Fig. 6d-g**). A lentiviral paired sgRNA vector was used to clone the MYC enhancer paired-sgRNA library, such that each vector expresses two sgRNAs driven by either the hU6 or mU6 promoter. Seven previously characterized MYC enhancers (e1-7), the MYC TSS, two constituent cCREs adjacent to e6 and e7, and a nearby negative control DHS peak (NS1) were selected as target cCREs in this experiment[22].

We designed a lentiviral guideRNA library containing 10,080 pairs of sgRNAs (ENCFF118BUP). For significant CREs characterized in the previous MYC enhancer screen[22], we first selected two sgRNAs that had been used in low-throughput enhancer CRISPRi and H3K27ac ChIP-seq assays, then selected six other sgRNAs with the greatest average effect across the previous MYC enhancer screen's replicates. For NS1, we selected the two sgRNAs that had been used in low-throughput enhancer CRISPRi and H3K27ac ChIP-seq assays, then selected other sgRNAs by first sorting all GuideScan sgRNAs in the region with CFD specificity scores greater than or equal to 0.2 and efficiency scores greater than or equal to 50, then selecting six sgRNAs such that their rank-ordered genomic positions are evenly spaced apart (six sgRNAs are selected in an every $n^{th}$ fashion after ordering them according to their PAM site's genomic position). For the two constituent cCREs adjacent to e6 and e7, we selected eight sgRNAs using only the GuideScan filters and every $n^{th}$ sgRNA approach, as above. Finally, we included 12 sgRNAs targeting genomic "Safe" harbor regions that lack epigenetic annotations associated with active regulatory elements[30].

These 100 targeting sgRNAs were paired in an all-by-all fashion, such that each sgRNA is paired with every sgRNA (including itself) in both orientations (sgRNA1-sgRNA2 and sgRNA2-sgRNA1). Since recombination between sgRNAs can occur at some frequency due to either lentiviral recombination or PCR-based recombination, we included 80 additional pgRNA vectors in which 5 new "Safe"-targeting sgRNAs that were not among the other 12 "Safe"-targeting sgRNAs were paired with all eight NS1 negative control sgRNAs in both orientations. This allows us to empirically calibrate the recombination rate, by detecting the frequency with which the 5 new "Safe"-targeting sgRNAs are paired with any of the 92 non-NS1-targeting sgRNAs. The entire library consists of 10,080 paired sgRNA vectors.

23.2. *MYC enhancer paired-sgRNA screen.* The pgMYC lentiviral library was synthesized first as an oligo pool from Twist Bioscience, then cloned into a paired sgRNA vector that constitutively expresses the selectable marker Puromycin. Each oligo encodes a pair of sgRNAs, which is first cloned into a vector with a single hU6 promoter, and Puromycin, which is constitutively expressed under the Ef1a promoter. In a second cloning step, a cut site between the pair of sgRNAs linearizes the vector, into which we insert the sgRNA stem loop sequence for the upstream sgRNA, and the mU6 promoter for the downstream sgRNA. The lentiviral plasmid library was transfected into HEK293T cells to generate lentivirus, which was then transduced into two separate biological replicates of 55 million K562 doxycycline-inducible dCas9-KRAB cells each at an MOI of 0.1 (>500X coverage per replicate). 48 hours after transduction, cells were selected for 96 hours with 1.0 ug/mL Puromycin, until a non-transduced pool of the same cells was comprised of over 99% dead cells. Cells were recovered in Puromycin-free media and cultured in the absence of doxycycline for an additional week to grow to scale. Throughout their culture, the K562 cells are split back each day to a maximum of 500,000 cells/mL. 100 million cells per replicate were induced with doxycycline at 1 ug/mL for 48 hours prior to harvesting them for FlowFISH. Samples from each biological replicate were split into seven technical staining replicates of 5 million cells, and processed for FlowFISH using probesets for *MYC* and *RPL13A* as previously described[1] (**Extended Data Fig. 7**).

23.3. *Sorting and cytometric analysis.* We diluted the stained cells in PBS with 0.5% BSA to a concentration of 2 × 107 ml–1 and filtered using 35-μm filter tubes (Falcon, no. 352235). We sorted 5 million cells for each replicate into six bins based on the fluorescence intensity of target genes, using the Stanford FACS Core's Influx Sorter (BD Influx, Special Order). Prior to sorting, measurements of the fold-change (FC) for each probeset between a control unprobed sample and each replicate were made to ensure that each of the samples were adequately probed and amplified to a FC >2. To control for differences in staining efficiency for each cell, we normalized the fluorescence associated with the gene of interest to that of the control gene (RPL13a) as previously described[1]. We set the gates for each bin on the compensated signal to capture 10% of the cells according to the percentiles (1) 0–10%, (2) 10–20%, (3) 35–45%, (4) 55–65%, (5) 80–90% and (6) 90–100% (**Extended Data Fig. 7**).

*23.4.* *DNA isolation.* We collected the sorted cells by centrifugation at 800*g* for 5 min, resuspended them in 100 µl of lysis buffer (50 mM Tris-HCl, pH 8.1, 10 mM EDTA, 1% SDS) and incubated them at 65 °C for 10 min for reverse cross-linking. We then added 10 µl Proteinase K (NEB, no. P8107S), mixed well and incubated the mixture at 37 °C for 2 h followed by incubation at 95 °C for 20 min. We extracted genomic DNA using AMPure XP (SPRI) beads (Beckman Coulter, no. A63882) at 0.7 beads:cell lysate concentration.

*23.5.* *MYC dual guide screen library preparation and sequencing.* The integrated sgRNAs were amplified using custom primers compatible with Illumina sequencing: AATGATACGGCGACCACCGAGATCTACAC-NNNNNNNNNN-CGTCCGCGGGCTTACCGTAACTTGAAAGTATTTCGATTTCTTGGC (where NNNNNNNNNN denotes i5 10bp 96-well barcode) and CAAGCAGAAGACGGCATACGAGATAGACAGCAGTCCCGTGTTCCGGTTCATTCTATCA-NNNNNN-GGATCCCCTAGGAAAAAAAGCACCG (where NNNNNN denotes i7 6bp replicate barcode)**.** PCR conditions: initial denaturation 98 °C for 30 s, 25 cycles of 98 °C for 30 s, 60 °C for 30 s and 72 °C for 2.5 min; and final amplification 72 °C for 5 min. The samples were pooled and purified with AMPure XP beads at a 1:1 beads:PCR product ratio. Sequencing was performed using an Illumina NextSeq with custom primers: Read 1, 25 bases - to read the sequence of Guide 1 (custom primer: CGATTTCTTGGCTTTATATATCTTGTGGAAAGGACGAAACACCG); Read 2, 20 bases - to read 6bp i7 index (custom primer: AGACAGCAGTCCCGTGTTCCGGTTCATTCTATCA); Index 1, 20 bases - to read the sequence of Guide 2 (custom primer: GTAATTGTGTGTTTTGAGACTATAAGTATCCCTTGGAGAACCACCTTGTTGG); Index 2, 10 bases - to read i5 10bp sample barcode (custom primer: ATCGAAATACTTTCAAGTTACGGTAAGCCCGCGGACG).

*23.6.* *CRISPRi-FlowFISH data processing.* For each sorted bin per sample, reads were aligned to an index of our guide spacers using Bowtie 2[31]. The abundance of each sgRNA pair was quantified using Samtools[32]. These count data were subjected to our previously described pipeline, in which we use the frequency of sgRNA counts in each of the sorted bins to estimate the mean fluorescence value for each sgRNA[1]. Briefly, the mean log10 fluorescent intensity of each sgRNA pair was computed using maximum likelihood estimation fitted to the normalized counts of each sgRNA in the fluorescently sorted bins. Effect sizes for each sgRNA pair were computed by dividing the means of each sgRNA pair by the means of the negative control sgRNAs. We then aggregated information across multiple gRNA pairs to estimate the effect size for perturbing each element or combination of elements. Followed our previous procedure, we adjust the FlowFISH effect size estimates to account for constant background signal that appears to result from nonspecific binding of the probeset[1]; here, we scaled the data using the observed qPCR estimated effects at E2 from our prior study[1] and comparing them to the E2 effects we saw in this CRISPRi experiment (1.57-fold correction). To calculate effect sizes for each element individually, we analyzed guide pairs in which enhancer targeting sgRNAs are paired to negative control sgRNAs. Leveraging this, we computed the effect sizes when targeting a single element and then subsetted our data by the 2 sgRNAs that had the greatest effects when configured with a negative control for every element to refine our dataset to guides of the highest quality. Further quality control was implemented by only including samples where at least 200 cells were observed for a given guide pair across a given sorted sample. From this subsetted data, we re-computed the effect sizes for every guide pair now including the guide pairs that target multiple elements. Each of the sgRNAs' mean effect sizes were computed for every technical replicate and aggregated by target. We performed a Student's two sided, one sample T-test for significance testing where each of the technical replicates' estimation of the effect of perturbing a given element pair is considered one data point.

*Modeling Enhancer-Gene Regulation MYC Data.* We compared the observed data to two null models: one where enhancers combine independently and additively in linear expression space ("additive model"), and one where they combine independently and multiplicatively in linear expression space ("multiplicative model", or additive in log space). The individual effects of each element were estimated as the average of all pairs that include one sgRNA targeting a given element and one negative control sgRNA. To compute the expected effect of perturbing a pair of elements under an additive model, we combinatorially summed the mean effects (% decrease in expression) of each of the two individual elements, and computed 95% confidence intervals by first computing the standard error of the mean of the effect of each element across replicate FlowFISH experiments, and then combining these estimates across the two individual elements using the variance sum law.

We repeat this same analysis for a multiplicative model where the null is the product of the independent effects of each element (Effect of Enhancer 1 (E1) * Effect of Enhancer 2 (E2) = Expected dual perturbation effect).

23.7. *MYC dual guide screen statistical analysis.* To build an estimate of the effect that each of our perturbations have on gene expression, we took the average effect across all relevant guide pairs for each element pair perturbation within each FlowFISH technical replicate.The mean of these FlowFISH replicate estimates represents our estimated effect for a given element pair perturbation. We then combinatorially added individual element perturbation effects to build a matrix of expected effect sizes under an additive model. To determine whether there are any interaction effects between enhancer pairs, we performed an F-test using a linear model of enhancer interactions:

Effect on expression = Effect of Enhancer 1 (E1) + Effect of Enhancer 2 (E2) + E1*E2

where the expectation of the paired mean is computed by adding the independent effects at each enhancer. We observe significant interaction effects for each enhancer pair, suggesting that each enhancer interacts in a super-additive fashion.

### 24. Additional analyses: Enhancer synergy analyses

24.1. *Analysis of CRISPRi tiling experiments.* We analyzed comprehensive tiling CRISPRi experiments to determine whether the sum of the effects of enhancers for a given gene would add up to more than 100% (**Fig. 6a**). To do so, we considered a total of 20 CRISPRi experiments where all candidate elements (DNase peaks) around a given gene were perturbed and the effect on gene expression was measured: PPIF in 6 cell types or conditions[2], HBE1 in K562[33], and 13 other genes in K562[1,22], all of which were previously aggregated[2] and available from https://github.com/engreitzlab/abc-gwas-paper.

Enhancers were filtered to those with a negative and significant ($P_{adj}$ < 0.05) effect on gene expressions and to those not overlapping promoters, except for two regions overlapping the lncRNA PVT1 promoter that were previously shown to act as enhancers to regulate MYC[22].

24.2. *Chromatin immunoprecipitation for H3K27ac following CRISPRi perturbations to MYC enhancers.* We conducted experiments to measure the effects of enhancer perturbations on H3K27ac signals at nearby enhancers (**Fig. 6g, Extended Data Fig. 8 a,b**). We selected all 7 enhancers that we previously identified to regulate *MYC* in K562 cells [e1-e7 from [22]], as well as one additional element in the same locus that did not regulate *MYC* (NS1 from [22]). We perturbed these enhancers with CRISPRi and conducted ChIP-seq for H3K27ac as previously described[34]. Briefly, we stably delivered individual gRNAs by lentivirus into CRISPRi K562 cells (1 gRNAs per element, as well as 4 negative control gRNAs, see [22]), activated CRISPRi with doxycycline for 48 hours, and then harvested cells for formaldehyde-crosslinked H3K27ac ChIP-seq. We performed 2 replicate ChIP-seq experiments on each of 2 biological replicates per guideRNA (3-4 ChIP-seq experiments per guideRNA, yielding 3-4 experiments per element and 16 experiments including negative control guideRNAs). We aligned and processed these data in genome build hg19 as previously described[2]. We computed a fold-change in H3K27ac reads per million at each ABC candidate enhancer in K562 as defined in[2] by comparing the 3-4 experiments per element to the 16 negative control experiments.

24.3. *Analysis of CRISPRi-H3K27ac ChIP-seq from Fuentes et al.* We analyzed data from a previous study in which CRISPRi was directed simultaneously to hundreds of long terminal repeats (LTRs) in the NCCIT cell line using the CARGO system, followed by H3K27ac ChIP-seq to detect changes in enhancer activity[35] (**Fig. 6g**). We used this data to assess how perturbations to enhancers affect H3K27ac ChIP-seq signal at other nearby enhancers. Data from Fuentes et al. was processed similarly to our previous analysis[1]. We first identified enhancer elements by calling peaks from ATAC-Seq data generated in the NCCIT cell line. Each peak was resized to be approximately 500bp centered on the peak summit. For each of these elements we computed the log fold change of H3K27ac ChIP-Seq signal in the wild-type and CRISPRi conditions. We next identified regions potentially targeted by CRISPRi as previously described[1]. Briefly, we first identified LTR regions as the union of DNA elements annotated as either LTR5HS, LTR5A or LTR5B repeats in version 4.0.5 of the RepeatMaster database or as peaks called from dCas9 ChIP-Seq performed by Fuentes et al. This resulted in 1427 candidate LTR regions. Given that these LTR regions may have high sequence similarity, we only considered LTR regions that were sufficiently uniquely mappable. As described in Fulco et al. 2019[1], we implemented a simulation approach to assess the mappability of these regions and only considered LTR regions which had >95% uniquely mapping simulated reads. We also only considered LTR regions which were at least 250kb away from any other LTR region. This resulted in 625 LTR regions. We analyzed the fold change in H3K27ac signal for each ATAC-Seq peak within 2mb of an LTR for a total of 136713 peak-LTR pairs.

24.4. *Analysis of H3K27ac chromatin-QTLs from Delaneau et al. (2019)*. We analyzed H3K27ac chromatin-QTLs[36] to determine how a variant that affects H3K27ac signal in an overlapping peak can affect H3K27ac signal at other nearby peaks (**Extended Data Fig. 8c**). We downloaded data from https://zenodo.org/record/2572871#.Y_P5G-zMK3I. We considered chromatin-QTLs affecting both the H3K27ac peak the variant is located in, and at least one other H3K27ac peak, where the self-peak is less than 5kb in length and where the distance between the self-peak and other peak is greater than 1kb. The relative effect size was defined as negative beta for the nearby peak divided by beta of the self-peak.

24.5. *Analysis of H3K27ac ChIP-seq from Huang et al. (2018).* We analyzed data from a previous study in which individual enhancers were genetically deleted with CRISPR followed by ChIP-seq data of H3K27ac[37] (**Fig. 6h**). Data in bigwig format was downloaded from NCBI GEO (GSE107726) and H3K27ac RPMs for control and perturbed regions were computed directly from the bigwig files. RPMs were then averaged across two replicates. We analyzed perturbation-element pairs separated by greater than 1kb, and calculated the fold-change in H3K27ac RPM at the element after perturbation as log2 (H3K27ac RPM after knockout) / (H3K27ac RPM control).

### 25. Additional analyses: Enhancer-promoter correlation

We conducted an analysis to determine whether eQTL variant-gene pairs were enriched for having high GLS Coefficients (DNase-DNase correlations, see **Methods Section 7**) compared to control variant-gene pairs (**Fig. S3d**). To compare GLS coefficients with eQTL data, we downloaded the GTEx lymphoblastoid cell line eQTL data from the EMBL/EBI eQTL Catalogue[38]. We set a threshold on the GM12878 DNase-seq data of 1.5 to consider a particular ENCODE cCRE active in GM12878, and kept the eQTL variants that lie within these cCREs but do not overlap the promoters and exons of protein-coding genes. We considered eQTL variants with PIP > 0.05 (N=1,083 eQTLs). To compare eQTLs to a control set of variants, we selected non-significant eQTLs ($P > 0.5$) that connect active enhancers and nearby genes, and sampled 8,000 instances that follow the same distance to target gene distribution as significant eQTLs.

We also analyzed whether genes or elements with particular patterns of activity across biosamples behaved differently with respect to whether their GLS coefficients predict the results of CRISPR perturbations in K562 cells (**Fig. S3f-k**). We defined a gene as “broadly expressed” if its DNase-seq signal value was above 2 in all 96 biosamples, and “variably expressed” otherwise. We defined a candidate element as “broadly active” if it showed DNase-seq signal above 1.5 in at least 3 of 96 biosamples, and “specifically active” otherwise. We selected these thresholds to obtain the similar amounts of CRISPRi positive enhancer-promoter pairs across different combinations of promoter expression and enhancer activities.

### 26. Data visualization

Box plots are defined as follows: the middle line corresponds to the median; lower and upper hinges correspond to first and third quartiles; the upper whisker extends from the hinge to the largest value no further than 1.5 × IQR from the hinge (where IQR is the interquartile range, or distance between the first and third quartiles); and the lower whisker extends from the hinge to the smallest value, at most 1.5 × IQR of the hinge. Data beyond the end of the whiskers are outlying points and are plotted individually[39], unless box plots are shown on top of all data points or data distribution.

### References methods


1. Fulco, C. P. *et al.* Activity-by-contact model of enhancer-promoter regulation from thousands of CRISPR perturbations. *Nat. Genet.* **51**, 1664–1669 (2019).
2. Nasser, J. *et al.* Genome-wide enhancer maps link risk variants to disease genes. *Nature* (2021) doi:10.1038/s41586-021-03446-x.
3. Bergman, D. T. *et al.* Compatibility rules of human enhancer and promoter sequences. *Nature* **607**, 176–184 (2022).
4. Boix, C. A., James, B. T., Park, Y. P., Meuleman, W. & Kellis, M. Regulatory genomic circuitry of human disease loci by integrative epigenomics. *Nature* **590**, 300–307 (2021).
5. Nurtdinov, R. & Guigó, R. EPIraction. *In preparation*.
6. Karbalayghareh, A., Sahin, M. & Leslie, C. S. Chromatin interaction–aware gene regulatory modeling with graph attention networks. *Genome Res.* **32**, 930–944 (2022).
7. Sahin, M. *et al.* HiC-DC+ enables systematic 3D interaction calls and differential analysis for Hi-C and HiChIP. *Nat. Commun.* **12**, 3366 (2021).
8. Luo, R. *et al.* Dynamic network-guided CRISPRi screen reveals CTCF loop-constrained nonlinear enhancer-gene regulatory activity in cell state transitions. *bioRxiv* (2023) doi:10.1101/2023.03.07.531569.
9. Thurman, R. E. *et al.* The accessible chromatin landscape of the human genome. *Nature* **489**, 75–82 (2012).
10. Andersson, R. *et al.* An atlas of active enhancers across human cell types and tissues. *Nature* **507**, 455–461 (2014).
11. Liu, Y., Sarkar, A., Kheradpour, P., Ernst, J. & Kellis, M. Evidence of reduced recombination rate in human regulatory domains. *Genome Biol.* **18**, 193 (2017).
12. Granja, J. M. *et al.* Single-cell multiomic analysis identifies regulatory programs in mixed-phenotype acute leukemia. *Nat. Biotechnol.* **37**, 1458–1465 (2019).
13. Gao, T. & Qian, J. EnhancerAtlas 2.0: an updated resource with enhancer annotation in 586 tissue/cell types across nine species. *Nucleic Acids Res.* **48**, D58–D64 (2020).
14. Sheffield, N. C. *et al.* Patterns of regulatory activity across diverse human cell types predict tissue identity, transcription factor binding, and long-range interactions. *Genome Res.* **23**, 777–788 (2013).
15. Cao, Q. *et al.* Reconstruction of enhancer–target networks in 935 samples of human primary cells, tissues and cell lines. *Nat. Genet.* **49**, 1428–1436 (2017).
16. Whalen, S., Truty, R. M. & Pollard, K. S. Enhancer–promoter interactions are encoded by complex genomic signatures on looping chromatin. *Nat. Genet.* **48**, 488–496 (2016).
17. Schraivogel, D. *et al.* Targeted Perturb-seq enables genome-scale genetic screens in single cells. *Nat. Methods* **17**, 629–635 (2020).

36. Delaneau, O. *et al.* Chromatin three-dimensional interactions mediate genetic effects on gene expression. *Science* **364**, (2019).
37. Huang, J. *et al.* Dissecting super-enhancer hierarchy based on chromatin interactions. *Nat. Commun.* **9**, 943 (2018).
38. Kerimov, N. *et al.* A compendium of uniformly processed human gene expression and splicing quantitative trait loci. *Nat. Genet.* **53**, 1290–1299 (2021).
39. Wickham, H. *ggplot2: Elegant Graphics for Data Analysis*. (Springer New York, 2009).
